# Multi-domain analysis of the fecal microbiome of dairy heifers, beef heifers, bred beef heifers, and adult beef cows in the central United States using paired 16S rRNA and 18S rRNA metabarcoding

**DOI:** 10.64898/2026.10.08.757609

**Authors:** Dana Palmer, Pamela R. F. Adkins, Paige Isensee, Rebecca A. Dorfmeyer, Elaine Su, Giedre Turner, Daniel J. Davis, Nathan J. Bivens, Zachary L. McAdams, Aaron C. Ericsson

**Affiliations:** University of Missouri College of Veterinary Medicine, Columbia, MO 65211, USA; Veterinary Medicine and Surgery, College of Veterinary Medicine, University of Missouri, Columbia, MO 65211, USA; University of Missouri Metagenomics Center, Columbia, MO 65201, USA; University of Missouri Animal Modeling Core, Columbia, MO 65201, USA; University of Missouri Veterinary Medical Diagnostic Laboratory, Columbia, MO 65211, USA; Pathobiology and Integrative Biomedical Sciences, College of Veterinary Medicine, University of Missouri, Columbia, MO 65201, USA; University of Missouri, Genomics Technology Core, Bond Life Sciences Center, Columbia, MO 65211

**Keywords:** 18S rRNA, 16S rRNA, eukaryotic, prokaryotic, microbiome, metabarcoding, bovine, cattle

## Abstract

The growth and performance of cattle is dependent on efficient feed conversion, facilitated by the gut microbiome, including both prokaryotic and eukaryotic organisms. Extrinsic factors such as management practices and intrinsic factors like age and genetics can shape these communities. To characterize the relative effects of such factors on the prokaryotic and eukaryotic microbiomes, fresh fecal samples from dairy heifers (*n* = 8), beef heifers (*n* = 8), bred beef heifers (*n* = 8), and adult beef cows (*n* = 8) were subjected to paired 16S rRNA (V4) and a novel 18S rRNA (V4-V5) metabarcoding assay, and 18S rRNA amplicons were annotated using two databases in parallel.

Taxonomic resolution of 18S rRNA amplicons was excellent, with SILVA and EUKARYOME databases each providing better resolution of different groups of organisms. While no differences in eukaryotic richness or alpha diversity were detected, beta diversity differed in all pairwise comparisons except between beef heifers and bred beef heifers. Prokaryotic richness was lower in adult beef cows than the other groups, and beta diversity differed in all pairwise comparisons. Adult beef cows were the most distinct, characterized by replacement of fungal organisms with protozoa and loss of thermophilic bacteria in the eukaryotic and prokaryotic compartments, respectively. Among numerous differentially abundant features, dairy heifers harbored greater abundance of *Cooperia oncophora* while beef heifers contained greater abundance of *Buxtonella sulcata* and *Mucor*. Correlation network analysis revealed multiple significant positive and negative associations between bacterial, fungal, and protozoal features during maturation of the fecal microbiome.

Collectively, these data highlight the richness and diversity of both the prokaryotic and eukaryotic compartments of the bovine fecal microbiome, and the influence of management practices and age on both communities. From a technical standpoint, these data demonstrate the ability to resolve fungal, protozoal, and metazoan organisms, including potential parasites and pathogens, with a single metabarcoding assay. They also highlight the nuanced differences between databases and the value of parallel analyses using more than one database.

## Introduction

The gastrointestinal tract (GIT) of livestock species harbors a rich community of prokaryotic and eukaryotic microbes that are essential for optimal feed conversion, growth, and disease resistance. In healthy ruminants, the same dominant bacteria and archaea are found in the rumen regardless of geography and genotype, while the contribution of protozoa is more variable^1^. The prokaryotic fecal microbiome of ruminants is shaped by dietary content and genotype^2–4^, while factors shaping the fecal eukaryotic microbiome are less clearly defined for several reasons related to the traditional methods of detection. Historically, gastrointestinal (GI) parasites were detected or enumerated via direct identification of nematode eggs or protozoal cysts or oocysts in fecal material. This approach has numerous limitations including poor sensitivity and specificity, and poor correlation with true pathogen burden. Alternatively, molecular approaches offer excellent sensitivity and specificity via detection of organismal DNA.

Prokaryotic and eukaryotic communities can be characterized using shotgun metagenomics, wherein all nucleic acid is fragmented and sequenced, or metabarcoding, wherein a variable (i.e., barcode) region of a gene is amplified via PCR and then sequenced. The latter option is considerably less expensive and time-intensive but requires a variable (V) region of a target gene capable of resolving taxonomies of interest, flanked by universally conserved regions to serve as primer targets. Metabarcoding of prokaryotic and eukaryotic DNA often relies on the genes encoding the 16S rRNA and 18S rRNA small ribosomal subunits, respectively. While the 16S rRNA gene is sufficiently variable across individual V regions to provide good taxonomic resolution across the majority of prokaryotes, different individual V regions of the 18S rRNA gene are better suited to resolve certain clades of eukaryotic organisms^5^.

Long-read (LR) sequencing platforms (e.g., Oxford Nanopore Technologies, Pacific Biosciences) allow full-length metabarcoding of the 18S rRNA gene^6^. However, short-read (SR) platforms (e.g., Illumina) are more widely available and account for an estimated 95-98% of total global sequence data. The chemistries used in SR sequencing platforms have been limited to 300 cycles, meaning each strand of DNA can be sequenced from either end (forward and reverse reads) for a total of 300 base pairs (bp). An overlapping sequence region of 50 bp needed to assemble the complete sequence places an upper limit on total read lengths of 550 bp. Thus, 18S rRNA metabarcoding using SR platforms has been restricted to either a single variable (V) region of the gene, or multiple contiguous (but smaller) V regions^7,8^. The current study takes advantage of new 2×500 paired-end (PE) read sequencing chemistry and an 18S rRNA primer pair spanning the two largest contiguous variable regions (V4 and V5) and yielding roughly 650 bp products. In terms of assay development, our goal was to assess the taxonomic resolution of eukaryotic DNA with this platform, and the effect of database selection during assignment of taxonomies.

Studies applying metabarcoding platforms to the microbiome of cattle have focused primarily on the rumen rather than feces, and are often restricted to a single age or breed of cattle. Dairy and beef cattle are typically exposed to different management practices, each carrying risks associated with exposure to different eukaryotic organisms. Moreover, growth and gestation place substantial physiological stress on heifers. In terms of biology, the objectives of the present study were to identify differences in the fecal prokaryotic and eukaryotic fecal microbiomes of beef and dairy heifers, to identify age-associated differences in the fecal prokaryotic and eukaryotic microbiomes of beef cattle, to assess the ability of the 18S rRNA metabarcoding assay to detect intestinal parasites and pathogens, and to identify significant positive and negative relationships among an integrated multi-domain dataset.

## Methods

### Animals and study design

The present research comprised two overlapping cross-sectional study designs comparing beef cattle of three different ages and age-matched beef and dairy heifers. Dairy cattle were Holstein heifers, ranging in age from 136 days (4.5 months) to 180 days (6 months). Dairy heifers were housed on a small pasture lot with access to three-sided shelters. Heifers received balanced nutrition through total mixed rations (TMR) and had *ad libitum* access to well-water through group waterers.

Beef cattle were mixed breed, with three age groups including heifers, ranging in age from 159 days (5.3 months) to 184 days (6.1 months); bred heifers, ranging in age from 16.6 to 18 months; and adult cows, ranging from 6 to 13 years old. Bred heifers and adult beef cows were bred between 20 and 80 days prior to sample collection. Beef cattle of all ages were maintained on pasture in Boone County, central Missouri, with access to shelter and calf huts for the younger group of beef heifers. Beef cattle had access to vitamin and mineral supplements and potable water from pond water and automatic waterers. Details related to individual animals are provided in **Table S1**.

### Sample collection

Fecal samples were collected manually *per rectum*, placed in a sterile specimen container, and placed on ice during transport to the lab. Dairy heifers were manually restrained in pasture for sample collection. Beef cattle of all ages were run through a chute and briefly restrained in a head-catch for sample collection.

### DNA extraction

DNA was extracted at the University of Missouri (MU) Metagenomics Center using QIAamp PowerFecal Pro DNA extraction kits (Qiagen) according to manufacturer instructions, with the exception that samples were homogenized in bead tubes using a TissueLyser II (Qiagen, Venlo, Netherlands) for ten minutes at 30/sec, rather than using the vortex adapter described in the protocol, before proceeding according to the protocol and eluting in 100 µL of elution buffer (Qiagen).

### *18S* rRNA library preparation and sequencing

Eukaryotic 18S rRNA amplicon libraries were generated using two rounds of PCR. In the first round of PCR, performed at the MU Metagenomics Center, 18S rRNA amplicons were constructed via amplification of the V4 and V5 regions of the 18S rRNA gene with universal primers (F566Euk/R1200Euk)^9^, flanked by Illumina standard adapter sequences. PCR was performed in 50 µL reactions containing 100 ng template DNA, primers (10 µM each), and EmeraldAmp Supermix PCR kits (Thermo Fisher). PCR cycling parameters were 95°C^(3 min)^ + [95°C^(30 sec)^ + 61°C^(30 sec)^ + 72°C^(60 sec)^] × 20 cycles + 72°C^(7 min)^. Libraries were then purified using Nucleospin Gel and PCR clean-up kits (Fisher Scientific) per the manufacturer’s protocol.

The second round of PCR, performed at the MU Genomics Technology Core, extended adapter sequences and added sample-specific nucleotide indices to both F and R primers to allow multiplexing. Here, PCR was performed in 50 µL reactions containing 10 ng template DNA, primers (2 µM each), dNTPs (10 mM), and Phusion Polymerase (Thermo Fisher). PCR cycling parameters were 98°C^(3 min)^ + [98°C^(15 sec)^ + 55°C^(30 sec)^ + 72°C^(45 sec)^] × 6 cycles + 72°C^(7 min)^. Amplicon pools (5 µL/reaction) were combined, thoroughly mixed, and then purified by addition of Axygen Axyprep MagPCR clean-up beads to an equal volume of 50 µL of amplicons and incubated for 15 minutes at room temperature. Products were then washed twice with 80% ethanol, and the dried pellet was resuspended in 32.5 µL EB buffer (Qiagen), incubated for two minutes at room temperature, and then placed on the magnetic stand for five minutes. The final amplicon pool was evaluated using the Advanced Analytical Fragment Analyzer automated electrophoresis system, quantified using quant-iT HS dsDNA reagent kits, and diluted according to Illumina’s standard protocol for sequencing as 2×500 bp paired-end reads on the MiSeq i100 instrument (Illumina).

### *1cS* rRNA library preparation and sequencing

Prokaryotic 16S rRNA amplicon libraries were constructed via amplification of the V4 region of the 16S rRNA gene with universal primers (U515F/806R), flanked by Illumina standard adapter sequences^10, 11^. Dual-indexed forward and reverse primers were used in all reactions. PCR was performed in 50 µL reactions containing 100 ng metagenomic DNA, primers (0.2 µM each), dNTPs (200 µM each), and Phusion high-fidelity DNA polymerase (1U, Thermo Fisher). Amplification parameters were 98 °C^(3 min)^ + [98 °C^(15 sec)^ + 50 °C^(30 sec)^ + 72 °C^(30 sec)^] × 25 cycles + 72 °C^(7 min)^. Amplicon pools (5 µL/reaction) were combined, thoroughly mixed, and then purified by addition of Axygen Axyprep MagPCR clean-up beads to an equal volume of 50 µL of amplicons and incubated for 15 minutes at room temperature. Products were then washed twice with 80% ethanol and the dried pellet was resuspended in 32.5 µL EB buffer (Qiagen), incubated for two minutes at room temperature, and then placed on the magnetic stand for five minutes. The final amplicon pool was evaluated using the Advanced Analytical Fragment Analyzer automated electrophoresis system, quantified using quant-iT HS dsDNA reagent kits, and diluted according to Illumina’s standard protocol for sequencing as 2×300 bp paired-end reads on the MiSeq i100 instrument (Illumina).

### Bioinformatics

Demultiplexed paired-end reads were processed using QIIME2 v2024.5^12^. Primers were designed to match the 5’ ends of the forward and reverse reads. Cutadapt^13^ (version 4.8) was used to remove the primer from the 5’ end of the forward read. If found, the reverse complement of the primer to the reverse read was then removed from the forward read as were all bases downstream. Thus, a forward read could be trimmed at both ends if the insert was shorter than the amplicon length. The same approach was used on the reverse read, but with the primers in the opposite roles. Read pairs were rejected if one read or the other did not match a 5’ primer, and an error-rate of 0.1 was allowed. Two passes were made over each read to ensure removal of the second primer. A minimal overlap of three bp with the 3’ end of the primer sequence was required for removal. The QIIME2^12^ DADA2^14^ plugin (version 1.10.0) was used to denoise, de-replicate, and count ASVs (amplicon sequence variants), incorporating the following parameters: 1) forward and reverse reads were truncated to 150 (16S rRNA) or 400 (18S rRNA) bases, 2) forward and reverse reads with number of expected errors higher than 2.0 were discarded, and 3) Chimeras were detected using the consensus method and removed. R version 3.5.1 and Biom version 2.1.7 were used in QIIME2. Taxonomies were assigned to final sequences using the SILVA.v144^15^ and EUKARYOME 3.0^16^ databases, using the classify-sklearn procedure^17^.

### Statistical analysis

Richness and alpha-diversity data were first tested for normality and equal variance using the Shapiro-Wilk test and Brown-Forsythe test, respectively. Group- and age-associated differences in richness and alpha-diversity were then tested using analysis of variance (ANOVA) or Kruskal-Wallis ANOVA on ranks. Main effects and interactions of database and taxonomic group on taxonomic resolution were tested using two-way ANOVA via a general linear model. Post hoc pairwise comparisons were performed using the Holm-Sidak method. Differences in beta-diversity were tested using permutational multivariate ANOVA (PERMANOVA), while homogeneity of dispersion was tested using permutational analysis of multivariate dispersions (PERMDISP). PERMANOVA and PERMDISP were conducted using MicrobiomeAnalyst 3.0^18^. Heat tree comparing dairy and beef heifers was generated using MicrobiomeAnalyst 3.0. Differential abundance (DA) testing was performed using one-way analysis of variance (ANOVA) followed by pairwise comparisons using Fisher’s least significant difference. Correction to control the false discovery rate (FDR) during DA testing was performed using the method of Benjamini and Hochberg^19^. Correlation between total 16S rRNA and 18S rRNA richness was tested using Spearman rank order (SRO) correlation. The correlation network analysis was performed using SRO with sparse correlations for compositional data (SparCC) using 100 permutations to control the FDR.

## Results

### *18S* rRNA metabarcoding of eukaryotic DNA

#### Data quality and diversity metrics

During quality control of 18S rRNA sequence data, three samples were removed from all subsequent analyses due to low read counts (0, 276, and 5079 sequence reads). The remaining samples yielded a mean (± SE) of 26,675 (± 1123) sequence reads per sample, and no difference was detected between groups in the number of sequence reads obtained (*p* = 0.373, F = 1.1; **Fig. 1A**). Thus, using all sequence reads in the remaining 29 samples, we first assessed and compared traditional taxonomy-independent metrics of community ecology, including richness, alpha-, and beta-diversity among the four groups. A comparison of the richness (i.e., number of distinct eukaryotic DNA sequences identified in each sample) failed to detect a significant difference (*p* = 0.295, F = 1.1; **Fig. 1B**). Similarly, no difference in alpha-diversity (i.e. Shannon index) was detected (*p* = 0.207; **Fig. 1C**).

**Figure 1.**
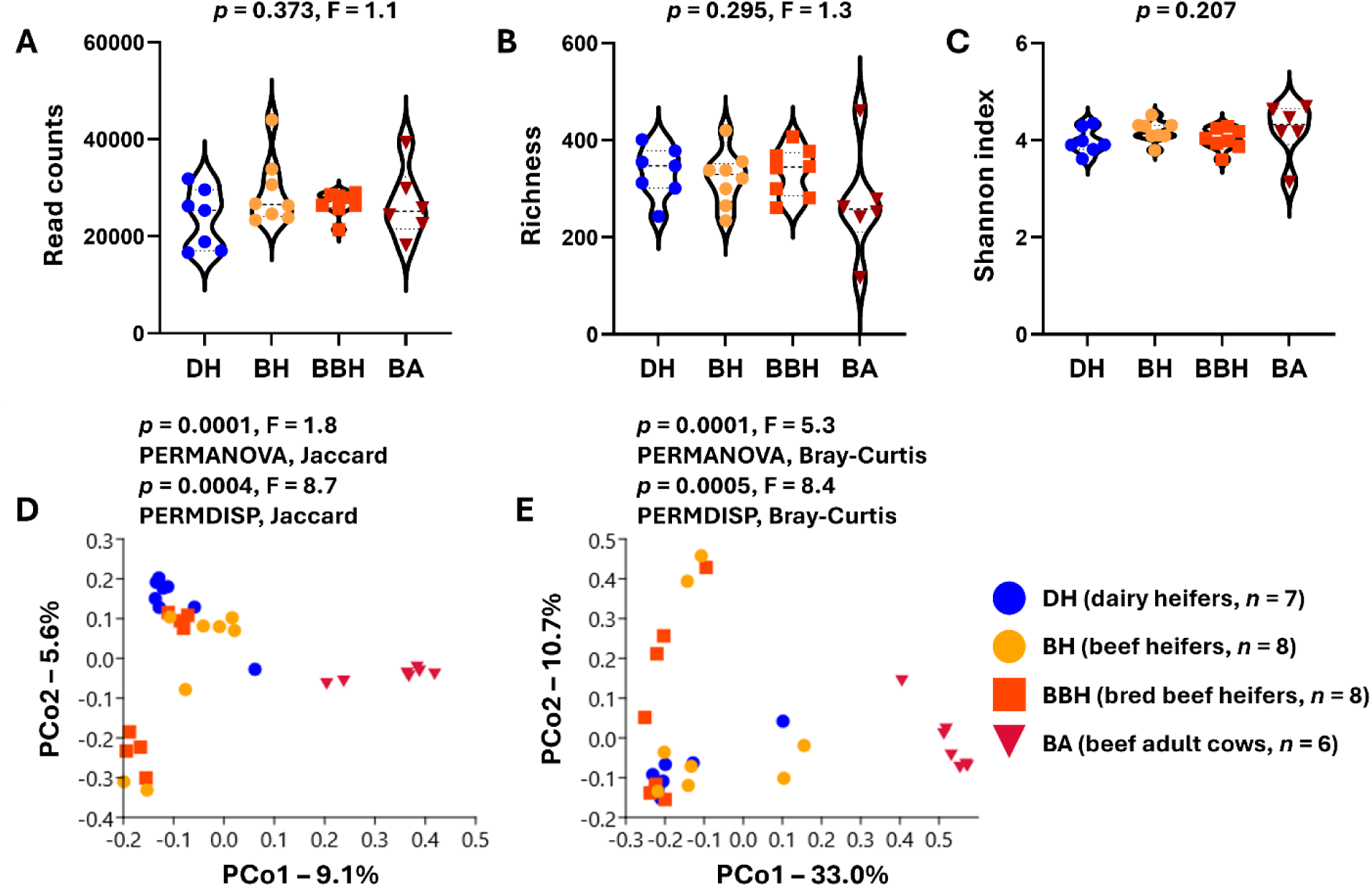
Violin plots showing total read counts (**A**), total richness (**B**), and Shannon diversity index (**C**) of eukaryotic 18S rRNA sequences isolated from the feces of 4-6-month-old dairy heifers (DH) and beef heifers (BH), 16-18-month-old bred beef heifers (BBH), and adult beef cows (BA) greater than 5 years old. Results of one-way ANOVA (**A**, **B**) or Kruskal-Wallis ANOVA on ranks (**C**) shown above each plot. (**D**, **E**) Principal coordinate plots showing beta-diversity among 18S rRNA sequences isolated from the feces of dairy heifers (4-6 months old, *n* = 7), beef heifers (4-6 months old, *n* = 8), bred beef heifers (16-18 months old, *n* = 8), and beef cows (> 5 years old, *n* = 6), ordinated using unweighted (**D**) and weighted (**E**) dissimilarities. Legend shown at right. Results of permutational multivariate analysis of variance (PERMANOVA) shown above each plot.

Beta-diversity among all samples was assessed using both unweighted (Jaccard) and weighted (Bray-Curtis) dissimilarities. Principal coordinate analysis (PCoA) based on unweighted dissimilarities (**Fig. 1D**) revealed clear separation of samples from BA from all other groups along principal coordinate 1 (PCo1) while the samples from DH, BH, and BBH separated along PCo2, albeit incompletely. Permutational multivariate ANOVA (PERMANOVA) using unweighted dissimilarities detected a significant but modest overall difference (*p* = 0.0001, F = 1.8), with significant differences in all pairwise comparisons (**Table S2A**) except between samples from beef heifers and bred beef heifers (*p* = 0.0568, F = 1.2). While permutational multivariate analysis of dispersion (PERMDISP) also detected a significant difference (*p* = 0.0004, F = 8.7), the distinct clustering on PCoA suggests true differences in beta-diversity. PCoA using weighted dissimilarities (**Fig. 1E**) yielded a similar outcome with clear separation between samples from BA and samples from all other groups. Results of PERMANOVA using weighted dissimilarities detected an even stronger overall effect (*p* = 0.0001. F = 5.3) and pairwise differences (**Table S2B**), and similar differences in dispersion (*p* = 0.0005, F = 8.4, PERMDISP).

### Database selection and eukaryotic taxonomic profile

To optimize taxonomic assignments, amplicon sequence variants (ASVs) were annotated using two databases, SILVA (S) and EUKARYOME (E), and compared in terms of the number and taxonomic resolution of annotations provided by each database. Following alignment and careful manual curation to ensure an accurate record of the taxonomic level achieved in each assignment, the number and level of all unique taxa were compared between SILVA- and EUKARYOME-generated assignments. Of the 6059 total ASVs, 944 (15.6%) achieved species-level assignments from both databases (**Fig. S1A**). Individually, 55.2% (3344/6059) and 49.1% (2976/6059) of ASVs were annotated to at least the level of genus with SILVA and EUKARYOME, respectively (**Fig. S1B**). Using the annotations from both databases in combination, 35.9% (2177/6059) of ASVs were annotated to the level of species, and 65.2% (3950/6059) of ASVs were annotated to at least the level of genus. To compare taxonomic resolution between databases and among different types of organisms, 55 ASVs receiving only domain-level assignments from both databases were removed, and the remaining 6004 ASVs were collapsed into 12 phyla or other groups based on EUKARYOME annotations (**Fig. S1C**). Two-way analysis of variance (ANOVA) identified significant effects of database (*p* = 0.017, F = 5.7) and taxonomic group (*p* < 0.001, F = 220.6), and a significant interaction effect (*p* < 0.001, F = 54.8). Pairwise comparisons found that both SILVA and EUKARYOME outperformed the alternative database in different groups of organisms. Specifically, SILVA provided superior taxonomic resolution of ASVs within *Straminipila*, *Alveolata*, and *Rhizaria*, while EUKARYOME provided superior resolution of ASVs within *Amoebozoa*, *Ascomycota*, and *Viridiplantae*. Independent of database, the best-resolved taxonomic groups included *Neocallimastigomycota*, *Mucoromycota*, and *Basidiomycota*, while *Straminipila*, *Alveolata*, *Rhizaria*, *Amoebozoa*, and *Ascomycota* were also well-annotated. In contrast, ASVs within kingdoms *Metazoa* and *Viridiplantae* and the two groups of fungal and non-fungal ASVs not contained in the other ten groups tended to receive assignments at lower levels.

### Taxonomic profiling of eukaryotic fecal microbiome of cattle

Using those taxonomic assignments, our goals were to identify dominant sources of eukaryotic DNA at the phylum- and genus-level, to identify environment- and age-dependent differences in the fecal eukaryome of cattle, and to assess the utility of the current assay as a tool for surveillance of problematic eukaryotic organisms encountered in livestock management.

Visualization of the phylum-level composition via stacked bar charts (**Fig. 2A**) revealed several interesting findings. First, fungi (phyla *Ascomycota*, *Basidiomycota*, *Mucoromycota*, *Neocallimastigomycota*, and *Rozellomycota*) were the dominant source of eukaryotic DNA in the feces of DH, BH, and BBH, while protists (phyla *Evosea*, *Parabasalida*, *Blastocystis*, and *Cercozoa*) represented the dominant source of eukaryotic fecal DNA in the BA group. Other taxa selectively enriched in one or more groups included *Nematoda* in DH and *Ciliophora* in BH and BBH. Annotated to the level of genus (**Fig. 2B**), the dominant fungal taxa were identified as *Thermomyces*, *Talaromyces*, and *Aspergillus* (phylum *Ascomycota*); *Tausonia*, *Tritirachium*, and *Wallemia* (phylum *Basidiomycota*); *Lichtheimia* and *Mucor* (phylum *Mucoromycota*); *Cyllamyces* (phylum *Neocallimastigomycota*); and *Nosematida* (phylum *Rozellomycota*). The dominant protozoal genera detected in BA samples were *Entamoeba*, *Tetratrichomonas*, *Simplicimonas*, and *Blastocystis*, while the *Ciliophora* identified at high RA in several BH and BBH samples was annotated as *Buxtonella*.

**Figure 2.**
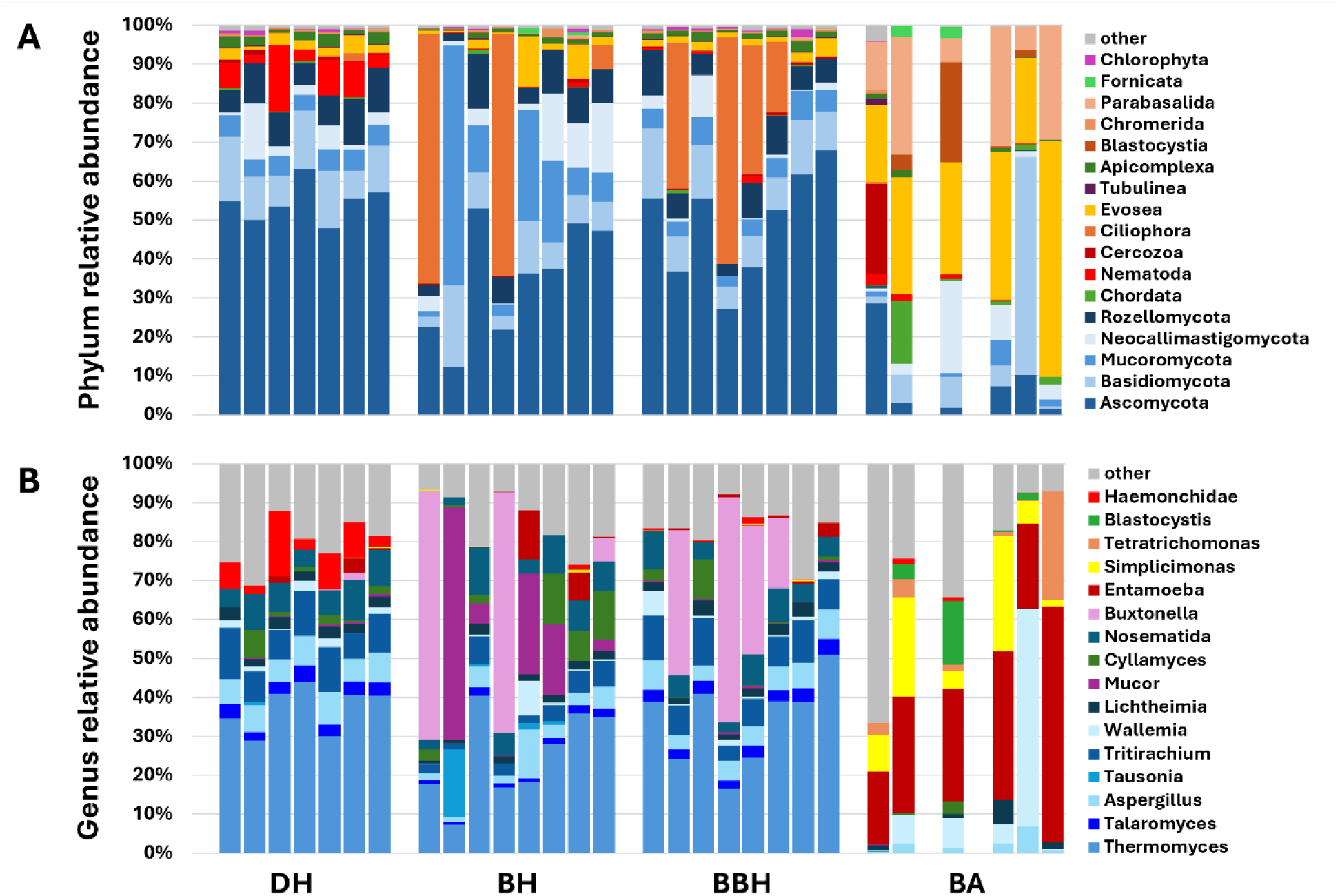
Stacked bar charts showing the relative abundance of the 14 dominant phylum-level eukaryotic annotations with a mean RA > 1% (**A**), and the 16 dominant genus-level annotations with a mean RA > 2% (**B**) in DNA isolated from the feces of dairy heifers (DH), beef heifers (BH), bred beef heifers (BBH), and adult beef cows BA). Legends at right. The remaining 37 phyla or 427 genera are combined as ‘other’.

To identify differentially abundant 18S rRNA amplicon sequence variants (ASVs), one-way analysis of variance (ANOVA) was performed with FDR correction using the method of Benjamini and Hochberg and Fisher’s least significant difference (LSD) test for pairwise comparisons among all groups. A total of 32 ASVs withstood correction for multiple tests, representing 13 different taxonomic annotations (**Table S3**). Summarized, these included the enrichment of *Entamoeba bovis* and *Neocallimastix frontalis* in BA; the depletion of *Aspergillus*, *Penicillium*, *Talaromyces*, *Thermomyces lanuginosus*, *Bulbithecium pinkertoniae*, *Microascus cirrosus*, *Chaetomium*, and *Tritirachium* in BA; and the enrichment of *Haemonchidae* (including *Cooperia oncophora*) in DH.

Regarding the presence of eukaryotic parasites, pathogens, and intoxicants, several organisms of concern were detected among the kingdom fungi. DNA from *Aspergillus* was present in all samples, typically at high RA (**Fig. 2B**). Also from the phylum *Ascomycota*, DNA from *Scopulariopsis brevicaulis*, associated with generalized hyperkeratosis in calves^20^ and onychomycoses in humans^21^, was identified in all samples from BA but was absent from other groups. *Basidiomycota* opportunists associated with otitis externa (*Malassezia*) and mastitis (*Trichosporon*) were detected in three samples at low RA. Notably, several causative agents of bovine mucormycosis including *Mucor*, *Lichtheimia*, *Rhizomucor*, and *Rhizopus* spp. were identified in all groups, with particularly high RA of *Mucor* DNA in samples from beef heifers. Samples from DH, BH, and BBH all contained substantial DNA annotated to *Lichtheimia corymbifera*, *Rhizomucor pusilus*, and *Rhizomucor miehei*.

In addition to opportunistic fungi, several sources of mycotoxins from the phylum *Ascomycota* were detected. *Fusarium* was resolved to the genus level in three samples, one beef heifer, one bred beef heifer, and one adult beef cow, while the family containing *Fusarium* (*Netriaceae*) was detected in 65.6% (19/29) of all samples. *Chaetomium* was also identified at high RA in all samples except those from adult beef cows. DNA from the xerophile *Wallemia sebi* (phylum *Basidiomycota*) was detected in all samples, at extremely high RA in one sample from an adult beef cow. In addition to mycotoxin production, *Wallemia sebi* is the source of airborne spores associated with hypersensitivity pneumonitis in cattle and Farmer’s Lung disease in humans^22^.

Several opportunistic protozoal organisms were detected, including members of the phyla *Apicomplexa* (*Cryptosporidium andersoni*, *Eimeria bovis*, *E. cylindrica*), *Ciliophora* (*Buxtonella sulcata*), *Amoebozoa* (*Entamoeba bovis*), and *Straminipila* (*Blastocystis*).

Assignments to metazoan parasites and opportunists included *Cooperia oncophora*, *Trichostrongylus colubriformis*, genus *Ostertagia*, rumen flukes (*Orthocoelium orthocoelium* and *Calicophoron daubneyi*, family *Gastrothylacidae*), along with several free-living soil and water nematodes (*Halicephalobus gingivalis*, *Cephaloboides nidrosiensis*, *Distolabrellus veechi*, *Oscheius tipulae*) and plant parasites (*Cactodera chenopodiae*), likely representing pass-through following inadvertent ingestion while grazing. Similarly, DNA from several plants was detected, including families and genera of grasses (*Poaceae*, *Brachypodium distachyon*, *Juncus effusus*), sedges (*Carex*), legumes (*Fabacieae*), and other plants (*Typha angustifolia*) and trees (*Fagaceae*, *Ulmus*, and *Quercus*), presumably remnants of dietary roughage.

### *1CS* rRNA metabarcoding of prokaryotic DNA

#### Data quality and diversity metrics

During quality control of 16S rRNA sequence data, two samples were removed from subsequent analyses due to low read counts (1407 and 1753 sequence reads). The remaining samples yielded a mean (± SE) of 205,944 (± 37,600) sequence reads per sample with no difference between groups in the number of sequence reads obtained (*p* = 0.430, F = 1.0; **Fig. 3A**). A significant difference in richness was detected (*p* = 0.042) with the only significant pairwise comparison between BBH and BA (**Fig. 3B**). No difference in alpha diversity was detected (**Fig. 3C**, *p* = 0.087, F = 2.4). In contrast, 16S rRNA beta diversity differed significantly between all groups. Following PCoA using unweighted dissimilarities, BA and DH each formed discrete clusters while BH and BBH partially overlapped (**Fig. 3D**). PERMANOVA detected a significant main effect (*p* = 0.0001, F = 5.2) and significant differences in all pairwise comparisons (**Table S2C**). A similar pattern was observed using weighted dissimilarities (**Fig. 3E**) with similar outcomes of PERMANOVA (*p* = 0.0001, F = 6.5; **Table S2D**). As with the pairwise comparisons of beta diversity based on 18S rRNA ASVs, the greatest F values were obtained in comparisons between BA and other groups, particularly BH. Testing for differences in dispersion of 16S rRNA ASVs also revealed significant differences (*p* = 0.0001, F = 10.5, Jaccard; *p* = 0.00009, F = 10.8, Bray-Curtis; PERMDISP).

**Figure 3.**
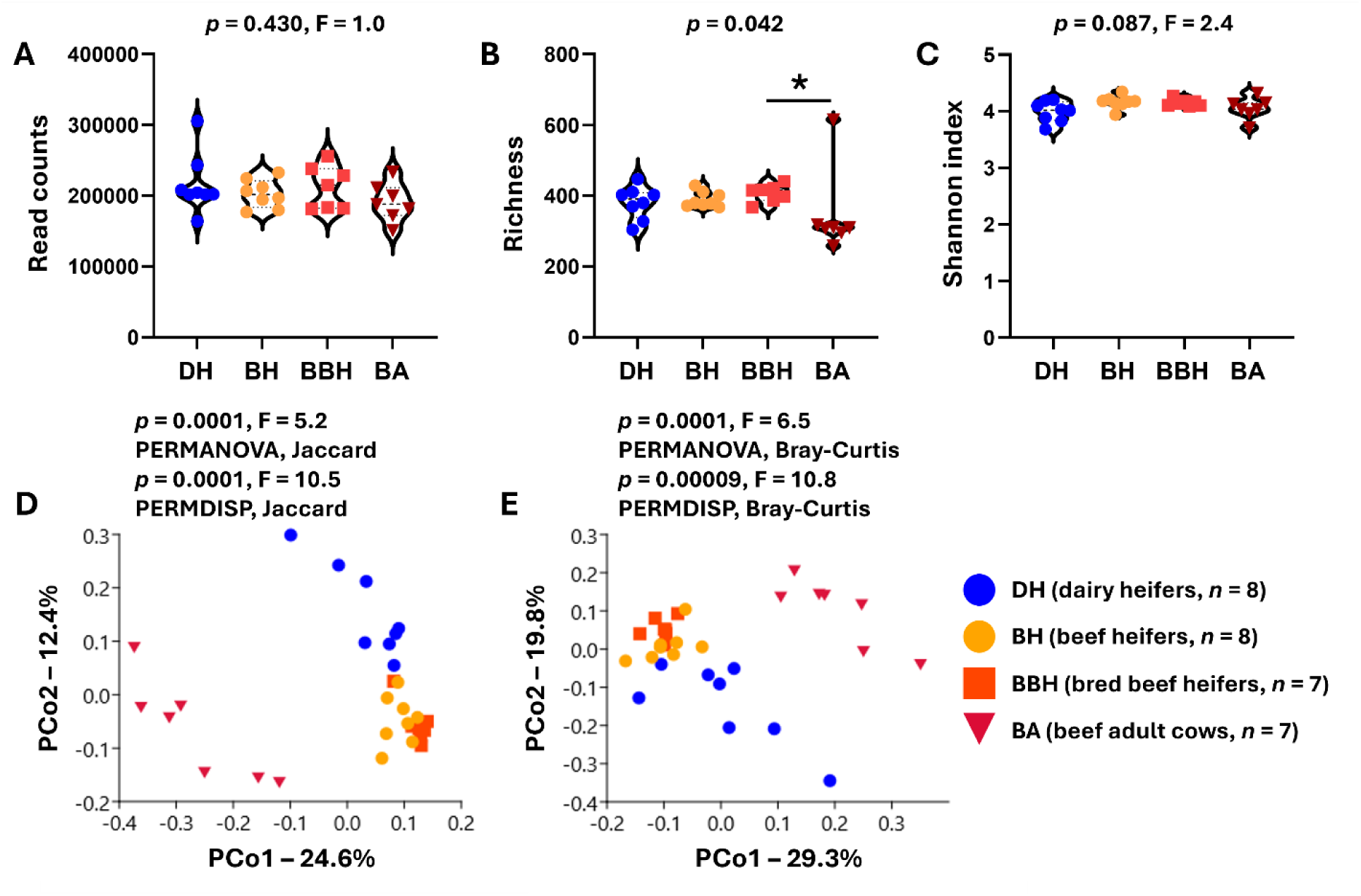
Violin plots showing total read counts (**A**), total richness (**B**), and Shannon diversity index (**C**) of prokaryotic 16S rRNA sequences isolated from the feces of 4-6-month-old dairy heifers (DH) and beef heifers (BH), 16-18-month-old bred beef heifers (BBH), and adult beef cows (BA) greater than 5 years old. Results of one-way ANOVA (**A**, **C**) or Kruskal-Wallis ANOVA on ranks (**B**) shown above each plot. (**D**, **E**) Principal coordinate plots showing beta-diversity among 16S rRNA sequences isolated from the feces of dairy heifers (4-6 months old, *n* = 7), beef heifers (4-6 months old, *n* = 8), bred beef heifers (16-18 months old, *n* = 8), and beef cows (> 5 years old, *n* = 6), ordinated using unweighted (**D**) and weighted (**E**) dissimilarities. Legend shown at right. Results of permutational multivariate analysis of variance (PERMANOVA) shown above each plot.

### Taxonomic profiling of prokaryotic fecal microbiome of cattle

Based on taxonomic assignments of the 16S rRNA sequence data, our goals were to identify dominant sources of prokaryotic DNA at the phylum- and genus-level, and to identify environment- and age-dependent differences in the prokaryotic fecal microbiome of cattle. At the level of phylum, differences between BA and the other groups were readily apparent in stacked bar charts, including a loss of *Actinomycetota* and *Euryarchaeota* and enrichment of *Pseudomonadota* and *Spirochaetota* (**Fig. 4A**). Similar charts showing ASVs collapsed to the level of genus suggested a depletion of *Bifidobacterium* and *Methanobrevibacter* in BA relative to the other groups (**Fig. 4B**).

**Figure 4.**
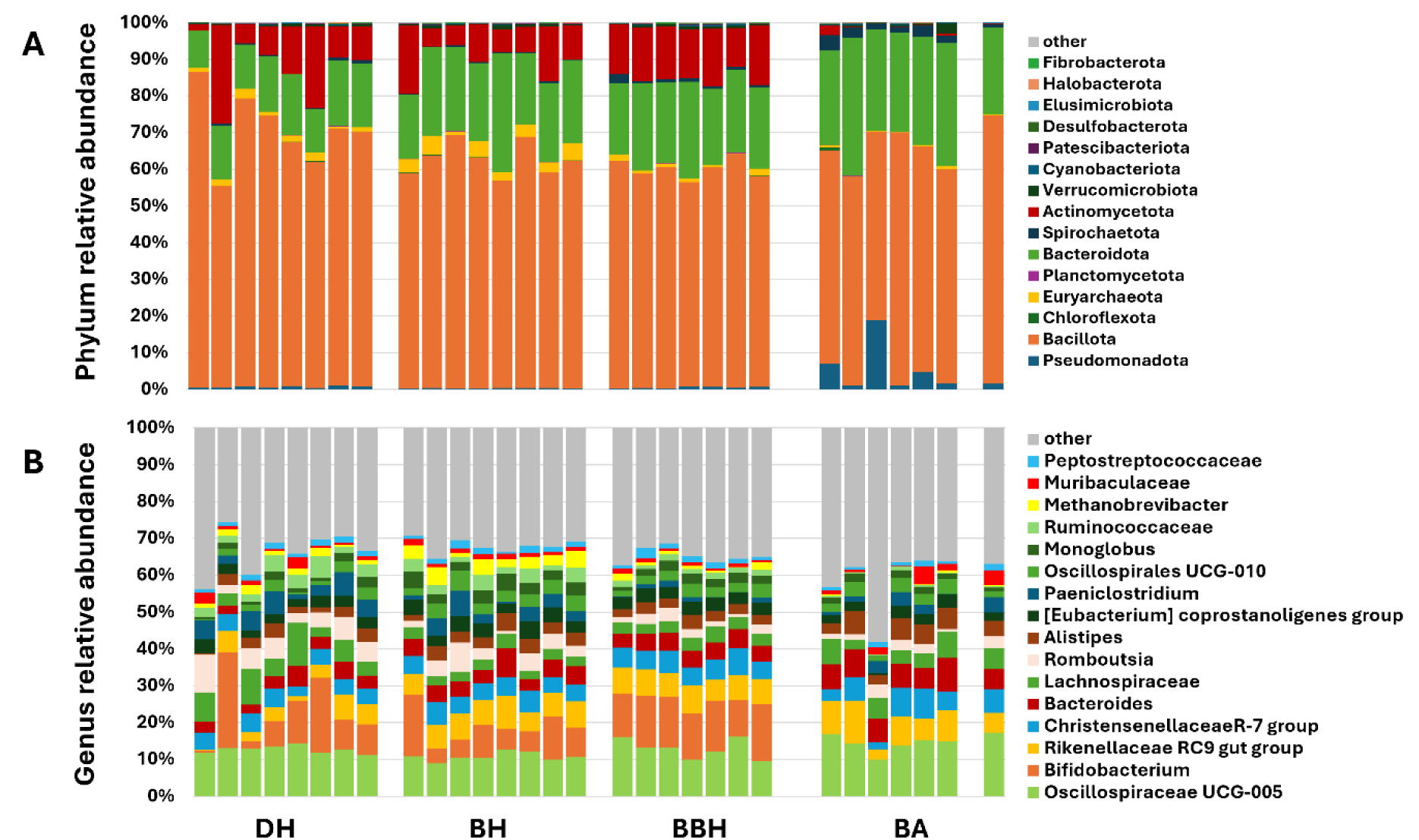
Stacked bar charts showing the relative abundance of the 14 dominant phylum-level prokaryotic annotations with a mean RA > 1% (**A**), and the 16 dominant genus-level annotations with a mean RA > 2% (**B**) in DNA isolated from the feces of dairy heifers (DH), beef heifers (BH), bred beef heifers (BBH), and adult beef cows BA). Legends at right. The remaining 17 phyla or 657 genera are combined as ‘other’.

Differential abundance testing of 16S rRNA ASVs yielded 167 ASVs with FDR-corrected *p* < 0.05 (**Table S4**), with 20 ASVs enriched in DH, 23 enriched in BH, 83 enriched in BBH, and 41 enriched in BA, relative to at least one other group. The ASVs enriched in DH and BH both comprised annotations from five phyla or classes, *Actinomycetota*, *Clostridia*, *Bacilli*, *Bacteroidota*, and *Euryarchaeota* (**Fig. S2A, B**). *Actinomycetota* enriched in DH and BH included *Aeriscardovia* (DH), an uncharacterized genus within family *Bifidobacteriaceae* (BH), and seven taxa within order *Coriobacteriales*. *Bacilli* enriched in DH were all identified as members of family *Lactobacillaceae*, while *Bacilli* enriched in BH were annotated as members of *Bacillaceae*, *Planococcaceae*, and *Erysipelotrichaceae*.

ASVs enriched in BBH represented a wider phylogenetic range of organisms including *Actinomycetota*, *Clostridia*, *Bacilli*, *Negativicutes*, *Symbiobacteriia*, *Bacteroidota*, *Pseudomonadota*, *Spirochaetota*, *Patescibacteriota*, and sulfate-reducing bacteria (**Fig. S2C**). In contrast to DH and BH, *Actinomycetota* enriched in BBH were predominantly members of class *Actinobacteria*, including *Bifidobacterium*, *Saccharopolyspora*, *Streptomyces*, *Thermobifida*, *Thermopolyspora*, and *Thermobispora*. *Bacilli* enriched in BBH included eight genera or species within the orders *Bacillales* (6 ASVs) or *Brevibacillales* (2 ASVs), five members of the *Erysipelotrichales*, *Aerococcus vaginalis* (order *Lactobacillales*), three members of order *Paenibacillales* including *Thermobacillus composti*, and genus *Thermoactinomyces*. *Bacteroidota* enriched in BBH included three ASVs annotated as *Bacteroidales* RF16 group, four ASVs annotated as members of family *Prevotellaceae*, and ASVs annotated as *Paludibacteracae*, *Dysgonomonadaceae*, *Rikenellaceae*, and the uncultured group p-251-o5. The member of *Negativicutes* enriched in BBH was identified as genus *Phascolarctobacterium*. The sulfate-reducing bacteria (SRB) enriched in BBH were identified as *Mailhella*. The *Spirochaetota* enriched in BBH was annotated as genus *Sphaerochaeta*. *Pseudomonadota* enriched in BBH were all members of class *Alphaproteobacteria*, including genus *Methylobacterium*-*Methylorubrum*.

ASVs enriched in BA were annotated as members of *Clostridia*, *Bacilli*, *Negativicutes*, *Bacteroidota*, *Pseudomonadota*, and *Spirochaetota*. ASVs within *Bacilli* enriched in BA include *Anaeroplasma* and uncultured EMP-G18 group (family *Acholeplasmataceae*), two members of the *Erysipelotrichales*, and *Lactobacillus ruminis*. ASVs from the phylum *Bacteroidota* enriched in BA included the genera *Bacteroides*, *Parabacteroides*, *Alistipes*, and uncultured members of order *Bacteroidales* and families *Prevotellaceae* and *Rikenellaceae*. The *Negativicutes* enriched in BA included genus *Anaerovibrio* and *Megasphaera elsdenii*. The three ASVs within *Spirochaetota* enriched in BA were annotated as *Treponema berlinense* or genus *Treponema*. Genus *Sutterella* and unclassified member of order *Rhodospirillales* were the *Pseudomonadota* enriched in BA.

In all four groups, class *Clostridia* represented the dominant source of enriched ASVs. Within *Clostridia*, closely related ASVs from several families were often enriched in multiple groups. For example, three distinct ASVs from family *Lachnospiraceae*, each annotated as *Ruminococcus gauvreauii* group were each selectively enriched in BH, BBH, or BA, while a fourth ASV annotated as *Ruminococcus torques* group was enriched in DH. Multiple other pairs and trios of ASVs from the families *Lachnospiraceae*, *Ruminococcaceae*, and *Anaerovoracaceae* with identical annotations were enriched in multiple groups. In contrast, several ASVs annotated as members of family *Oscillospiraceae* were selectively enriched in either the BBH or BA groups.

### Multi-domain correlation analyses

Next, data from the two metabarcoding assays were combined to identify correlations in richness and relative abundance of features, and possible biological relationships among bacterial, fungal, protozoal, and other organisms. Using Spearman’s rank order correlation, a significant correlation was detected between eukaryotic and prokaryotic richness (*p* = 0.0154, R^2^ = 0.462, **Fig. S3**). To identify significant positive and negative correlations in the relative abundance (RA) of biological features, the species-level assignments from SILVA of all 18S rRNA and 16S rRNA ASVs were used in a single correlation network analysis. With thresholds of *p* < 0.05 and rho > 0.5, and 100 permutations using sparse correlations for compositional data (SparCC) to eliminate false positives, a robust correlation network was generated containing ASVs annotated as bacteria, fungi, protozoa, and metazoa (**Fig. 5A**, **Table S5**). Network nodes with bacterial assignments represented a wide phylogenetic range but were enriched in thermophilic organisms (*Symbiobacterium thermophilum*, *Thermasporomyces composti*, *Thermomonospora curvata*, *Brevibacillus thermoruber*, *Bacillus thermoamylovorans*, *Thermobacillus composti*, *Thermobispora bispora*, *Thermobifida fusca*, and uncultured *Thermobacillus*) and other members of class *Bacilli* (*Lactobacillus mucosae*, *L. amylovorus*, *Caenibacillus caldisaponilyticus*, *Paenibacillus barengoltzii*, *Brevibacillus limnophilus*, *Bacillus coagulans*, *B. smithii*, and *B. vietnamensis*). All pairwise comparisons of thermophilic bacteria revealed significant positive correlations while the relationships between thermophiles and other bacteria and eukaryotic features varied (**Fig. 5B**). Consistent negative correlations were identified between the majority of thermophilic bacteria and certain features present at high RA in the adult beef cows including *Neocallimastigaceae*, *Simplicimonas*, and *Bacillus vietnamensis*. Prominent nodes with eukaryotic assignments included *Entamoeba*, *Simplicimonas*, *Neocallimastigaceae*, and *Fusarium*. With the exception of a strong positive correlation between the RA of *Entamoeba* and *Simplicimonas*, there were no significant correlations among these four features (**Fig. 5C**). Nonetheless, all four features showed strikingly similar (and somewhat selective) negative correlations with thermophilic bacteria and members of class *Bacilli*. These findings integrate the separate results of differential abundance testing performed with 18S rRNA and 16S rRNA data and highlight potential inter-domain relationships involved in the maturation of the bovine fecal microbiome.

**Figure 5.**
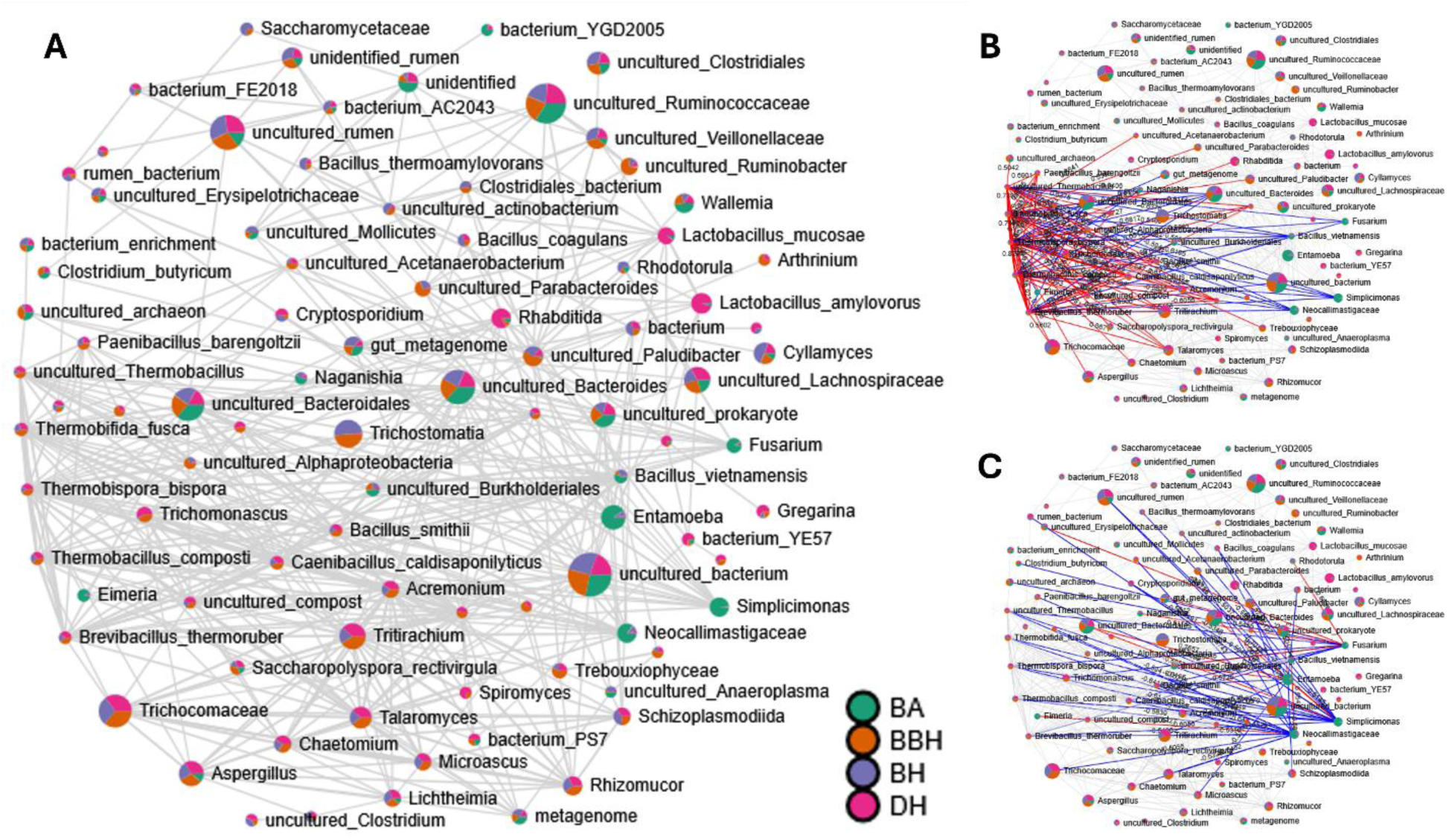
Correlation network showing all significant (p < 0.05) correlations with rho > 0.5 (Spearman rank order correlation) between species-level assignments of 16S rRNA and 18S rRNA data annotated using SILVA v144. Nodes are pie charts showing mean relative abundance of each taxon among the four groups, legend at lower right (**A**). Identical clusters showing the positive (red) and negative (blue) correlations associated with five 16S rRNA amplicon sequence variants (ASVs) annotated as thermophilic bacteria (uncultured *Thermobacillus*, *Thermobifida fusca*, *Thermobispora bispora*, *Thermobacillus composti*, and *Brevibacillis thermoruber*) (**B**) and four 18S rRNA ASVs annotated as protozoa (*Entamoeba*, *Simplicimonas*) or fungi (*Neocallimastigaceae*, *Fusarium*) (**C**).

## Discussion

These data demonstrate the combined prokaryotic and eukaryotic diversity of the bovine hindgut, and differences between age-matched dairy and beef heifers, and between beef cattle of different ages. While numerous studies have investigated the effects of age, diet, and other factors on the rumen and fecal microbiome of dairy or beef cattle^23–27^, surprisingly few have directly compared the gut microbiome of age-matched dairy and beef cattle. One study comparing the fecal microbiome of dairy and beef calves at four weeks of life using 16S rRNA metabarcoding failed to detect a significant difference^28^. In contrast, several studies have used metagenomic approaches to identify differences in antimicrobial resistance genes (ARGs) in the feces of adult dairy and beef cattle^29–31^.

Similarly, most studies focus on the bacterial microbiome of cattle, with only a few studies using parallel metabarcoding platforms to analyze prokaryotic and eukaryotic components of the same samples. Rumen fluid analyses in Italian Simmental cows (*N* = 7) during different stages of lactation using 16S rRNA (V3-V4) and 18S rRNA (V9) metabarcoding identified *Ciliophora* as the dominant eukaryotic taxon with very minor contributions from *Amoebozoa*, *Chytridiomycota*, and other taxa^32^. Fungi were apparently not resolved with this set of primers. Estimates of rumen bacterial richness in that study (range of means, 464 to 692) were slightly higher than our estimates of fecal richness (mean ± SE, 391 ± 12), while their estimates of eukaryotic richness (range of means, 40 to 68) were far less than our estimates of eukaryotic richness (mean ± SE, 321 ± 14). Similarly, a study of Angus (*n* = 5), dual-purpose Braunvieh (*n* = 3), and Simmental×Braunvieh F1 crossbreed (*n* = 3) cattle analyzed using 16S rRNA (V4) and 18S rRNA (V4) metabarcoding found significant breed-associated differences in bacterial, fungal, and protozoal fecal communities^33^. Again however, observed fungal richness was much lower than that of the present study. Notably, two studies of the fecal bacteria and fungi^34^, or protists and fungi^35^, in Peruvian Simmentals (*N* = 21) using combinations of 16S rRNA (V4), ITS2, and 18S rRNA (V4) metabarcoding contained male and female subjects in similar age groups to the present study. While PERMANOVA of bacterial and fungal ASVs indicated significant differences between age groups, modest *p* values and minimal separation during PCoA suggested subtle age-associated differences in beta diversity. No age-dependent difference in protozoal beta-diversity was detected^35^. While several phyla of fungi were resolved with their ITS2 primers, observed fungal richness was roughly half that observed in the present study. The shift from a fungi-dominated to protozoa-dominated eukaryotic microbiome in adulthood was not reported in those studies^33, 35^. Given the strong influence of protozoa on enteric methane production in ruminants^36–40^, a shift of this nature may contribute to the significant increase in methane production between 10 months and 4 years of age^41^.

None of the aforementioned studies resolved any larger metazoan organisms such as nematodes and trematodes. ITS2-based metabarcoding has been used successfully to survey gastrointestinal nematodes in Canadian and Brazilian beef herds^42^. Notably, the majority of parasites identified in Canadian cattle were identified as *Cooperia oncophora* and *Ostertagia ostertagi*, the dominant nematode annotations in the present study.

The 18S rRNA data also demonstrate the ability to identify DNA from organisms colonizing the foregut of cattle, such as *Cooperia* and *Ostertagia* which exclusively colonize the duodenum and abomasum, and *Orthocoelium orthocoelium* and *Calicophoron daubneyi* (rumen flukes) which colonize the rumen and reticulum. Similarly, DNA from other nematodes detected inconsistently and at lower relative abundance, including *Oscheius tipulae*, *Panagrolaimus*, *Distolabrellus veechi*, *Halicephalobus gingivalis*, *Cephaloboides*, and *Heteroderidae*, likely represent pass-through DNA from accidental ingestion during grazing as these are free-living soil nematodes.

While DNA from *Ascomycota* and other phyla of fungi are frequently identified in bovine rumen and fecal samples^43^, the majority of obligately anaerobic gut fungi colonizing the GI tract of ruminants belong to the phylum *Neocallimastigomycota*^44, 45^. DNA from several members of this phylum was detected in the present study at high prevalence including *Cyllamyces aberensis*, *Neocallimastix frontalis*, *Pecoramyces ruminantium*, and genus *Piromyces*. *Ascomycota* present at high relative abundance included aerobic but heat-tolerant organisms including *Thermomyces*, *Talaromyces*, and *Tritirachium*, the spores of which are capable of surviving passage through the ruminant GI tract. While speculative, many of the 18S rRNA and 16S rRNA ASVs enriched in BBH suggest an influence of core body temperature on the survival or growth of gastrointestinal microbes during pregnancy. First, 34.4% (11/32) of the differentially abundant 18S rRNA ASVs were identified as *Thermomyces laguninosus*, and all of them were enriched in BBH. Similarly, 9.6% (8/83) of the 16S rRNA ASVs enriched in BBH were identified as thermophilic bacteria, including members of *Actinomycetota* (*Thermobifida fusca*, *Thermobispora bispora*, and genus *Thermopolyspora*) and *Bacillota* (*Thermobacillus composti*, and genus *Thermoactinomyces*). Due to the greater metabolic rate during growth and pregnancy, the thermogenic effects of progesterone^46^, and fetal heat production, the body temperature of BBH would presumably exceed that of BA. Even small changes in body temperature may facilitate the survival of aerobic but thermophilic fungi like *T. laguninosus* or growth of thermophilic bacteria.

The detection of problematic environmental fungi suggests this approach may also provide information on feed quality. For example, the *Aspergillus* identified in all samples likely represents spores present in feed. Other genera associated with mycotoxin production, including *Fusarium*, *Penicllium*, and *Wallemia*, were also detected. While *Wallemia* can cause a fatal toxicosis, mycotoxicosis more commonly present with non-specific signs including reduced feed intake, milk yield, or weight gain; intermittent diarrhea; and poor reproductive performance. The spores from *Fusarium* and *Wallemia* have also been implicated as causative agents^47^ of the granulomatous interstitial pneumonitis affecting agricultural workers (i.e., Farmer’s lung^48^).

The ciliate protozoa *Buxtonella sulcata* was detected sporadically at extremely high RA exclusively in young and bred beef heifers, but not in dairy heifers or adult beef cows. With a worldwide distribution, *Buxtonella* can act as an opportunistic pathogen, causing diarrhea and other clinical signs^49^. While numerous surveys and reports of *Buxtonella sulcata* in cattle have emerged from Asian^49–54^, European^55–58^, South American^59^, and African^60^ countries, the two reports from cattle in North America were published long before the molecular age^61, 62^, suggesting this may be an underreported organism in U.S. livestock.

*Simplicimonas similis* was detected sporadically at low RA in all groups except adult cows, in which it was detected at high RA in all samples. *Simplicimonas* is a parabasalid found in bovine vaginal swabs capable of cross-reacting with *Tritrichomonas foetus* in certain diagnostic assays^63^. Similarly, another parabasalid protozoa, *Tetratrichomonas*, was also identified only in adult beef cows. The prominent position of *Simplicimonas* in the correlation network with numerous significant negative correlations with bacterial features suggests potential roles for protozoa like *Simplicimonas* and *Entamoeba* in the maturation of the bovine hindgut microbiome.

The genus *Bifidobacterium* was reduced in BA relative to all other groups, including BBH. Recognized as a component of colostrum and the early life microbiome of many mammals, these data imply *Bifidobacterium* persists in the bovine hindgut at least beyond 1.5 years of age before receding to undetectable levels by a few years later. In contrast, ASVs enriched in BA included gram-negative members of *Spirochaetota*, *Pseudomonadota*, and *Bacteroidota*; and taxa phylogenetically related to gram-positive bacteria but lacking traditional cell walls including *Anaeroplasma* (class *Bacilli*), *Anaerovibrio*, and *Megasphaera elsdenii* (both class *Negativicutes*).

Regarding overall performance, the 18S rRNA metabarcoding assay provided excellent taxonomic resolution across a wide range of organisms, including fungi, protozoa, and nematodes. Using a combination of databases for more exhaustive annotation, over 65% of all ASVs received a genus-level assignment or better. It is worth noting that many annotations at lower levels (e.g., *Aspergillaceae*) were accompanied by higher level annotations (e.g., *Aspergillus*) in the same sample. In such cases, it is unclear whether the lower level assignments represent separate, less clearly resolved organisms or degraded genetic material from lysed cells. Database-dependent performance varied between different organisms, suggesting the value of similar comparisons in samples of unknown composition.

Despite several compelling findings, there are also limitations to the current study. First, the study lacks technical controls including negative reagents and positive mock community standards processed alongside the experimental samples. Second, samples were collected at a single time-point, during the season with a relatively low environmental parasite burden. It is unclear whether the outcomes reported here would be reproducible if the study were performed during mid-summer. Third, as with all metabarcoding assays, the presence of DNA does not necessarily reflect viable organisms. While bacterial DNA detected in feces is often interpreted as evidence of hindgut colonization, fecal eukaryotic DNA must be interpreted more carefully given that many fungi, nematodes, and protozoa are free-living environmental organisms incapable of surviving the anaerobic hindgut. Lastly, metabarcoding assays are often biased toward or against certain organisms

In conclusion, the current 18S rRNA metabarcoding assay offered outstanding taxonomic resolution of a wide range of protozoa and fungal organisms, as well as nematodes and trematodes, thus providing a means to interrogate several groups of organisms of relevance to human and animal health with a single assay. The use of 16S rRNA and 18S rRNA metabarcoding assays in tandem provided comprehensive information regarding the bacterial, archaeal, protozoal, fungal, and metazoan compartments of the bovine hindgut in the midwest United States. The differences between groups underscore the combined effects of age and environment, and suggest that the total community structure is not fully mature by 1.4 years of age. Prokaryotic and eukaryotic features selectively enriched in the BBH suggest an interaction or combined effect of growth and gestation. Future work is needed to better document the maturation of the eukaryotic fecal community across larger cohorts of dairy and beef cattle, and determine the effect of factors like diet, geography, and climate.

## Data availability statement

All sequence data supporting the present study are available at the National Center for Biotechnology Information, Sequence Read Archive as BioProjects PRJNA1517228 (18S rRNA) and PRJNA1529803 (16S rRNA).

## Ethics statement

All research performed here was conducted under the approval of the University of Missouri Animal Care and Use Committee, exemption 70081.

## Supplemental material

**Table S1.** Details related to animals and samples included in the current study

**Table S2.** Results of permutational multivariate analysis of variance (PERMANOVA) using unweighted (Jaccard) and weighted (Bray-Curtis) dissimilarities among 18S rRNA (**A**, **B**) and 16S rRNA (**C**, **D**) amplicon sequence variants detected in feces of dairy heifers (DH), beef heifers (BH), bred beef heifers (BBH), and adult beef cows (BA). Main effects for each test are followed to the right by p and F values for the respective pairwise comparisons.

**Table S3.** Results of differential abundance testing of 18S rRNA (eukaryotic) amplicon sequence variants via analysis of variance (ANOVA) followed by pairwise comparisons using Fisher’s least significant difference test.

**Table S4.** Results of differential abundance testing of 16S rRNA (prokaryotic) amplicon sequence variants via analysis of variance (ANOVA) followed by pairwise comparisons using Fisher’s least significant difference test.

**Table S5.** Results of correlation analyses of 16S rRNA and 18S rRNA metabarcoding data among all samples using Spearman’s rank correlation. Only comparisons with raw p value < 0.05 and rho > 0.5 are listed.

**Figure S1.**
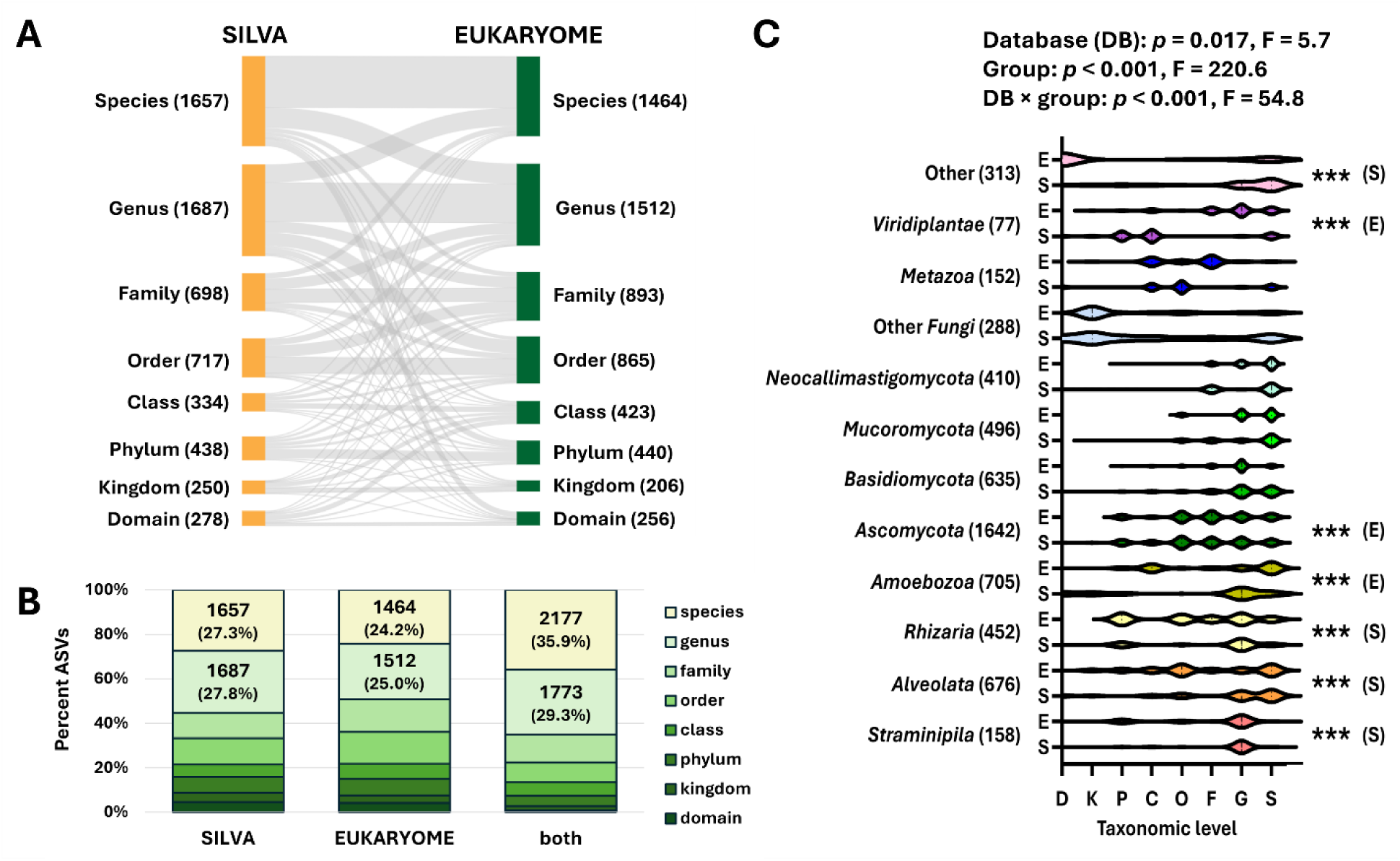
Sankey plot showing the number and relationship of 879 total non-arthropod amplicon sequence variants (ASVs) annotated at each taxonomic level using SILVA (left) or EUKARYOME (right) (**A**). Stacked bar charts showing the proportion of ASVs annotated at each taxonomic level using SILVA, EUKARYOME, or both sets of annotations. Number and percentage of ASVs annotated to the level of species or genus shown on plot (**B**). Violin plots showing the proportion of taxonomic assignments at each level from EUKARYOME (E) or SILVA (S) within the *Straminipila*, *Alveolata*, *Rhizaria*, *Amoebozoa*, *Ascomycota*, *Basidiomycota*, *Mucoromycota*, *Neocallmastigomycota*, other Fungi, *Metazoa*, *Viridiplantae*, and all other phyla (based on EUKARYOME annotations). Numbers of ASVs within each taxonomic group shown in parentheses. Asterisks denote p < 0.001 in pairwise comparisons, followed by a letter in parentheses indicating the database providing greater resolution (**C**).

**Figure S2.**
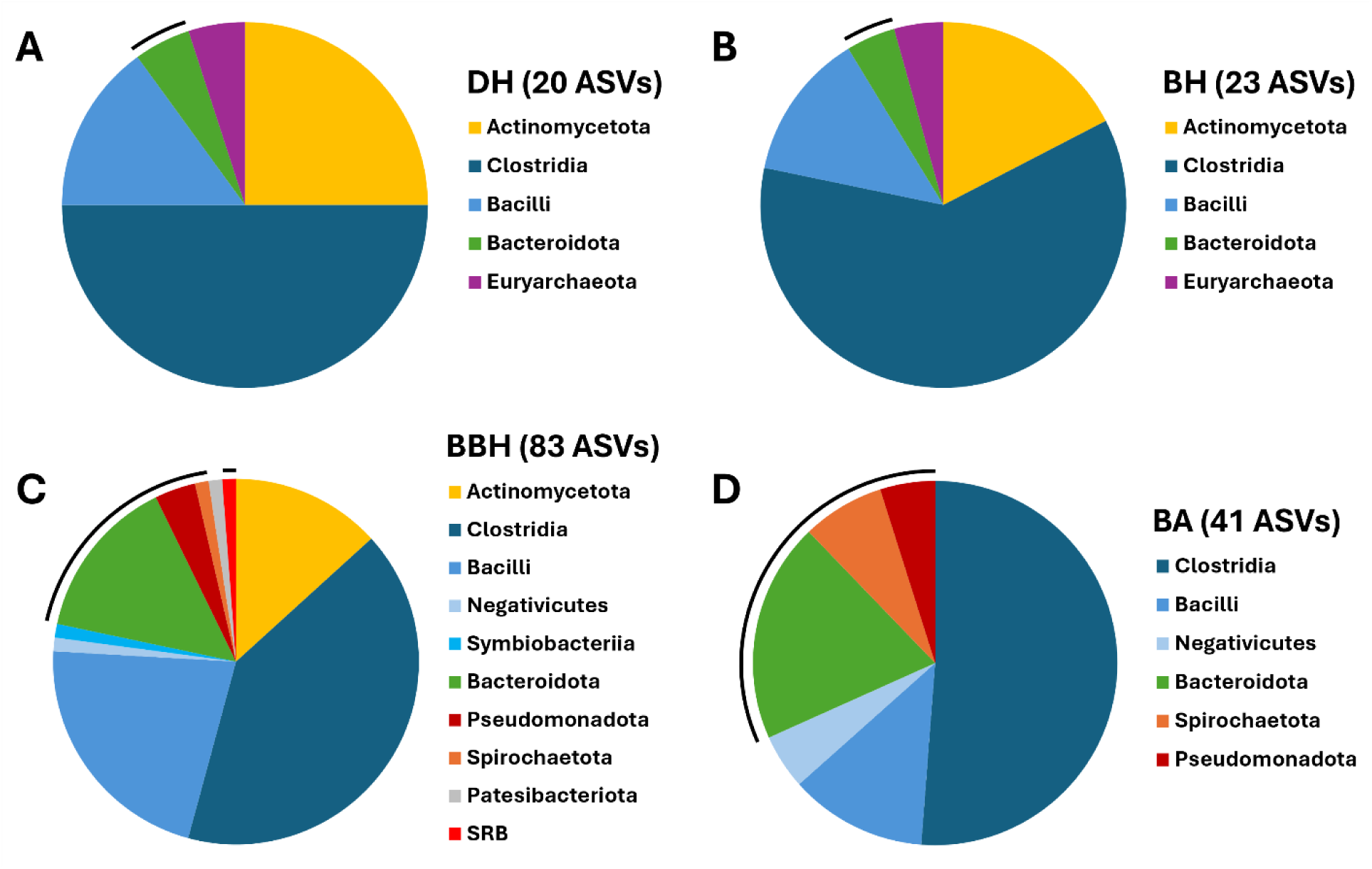
Pie charts showing the number and phylum-level assignments of 16S rRNA amplicon sequence variants (ASVs) enriched in feces from dairy heifers (DH, *n* = 8; **A**), beef heifers (BH, *n* = 8; **B**), bred beef heifers (BBH, *n* = 7; **C**), and adult beef cows (BA, *n* = 7, **D**). Black arcs indicate gram-negative phyla. See **Table S4** for details.

**Figure S3.**
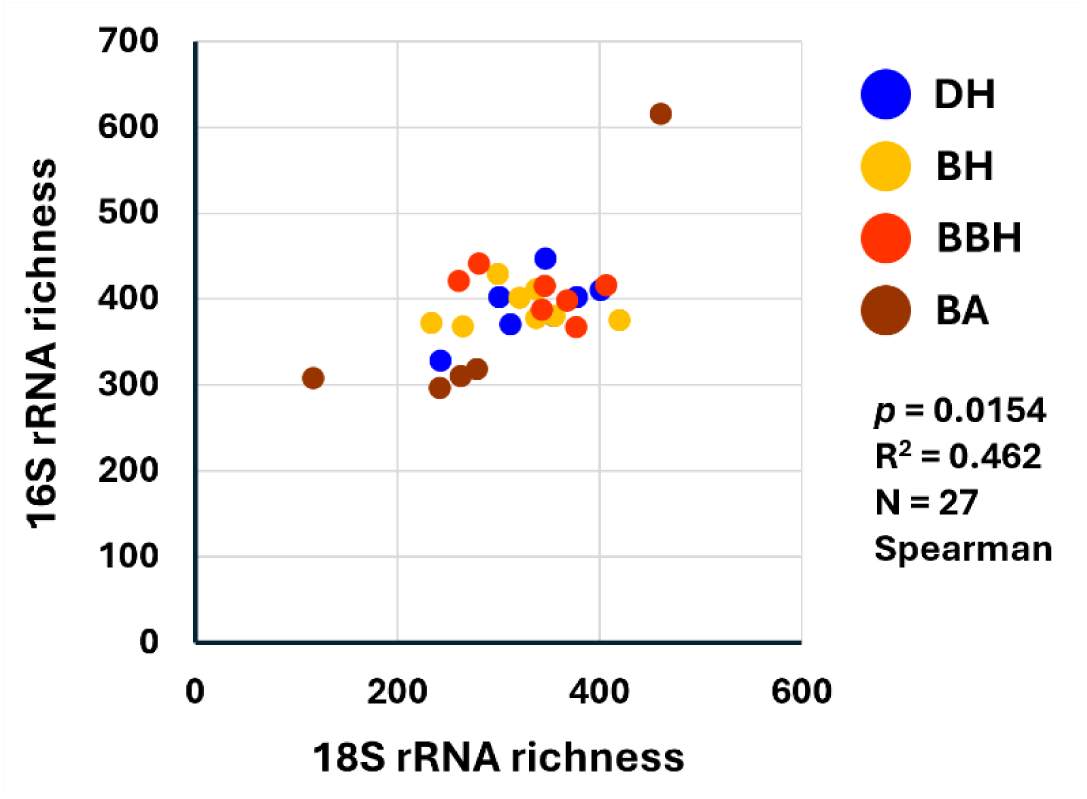
Dot plot showing the relationship between detected 18S rRNA and 16S rRNA richness in fecal samples from dairy heifers (DH, *n* = 8), beef heifers (BH, *n* = 8), bred beef heifers (BBH, *n* = 7), and adult beef cows (BA, *n* = 7). Legend at right, followed by results of Spearman’s rank order correlation test.

## References

1. Henderson G, Cox F, Ganesh S, et al. Rumen microbial community composition varies with diet and host, but a core microbiome is found across a wide geographical range. Sci Rep 2015;5:14567.

2. Shanks OC, Kelty CA, Archibeque S, et al. Community structures of fecal bacteria in cattle from different animal feeding operations. Appl Environ Microbiol 2011;77:2992–3001.

3. Conteville LC, da Silva JV, Andrade BGN, et al. Rumen and fecal microbiomes are related to diet and production traits in Bos indicus beef cattle. Front Microbiol 2023;14:1282851.

4. Brulin L, Ducrocq S, Estelle J, et al. The fecal microbiota of Holstein cows is heritable and genetically correlated to dairy performances. J Dairy Sci 2024;107:11254–11268.

5. Choi J, Park JS. Comparative analyses of the V4 and V9 regions of 18S rDNA for the extant eukaryotic community using the Illumina platform. Sci Rep 2020;10:6519.

6. Jamy M, Foster R, Barbera P, et al. Long-read metabarcoding of the eukaryotic rDNA operon to phylogenetically and taxonomically resolve environmental diversity. Mol Ecol Resour 2020;20:429–443.

7. Wang C, Chen X, Lei Y, et al. Optimizing primer selection for microbial diversity studies in freshwater lakes: a comprehensive evaluation of 16S rRNA, 18S rRNA, and ITS gene regions. FEMS Microbiol Ecol 2026;102.

8. Hestetun JT, Lanzen A, Kongsrud JA, et al. Metabarcoding and targeted barcoding can enhance Norwegian Continental Shelf macrofauna species inventories. PeerJ 2026;14:e20849.

9. Hadziavdic K, Lekang K, Lanzen A, et al. Characterization of the 18S rRNA gene for designing universal eukaryote specific primers. PLoS One 2014;9:e87624.

10. Caporaso JG, Lauber CL, Walters WA, et al. Global patterns of 16S rRNA diversity at a depth of millions of sequences per sample. Proc Natl Acad Sci U S A 2011;108 Suppl 1:4516–22.

11. Walters WA, Caporaso JG, Lauber CL, et al. PrimerProspector: de novo design and taxonomic analysis of barcoded polymerase chain reaction primers. Bioinformatics 2011;27:1159–61.

12. Bolyen E, Rideout JR, Dillon MR, et al. Reproducible, interactive, scalable and extensible microbiome data science using QIIME 2. Nat Biotechnol 2019;37:852– 857.

13. Martin M. Cutadapt removes adapter sequences from high-throughput sequencing reads. EMBnet.journal 2011;17:10–12.

14. Callahan BJ, McMurdie PJ, Rosen MJ, et al. DADA2: High-resolution sample inference from Illumina amplicon data. Nat Methods 2016;13:581–3.

15. Chuvochina M, Gerken J, Frentrup M, et al. SILVA in 2026: a global core biodata resource for rRNA within the DSMZ digital diversity. Nucleic Acids Res 2026;54:D334– D341.

16. Tedersoo L, Hosseyni Moghaddam MS, Mikryukov V, et al. EUKARYOME: the rRNA gene reference database for identification of all eukaryotes. Database (Oxford) 2024;2024.

17. Bokulich NA, Kaehler BD, Rideout JR, et al. Optimizing taxonomic classification of marker-gene amplicon sequences with QIIME 2’s q2-feature-classifier plugin. Microbiome 2018;6:90.

18. Lu Y, Zhou G, Ewald J, et al. MicrobiomeAnalyst 2.0: comprehensive statistical, functional and integrative analysis of microbiome data. Nucleic Acids Res 2023;51:W310–W318.

19. Benjamini Y HY. Controlling the false discovery rate: a practical and powerful approach to multiple testing. Journal of the Royal Statistical Society Series B 1995;57:289–300.

20. Ogawa S, Shibahara T, Sano A, et al. Generalized hyperkeratosis caused by Scopulariopsis brevicaulis in a Japanese Black calf. J Comp Pathol 2008;138:145–50.

21. Macura AB, Skora M. 21-year retrospective study of the prevalence of Scopulariopsis brevicaulis in patients suspected of superficial mycoses. Postepy Dermatol Alergol 2015;32:189–94.

22. Zajc J, Gunde-Cimerman N. The Genus Wallemia-From Contamination of Food to Health Threat. Microorganisms 2018;6.

23. Wu Y, Jiao C, Diao Q, et al. Effect of Dietary and Age Changes on Ruminal Microbial Diversity in Holstein Calves. Microorganisms 2023;12.

24. Amin N, Schwarzkopf S, Troscher-Mussotter J, et al. Host metabolome and faecal microbiome shows potential interactions impacted by age and weaning times in calves. Anim Microbiome 2023;5:12.

25. Furman O, Shenhav L, Sasson G, et al. Stochasticity constrained by deterministic effects of diet and age drive rumen microbiome assembly dynamics. Nat Commun 2020;11:1904.

26. Fan P, Nelson CD, Driver JD, et al. Host genetics exerts lifelong effects upon hindgut microbiota and its association with bovine growth and immunity. ISME J 2021;15:2306–2321.

27. Lourenco JM, Krause TR, Welch CB, et al. Longitudinal Changes of the Ruminal Microbiota in Angus Beef Steers. Animals (Basel) 2022;12.

28. Barden M, Richards-Rios P, Ganda E, et al. Maternal influences on oral and faecal microbiota maturation in neonatal calves in beef and dairy production systems. Anim Microbiome 2020;2:31.

29. Noyes NR, Yang X, Linke LM, et al. Characterization of the resistome in manure, soil and wastewater from dairy and beef production systems. Sci Rep 2016;6:24645.

30. Rovira P, McAllister T, Lakin SM, et al. Characterization of the Microbial Resistome in Conventional and “Raised Without Antibiotics” Beef and Dairy Production Systems. Front Microbiol 2019;10:1980.

31. Wang W, Wei X, Wu L, et al. The occurrence of antibiotic resistance genes in the microbiota of yak, beef and dairy cattle characterized by a metagenomic approach. J Antibiot (Tokyo) 2021;74:508–518.

32. Bailoni L, Carraro L, Cardin M, et al. Active Rumen Bacterial and Protozoal Communities Revealed by RNA-Based Amplicon Sequencing on Dairy Cows Fed Different Diets at Three Physiological Stages. Microorganisms 2021;9.

33. Quilcate C, Estrada R, Romero Y, et al. Changes in Gut Microbiota in Peruvian Cattle Genetic Nucleus by Breed and Correlations with Beef Quality. Vet Sci 2024;11.

34. Rojas D, Estrada R, Romero Y, et al. Sex-Induced Changes in Microbial Eukaryotes and Prokaryotes in Gastrointestinal Tract of Simmental Cattle. Biology (Basel) 2024;13.

35. Estrada R, Romero Y, Quilcate C, et al. Age-Dependent Changes in Protist and Fungal Microbiota in a Peruvian Cattle Genetic Nucleus. Life (Basel) 2024;14.

36. Lopez-Garcia A, Saborio-Montero A, Gutierrez-Rivas M, et al. Fungal and ciliate protozoa are the main rumen microbes associated with methane emissions in dairy cattle. Gigascience 2022;11.

37. Saborio-Montero A, Gutierrez-Rivas M, Goiri I, et al. Rumen eukaryotes are the main phenotypic risk factors for larger methane emissions in dairy cattle. Livestock Science 2022;263.

38. Guyader J, Eugene M, Noziere P, et al. Influence of rumen protozoa on methane emission in ruminants: a meta-analysis approach. Animal 2014;8:1816–25.

39. Whitelaw FG, Eadie JM, Bruce LA, et al. Methane formation in faunated and ciliate-free cattle and its relationship with rumen volatile fatty acid proportions. Br J Nutr 1984;52:261–75.

40. Schonhusen U, Zitnan R, Kuhla S, et al. Effects of protozoa on methane production in rumen and hindgut of calves around time of weaning. Arch Tierernahr 2003;57:279– 95.

41. Liu C, Meng Q, Chen Y, et al. Role of Age-Related Shifts in Rumen Bacteria and Methanogens in Methane Production in Cattle. Front Microbiol 2017;8:1563.

42. Avramenko RW, Redman EM, Lewis R, et al. The use of nemabiome metabarcoding to explore gastro-intestinal nematode species diversity and anthelmintic treatment effectiveness in beef calves. Int J Parasitol 2017;47:893–902.

43. Ramirez AL, Herrera G, Munoz M, et al. Describing the intestinal microbiota of Holstein Fasciola-positive and -negative cattle from a hyperendemic area of fascioliasis in central Colombia. PLoS Negl Trop Dis 2021;15:e0009658.

44. Meili CH, TagElDein MA, Jones AL, et al. Diversity and community structure of anaerobic gut fungi in the rumen of wild and domesticated herbivores. Appl Environ Microbiol 2024;90:e0149223.

45. Liggenstoffer AS, Youssef NH, Couger MB, et al. Phylogenetic diversity and community structure of anaerobic gut fungi (phylum Neocallimastigomycota) in ruminant and non-ruminant herbivores. ISME J 2010;4:1225–35.

46. Suthar VS, Burfeind O, Bonk S, et al. Endogenous and exogenous progesterone influence body temperature in dairy cows. J Dairy Sci 2012;95:2381–9.

47. Lappalainen S, Pasanen AL, Reiman M, et al. Serum IgG antibodies against Wallemia sebi and Fusarium species in Finnish farmers. Ann Allergy Asthma Immunol 1998;81:585–92.

48. Dickie HA, Rankin J. Farmer’s lung; an acute granulomatous interstitial pneumonitis occurring in agricultural workers. J Am Med Assoc 1958;167:1069–76.

49. Kornilova OA, Skazina MA, Belokon ME, et al. New findings of ciliates Buxtonella sulcata in the faeces of cattle from northern regions of Russia and comparative analysis of Buxtonella species. Vet Parasitol Reg Stud Reports 2025;63:101314.

50. Dianso JA, Garcia GG, Belotindos LP, et al. Molecular identification of Buxtonella sulcata from associated-diarrhea in water buffaloes (Bubalus bubalis) in the Philippines. Ann Parasitol 2018;64:93–100.

51. Ganai A, Parveen S, Kaur D, et al. Incidence of Buxtonella sulcata in bovines in R.S. Pura, Jammu. J Parasit Dis 2015;39:446–7.

52. Hasheminasab SS, Moradi P, Talvar HM, et al. Buxtonella spp. like infection in cattle in Sanandaj province, Iran. Ann Parasitol 2015;61:247–51.

53. Hong KO, Youn HJ. [Incidence of Buxtonella sulcata from cattle in Kyonggi-do]. Korean J Parasitol 1995;33:135–8.

54. Maurya PS, Ramakant, Singh A, et al. Prevalence, molecular characterization, and therapeutic management of Buxtonella sulcata in buffaloes from Western Uttar Pradesh, India. Vet Parasitol Reg Stud Reports 2025;65:101342.

55. Fox MT, Jacobs DE. Patterns of infection with Buxtonella sulcata in British cattle. Res Vet Sci 1986;41:90–2.

56. Henriksen SA. [Buxtonella sulcata, an intestinal ciliate of apparently frequent occurrence in Danish cattle (author’s transl)]. Nord Vet Med 1977;29:452–7.

57. Norman Grim J, Jirku-Pomajbikova K, Ponce-Gordo F. Light microscopic morphometrics, ultrastructure, and molecular phylogeny of the putative pycnotrichid Ciliate, Buxtonella sulcata. Eur J Protistol 2015;51:425–36.

58. Ribeiro M, Gomes-Goncalves S, Silva A, et al. Screening for the ciliate Buxtonella sulcata in free-ranging dairy cattle on Terceira Island, Azores archipelago. Parasite 2025;32:21.

59. Munoz-Zanzi C, Honores-Perez P, Godoy-Alfaro C, et al. First morphological and molecular description of the ciliate protozoa Buxtonella sulcata (Ciliophora, Vestibuliferida) in cattle in Chile. Vet Parasitol Reg Stud Reports 2025;57:101183.

60. El-Ashram S, Aboelhadid SM, Kamel AA, et al. Diversity of Parasitic Diarrhea Associated with Buxtonella Sulcata in Cattle and Buffalo Calves with Control of Buxtonellosis. Animals (Basel) 2019;9.

61. Rees CW. Studies on the morphology and behaviour of Buxtonella sulcata from cattle and Balantidium coli from the pig. Parasitology 1930;22:314–325.

62. Urman H, Kelley G. Buxtonella sulcata a ciliate associated with ulcerative colitis in a cow and prevalence of infection in Nebraska cattle. Iowa State University Veterinarian 1963;26.

63. Frey CF, Muller N, Stauber N, et al. Simplicimonas-like DNA in vaginal swabs of cows and heifers cross-reacting in the real-time PCR for T. foetus. Vet Parasitol 2017;237:30–36.

