## Supplementary material for "Multi-domain analysis of the fecal microbiome of dairy heifers, beef heifers, bred beef heifers, and adult beef cows in the central United States using paired 16S rRNA and 18S rRNA metabarcoding": Table S1

**Table S1. Details related to the heifers and cows providing samples for the current study**

| Sample_ID | Type | Age (yrs) | Status | DOB | Days bred |
|---|---|---|---|---|---|
| Sample 01 | dairy | 0.5 | Heifer | 8/15/2025 | 0 |
| Sample 02 | dairy | 0.5 | Heifer | 8/30/2025 | 0 |
| Sample 03 | dairy | 0.5 | Heifer | 8/18/2025 | 0 |
| Sample 04 | dairy | 0.5 | Heifer | 9/2/2025 | 0 |
| Sample 05 | dairy | 0.5 | Heifer | 9/2/2025 | 0 |
| Sample 06 | dairy | 0.5 | Heifer | 8/25/2025 | 0 |
| Sample 07 | dairy | 0.5 | Heifer | 8/29/2025 | 0 |
| Sample 08 | dairy | 0.4 | Heifer | 9/9/2025 | 0 |
| Sample 09 | beef | 1.5 | Bred heifer | 8/30/2024 | 80 |
| Sample 10 | beef | 1.5 | Bred heifer | 8/30/2024 | 60 |
| Sample 11 | beef | 1.5 | Bred heifer | 8/25/2024 | 60 |
| Sample 12 | beef | 1.4 | Bred heifer | 10/6/2024 | open |
| Sample 13 | beef | 1.4 | Bred heifer | 9/21/2024 | 80 |
| Sample 14 | beef | 1.4 | Bred heifer | 9/27/2024 | 80 |
| Sample 15 | beef | 1.4 | Bred heifer | 9/7/2024 | 60 |
| Sample 16 | beef | 1.4 | Bred heifer | 9/24/2024 | 70 |
| Sample 17 | beef | 0.4 | Heifer | 9/10/2025 | 0 |
| Sample 18 | beef | 0.4 | Heifer | 9/6/2025 | 0 |
| Sample 19 | beef | 0.4 | Heifer | 10/2/2025 | 0 |
| Sample 20 | beef | 0.4 | Heifer | 9/5/2025 | 0 |
| Sample 21 | beef | 0.5 | Heifer | 8/19/2025 | 0 |
| Sample 22 | beef | 0.5 | Heifer | 8/25/2025 | 0 |
| Sample 23 | beef | 0.4 | Heifer | 9/7/2025 | 0 |
| Sample 24 | beef | 0.4 | Heifer | 9/17/2025 | 0 |
| Sample 25 | beef | 6 | cow | 2019 | 60 |
| Sample 26 | beef | 8 | cow | 2017 | 80 |
| Sample 27 | beef | 11 | cow | 2014 | 80 |
| Sample 28 | beef | 8 | cow | 2017 | 80 |
| Sample 29 | beef | 13 | cow | 2012 | 70 |
| Sample 30 | beef | 7 | cow | 2018 | 80 |
| Sample 31 | beef | 9 | cow | 2016 | 20 |
| Sample 32 | beef | 9 | cow | 2016 | 70 |
