## Supplementary material for "Multi-domain analysis of the fecal microbiome of dairy heifers, beef heifers, bred beef heifers, and adult beef cows in the central United States using paired 16S rRNA and 18S rRNA metabarcoding": Table S2

**Table S2. Results of one-way permutational multivariate analysis of variance (PERMANOVA) using unweighted (Jaccard) and weighted (Bray-Curtis) dissimilarities**

### 18S rRNA (eukaryotic) sequences

**S2A. PERMANOVA using unweighted (Jaccard) dissimilarities**

| | |
|---|---|
| Permutation N: | 9999 |
| Total sum of squares: | 13.02 |
| Within-group sum of squares: | 10.9 |
| F: | 1.752 |
| p (same): | 0.0001 |

**Results of pairwise comparisons**

| p values | DH | BH | BBH | BA |
|---|---|---|---|---|
| DH | | 0.0012 | 0.0007 | 0.0002 |
| BH | 0.0012 | | 0.0568 | 0.0003 |
| BBH | 0.0007 | 0.0568 | | 0.0004 |
| BA | 0.0002 | 0.0003 | 0.0004 | |

| F values | DH | BH | BBH | BA |
|---|---|---|---|---|
| DH | | 1.302 | 1.376 | 2.287 |
| BH | 1.302 | | 1.172 | 2.349 |
| BBH | 1.376 | 1.172 | | 2.077 |
| BA | 2.287 | 2.349 | 2.077 | |

**S2B. PERMANOVA using weighted (Bray-Curtis) dissimilarities**

| | |
|---|---|
| Permutation N: | 9999 |
| Total sum of squares: | 8.462 |
| Within-group sum of squares: | 5.316 |
| F: | 5.326 |
| p (same): | 0.0001 |

**Results of pairwise comparisons**

| p values | DH | BH | BBH | BA |
|---|---|---|---|---|
| DH | | 0.0378 | 0.0062 | 0.0003 |
| BH | 0.0378 | | 0.0562 | 0.0001 |
| BBH | 0.0062 | 0.0562 | | 0.0003 |
| BA | 0.0003 | 0.0001 | 0.0003 | |

| F values | DH | BH | BBH | BA |
|---|---|---|---|---|
| DH | | 1.868 | 2.253 | 7.586 |
| BH | 1.868 | | 2.07 | 9.006 |
| BBH | 2.253 | 2.07 | | 5.714 |
| BA | 7.586 | 9.006 | 5.714 | |

### 16S rRNA (prokaryotic) sequences

**S2C. PERMANOVA using unweighted (Jaccard) dissimilarities**

| | |
|---|---|
| Permutation N: | 9999 |
| Total sum of squares: | 3.14 |
| Within-group sum of squares: | 1.97 |
| F: | 5.151 |
| p (same): | 0.0001 |

**Results of pairwise comparisons**

| p values | DH | BH | BBH | BA |
|---|---|---|---|---|
| DH | | 0.0004 | 0.0003 | 0.0005 |
| BH | 0.0004 | | 0.0003 | 0.0001 |
| BBH | 0.0003 | 0.0003 | | 0.0007 |
| BA | 0.0005 | 0.0001 | 0.0007 | |

| F values | DH | BH | BBH | BA |
|---|---|---|---|---|
| DH | | 4.541 | 4.08 | 5.331 |
| BH | 4.541 | | 2.707 | 6.35 |
| BBH | 4.08 | 2.707 | | 6.568 |
| BA | 5.331 | 6.35 | 6.568 | |

**S2D. PERMANOVA using weighted (Bray-Curtis) dissimilarities**

| | |
|---|---|
| Permutation N: | 9999 |
| Total sum of squares: | 1.938 |
| Within-group sum of squares: | 1.108 |
| F: | 6.488 |
| p (same): | 0.0001 |

**Results of pairwise comparisons**

| p values | DH | BH | BBH | BA |
|---|---|---|---|---|
| DH | | 0.0001 | 0.0002 | 0.0001 |
| BH | 0.0001 | | 0.0006 | 0.0004 |
| BBH | 0.0002 | 0.0006 | | 0.0005 |
| BA | 0.0001 | 0.0004 | 0.0005 | |

| F values | DH | BH | BBH | BA |
|---|---|---|---|---|
| DH | | 4.813 | 5.671 | 5.963 |
| BH | 4.813 | | 4.566 | 8.658 |
| BBH | 5.671 | 4.566 | | 8.524 |
| BA | 5.963 | 8.658 | 8.524 | |
