## Supplementary material for "Multi-domain analysis of the fecal microbiome of dairy heifers, beef heifers, bred beef heifers, and adult beef cows in the central United States using paired 16S rRNA and 18S rRNA metabarcoding": Table S3

**Table S3.** Results of differential abundance testing of 18S rRNA (eukaryotic) amplicon sequence variants via analysis of variance (ANOVA) followed by pairwise comparisons using Fisher's least significant difference test.

| ASV_ID | taxonomy_EUKARYOME | F value | p value (raw) | p value (FDR-adj) | Fisher's LSD |
|---|---|---|---|---|---|
| ASV0047 | k__Amoebozoa;p__Evosea;c__Archamoebea;o__Entamoebida;f__Entamoebidae;g__Entamoeba | 10.423 | 0.0001242 | 0.028022 | BA - BBH (p=7.2e-05); BA - BH (p=7.2e-05); BA - DH (p=0.000103) |
| ASV0060 | k__Amoebozoa;p__Evosea;c__Archamoebea;o__Entamoebida;f__Entamoebidae;g__Entamoeba;s__bovis | 16.262 | 4.53E-06 | 0.0016617 | BA - BBH (p=2e-06); BA - BH (p=5e-06); BA - DH (p=6e-06) |
| ASV0192 | k__Amoebozoa;p__Evosea;c__Archamoebea;o__Entamoebida;f__Entamoebidae;g__Entamoeba;s__bovis | 9.7108 | 0.00019876 | 0.038863 | BA - BBH (p=8e-05); BA - BH (p=0.000239); BA - DH (p=0.000114) |
| ASV0090 | k__Fungi;p__Ascomycota;c__Eurotiomycetes;o__Eurotiales | 11.278 | 7.23E-05 | 0.017329 | BBH - BA (p=7e-06); BH - BA (p=0.0106); DH - BA (p=0.000484); BBH - BH (p=0.00476) |
| ASV0006 | k__Fungi;p__Ascomycota;c__Eurotiomycetes;o__Eurotiales;f__Aspergillaceae;g__Aspergillus | 19.019 | 1.23E-06 | 0.00060043 | BBH - BA (p=1e-06); BH - BA (p=1.1e-05); DH - BA (p<1e-06) |
| ASV0043 | k__Fungi;p__Ascomycota;c__Eurotiomycetes;o__Eurotiales;f__Aspergillaceae;g__Aspergillus | 72.449 | 1.82E-12 | 1.07E-08 | BBH - BA (p<1e-06); BH - BA (p=0.0289); DH - BA (p<1e-06); BBH - BH (p<1e-06); DH - BH (p<1e-06) |
| ASV0049 | k__Fungi;p__Ascomycota;c__Eurotiomycetes;o__Eurotiales;f__Aspergillaceae;g__Aspergillus | 23.488 | 1.90E-07 | 0.00027039 | BBH - BA (p=2e-06); DH - BA (p=3e-06); BBH - BH (p=4e-06); DH - BH (p=7e-06) |
| ASV0097 | k__Fungi;p__Ascomycota;c__Eurotiomycetes;o__Eurotiales;f__Aspergillaceae;g__Aspergillus | 55.946 | 3.13E-11 | 9.17E-08 | BBH - BA (p<1e-06); DH - BA (p<1e-06); BBH - BH (p<1e-06); DH - BH (p<1e-06) |
| ASV0052 | k__Fungi;p__Ascomycota;c__Eurotiomycetes;o__Eurotiales;f__Aspergillaceae;g__Penicillium | 19.815 | 8.63E-07 | 0.00056265 | BBH - BA (p=1e-06); BH - BA (p=0.000128); DH - BA (p<1e-06); BBH - BH (p=0.0307); DH - BH (p=0.012) |
| ASV0005 | k__Fungi;p__Ascomycota;c__Eurotiomycetes;o__Eurotiales;f__Trichocomaceae;g__Talaromyces | 33.033 | 7.43E-09 | 1.45E-05 | BBH - BA (p<1e-06); BH - BA (p=0.00018); DH - BA (p<1e-06); BBH - BH (p=8.9e-05); DH - BH (p=0.000184) |
| ASV0001 | k__Fungi;p__Ascomycota;c__Eurotiomycetes;o__Eurotiales;f__Trichocomaceae;g__Thermomyces;s__lanuginosus | 17.646 | 2.31E-06 | 0.00090491 | BBH - BA (p=1e-06); BH - BA (p=3.6e-05); DH - BA (p=1e-06) |
| ASV0002 | k__Fungi;p__Ascomycota;c__Eurotiomycetes;o__Eurotiales;f__Trichocomaceae;g__Thermomyces;s__lanuginosus | 18.696 | 1.42E-06 | 0.00064161 | BBH - BA (p<1e-06); BH - BA (p=3.9e-05); DH - BA (p=1e-06) |
| ASV0007 | k__Fungi;p__Ascomycota;c__Eurotiomycetes;o__Eurotiales;f__Trichocomaceae;g__Thermomyces;s__lanuginosus | 18.053 | 1.91E-06 | 0.00080107 | BBH - BA (p=1e-06); BH - BA (p=4.4e-05); DH - BA (p=1e-06) |
| ASV0012 | k__Fungi;p__Ascomycota;c__Eurotiomycetes;o__Eurotiales;f__Trichocomaceae;g__Thermomyces;s__lanuginosus | 19.317 | 1.08E-06 | 0.00057347 | BBH - BA (p<1e-06); BH - BA (p=1.6e-05); DH - BA (p=1e-06) |
| ASV0032 | k__Fungi;p__Ascomycota;c__Eurotiomycetes;o__Eurotiales;f__Trichocomaceae;g__Thermomyces;s__lanuginosus | 12.328 | 3.83E-05 | 0.0097602 | BBH - BA (p=2.4e-05); BH - BA (p=0.000185); DH - BA (p=1.1e-05) |
| ASV0039 | k__Fungi;p__Ascomycota;c__Eurotiomycetes;o__Eurotiales;f__Trichocomaceae;g__Thermomyces;s__lanuginosus | 19.554 | 9.68E-07 | 0.00056775 | BBH - BA (p<1e-06); BH - BA (p=2.6e-05); DH - BA (p=1e-06) |
| ASV0059 | k__Fungi;p__Ascomycota;c__Eurotiomycetes;o__Eurotiales;f__Trichocomaceae;g__Thermomyces;s__lanuginosus | 9.957 | 0.00016862 | 0.036151 | BBH - BA (p=2.5e-05); BH - BA (p=0.00228); DH - BA (p=0.000192) |
| ASV0061 | k__Fungi;p__Ascomycota;c__Eurotiomycetes;o__Eurotiales;f__Trichocomaceae;g__Thermomyces;s__lanuginosus | 15.382 | 7.09E-06 | 0.002445 | BBH - BA (p=1e-06); BH - BA (p=4.2e-05); DH - BA (p=1.1e-05) |
| ASV0062 | k__Fungi;p__Ascomycota;c__Eurotiomycetes;o__Eurotiales;f__Trichocomaceae;g__Thermomyces;s__lanuginosus | 13.088 | 2.46E-05 | 0.0065791 | BBH - BA (p=3.2e-05); DH - BA (p=0.0138); BBH - BH (p=1.1e-05); BBH - DH (p=0.0224); DH - BH (p=0.00869) |
| ASV0078 | k__Fungi;p__Ascomycota;c__Eurotiomycetes;o__Eurotiales;f__Trichocomaceae;g__Thermomyces;s__lanuginosus | 14.413 | 1.18E-05 | 0.0036467 | BBH - BA (p=3e-06); BH - BA (p=0.00042); DH - BA (p=7e-06) |
| ASV0104 | k__Fungi;p__Ascomycota;c__Eurotiomycetes;o__Eurotiales;f__Trichocomaceae;g__Thermomyces;s__lanuginosus | 9.7937 | 0.00018801 | 0.03803 | BBH - BA (p=3.1e-05); BH - BA (p=0.00136); DH - BA (p=0.000175) |
| ASV0045 | k__Fungi;p__Ascomycota;c__Sordariomycetes;o__Hypocreales;f__Hypocreales.fam.incertae.sedis;g__Bulbithecium;s__pinkertoniae | 22.941 | 2.36E-07 | 0.00027039 | BBH - BA (p<1e-06); BH - BA (p=0.000113); DH - BA (p<1e-06); BBH - BH (p=0.00401); DH - BH (p=0.0106) |
| ASV0057 | k__Fungi;p__Ascomycota;c__Sordariomycetes;o__Microascales;f__Microascaceae;g__Microascus;s__cirrosus | 13.717 | 1.73E-05 | 0.0050688 | BBH - BA (p=1e-05); BH - BA (p=6.9e-05); DH - BA (p=7e-06) |
| ASV0063 | k__Fungi;p__Ascomycota;c__Sordariomycetes;o__Sordariales;f__Chaetomiaceae;g__Chaetomium | 20.439 | 6.59E-07 | 0.00052225 | BBH - BA (p=1e-06); BH - BA (p=1.2e-05); DH - BA (p<1e-06) |
| ASV0004 | k__Fungi;p__Basidiomycota;c__Tritirachiomycetes;o__Tritirachiales;f__Tritirachiaceae;g__Tritirachium | 20.258 | 7.12E-07 | 0.00052225 | BBH - BA (p=1e-06); BH - BA (p=0.00318); DH - BA (p=1e-06); BBH - BH (p=0.00112); DH - BH (p=0.00107) |
| ASV0009 | k__Fungi;p__Basidiomycota;c__Tritirachiomycetes;o__Tritirachiales;f__Tritirachiaceae;g__Tritirachium | 22.539 | 2.77E-07 | 0.00027039 | BBH - BA (p<1e-06); BH - BA (p=0.000694); DH - BA (p<1e-06); BBH - BH (p=0.000769); DH - BH (p=0.00368) |
| ASV0144 | k__Fungi;p__Basidiomycota;c__Tritirachiomycetes;o__Tritirachiales;f__Tritirachiaceae;g__Tritirachium | 13.084 | 2.47E-05 | 0.0065791 | BBH - BA (p=3.4e-05); BH - BA (p=0.00578); DH - BA (p=5e-06); BBH - BH (p=0.039); DH - BH (p=0.00564) |
| ASV0203 | k__Fungi;p__Basidiomycota;c__Tritirachiomycetes;o__Tritirachiales;f__Tritirachiaceae;g__Tritirachium | 11.243 | 7.39E-05 | 0.017329 | BBH - BA (p=0.000909); DH - BA (p=7.7e-05); BBH - BH (p=0.00267); DH - BH (p=0.000191) |
| ASV0231 | k__Fungi;p__Neocallimastigomycota;c__Neocallimastigomycetes;o__Neocallimastigales;f__Neocallimastigaceae;g__Neocallimastix;s__frontalis | 9.5112 | 0.00022744 | 0.043037 | BA - BBH (p=6.8e-05); BA - BH (p=0.000327); BA - DH (p=0.000159) |
| ASV0082 | k__Metazoa;p__Nematoda;c__Chromadorea;o__Strongylida;f__Haemonchidae | 9.9223 | 0.00017256 | 0.036151 | DH - BA (p=0.000332); DH - BBH (p=0.000102); DH - BH (p=0.000102) |
| ASV0123 | k__Metazoa;p__Nematoda;c__Chromadorea;o__Strongylida;f__Haemonchidae | 14.844 | 9.39E-06 | 0.0030587 | DH - BA (p=1.3e-05); DH - BBH (p=2.9e-05); DH - BH (p=5e-06) |
| ASV0442 | k__Metazoa;p__Nematoda;c__Chromadorea;o__Strongylida;f__Haemonchidae | 9.4334 | 0.00023981 | 0.043961 | DH - BA (p=0.000337); DH - BBH (p=0.000151); DH - BH (p=0.000151) |
