## Supplementary material for "Multi-domain analysis of the fecal microbiome of dairy heifers, beef heifers, bred beef heifers, and adult beef cows in the central United States using paired 16S rRNA and 18S rRNA metabarcoding": Table S4

**Table S4.** Results of differential abundance testing of 16S rRNA (prokaryotic) amplicon sequence variants via analysis of variance (ANOVA) followed by pairwise comparisons using Fisher's least significant difference test.

| ASV_ID | taxonomy | F value | p value (raw) | p value (FDR-a | Fisher's LSD |
|---|---|---|---|---|---|
| ASV0037 | d__Archaea;p__Euryarchaeota;c__Methanobacteria;o__Methanobacteriales;f__Methanobacteriaceae | 5.1847 | 0.0060944 | 0.045282 | DH - BA (p=0.00345); DH - BBH (p=0.00635); DH - BH (p=0.0026) |
| ASV0035 | d__Archaea;p__Euryarchaeota;c__Methanobacteria;o__Methanobacteriales;f__Methanobacteriaceae;g__Methanobrevibacter | 18.624 | 1.17E-06 | 6.37E-05 | BH - BA (p<1e-06); DH - BA (p=0.00816); BH - BBH (p=1.3e-05); BH - DH (p=0.000172) |
| ASV0886 | d__Bacteria;p__Actinobacteriota;c__Actinobacteria;o__Bifidobacteriales;f__Bifidobacteriaceae | 21.044 | 3.96E-07 | 2.62E-05 | BH - BA (p=1e-06); BH - BBH (p=8e-06); BH - DH (p<1e-06) |
| ASV0068 | d__Bacteria;p__Actinobacteriota;c__Actinobacteria;o__Bifidobacteriales;f__Bifidobacteriaceae;g__Aeriscardovia | 13.481 | 1.68E-05 | 0.00052915 | DH - BA (p=3.9e-05); DH - BBH (p=1.8e-05); DH - BH (p=2e-05) |
| ASV0882 | d__Bacteria;p__Actinobacteriota;c__Actinobacteria;o__Bifidobacteriales;f__Bifidobacteriaceae;g__Bifidobacterium | 8.5899 | 0.00039513 | 0.0063496 | BBH - BA (p=6.3e-05); BH - BA (p=0.00495); DH - BA (p=0.000519) |
| ASV0920 | d__Bacteria;p__Actinobacteriota;c__Actinobacteria;o__Pseudonocardiales;f__Pseudonocardiaceae;g__Saccharopolyspora | 6.622 | 0.0017961 | 0.019495 | BBH - BA (p=0.0466); DH - BA (p=0.000617); DH - BH (p=0.00118) |
| ASV0928 | d__Bacteria;p__Actinobacteriota;c__Actinobacteria;o__Pseudonocardiales;f__Pseudonocardiaceae;g__Saccharopolyspora;s__Saccharopolyspora_rectivirgula | 5.1783 | 0.006129 | 0.045282 | BBH - BA (p=0.00129); BBH - BH (p=0.00408) |
| ASV0629 | d__Bacteria;p__Actinobacteriota;c__Actinobacteria;o__Streptomycetales;f__Streptomycetaceae;g__Streptomyces | 10.039 | 0.00014314 | 0.0029467 | BBH - BA (p=1.6e-05); BH - BA (p=0.0267); DH - BA (p=0.000973); BBH - BH (p=0.00451) |
| ASV0855 | d__Bacteria;p__Actinobacteriota;c__Actinobacteria;o__Streptosporangiales;f__Nocardiopsaceae;g__Thermobifida;s__Thermobifida_fusca | 47.816 | 1.01E-10 | 4.03E-08 | BBH - BA (p<1e-06); BH - BA (p=1e-06); DH - BA (p<1e-06); BBH - BH (p=1.8e-05); BBH - DH (p=0.00625); DH - BH (p=0.0266) |
| ASV0647 | d__Bacteria;p__Actinobacteriota;c__Actinobacteria;o__Streptosporangiales;f__Streptosporangiaceae;g__Thermopolyspora | 25.716 | 6.07E-08 | 6.04E-06 | BBH - BA (p=1e-06); DH - BA (p<1e-06); BBH - BH (p=2.9e-05); DH - BH (p=3e-06) |
| ASV0660 | d__Bacteria;p__Actinobacteriota;c__Actinobacteria;o__Streptosporangiales;f__Thermomonosporaceae;g__Thermobispora | 13.763 | 1.43E-05 | 0.00047543 | BBH - BA (p=0.000444); BH - BA (p=0.00261); DH - BA (p=1e-06); DH - BBH (p=0.0365); DH - BH (p=0.00425) |
| ASV0661 | d__Bacteria;p__Actinobacteriota;c__Actinobacteria;o__Streptosporangiales;f__Thermomonosporaceae;g__Thermobispora;s__Thermobispora_bispora | 20.011 | 6.22E-07 | 3.91E-05 | BBH - BA (p=1e-06); BH - BA (p=0.0133); DH - BA (p=1e-06); BBH - BH (p=0.000392); DH - BH (p=0.000665) |
| ASV0542 | d__Bacteria;p__Actinobacteriota;c__Coriobacteriia;o__Coriobacteriales | 13.024 | 2.20E-05 | 0.00063926 | BBH - BA (p=0.0106); BH - BA (p=0.00604); DH - BA (p=1e-06); DH - BBH (p=0.0023); DH - BH (p=0.00249) |
| ASV0808 | d__Bacteria;p__Actinobacteriota;c__Coriobacteriia;o__Coriobacteriales;f__Atopobiaceae | 7.0745 | 0.0012488 | 0.014296 | BH - BA (p=0.00147); BBH - DH (p=0.0231); BH - DH (p=0.000366) |
| ASV0702 | d__Bacteria;p__Actinobacteriota;c__Coriobacteriia;o__Coriobacteriales;f__Atopobiaceae;g__Atopobium | 10.287 | 0.00012122 | 0.0025393 | BBH - BA (p=0.0216); BH - BA (p=8e-06); DH - BA (p=0.00494); BH - BBH (p=0.00586); BH - DH (p=0.0174) |
| ASV0700 | d__Bacteria;p__Actinobacteriota;c__Coriobacteriia;o__Coriobacteriales;f__Atopobiaceae;g__Olsenella | 7.0236 | 0.0013003 | 0.014646 | DH - BA (p=0.00102); DH - BBH (p=0.0011); DH - BH (p=0.000894) |
| ASV1173 | d__Bacteria;p__Actinobacteriota;c__Coriobacteriia;o__Coriobacteriales;f__Coriobacteriales_Incertae_Sedis;g__Raoultibacter | 5.5246 | 0.0045205 | 0.03551 | DH - BA (p=0.00176); DH - BBH (p=0.00176); DH - BH (p=0.0272) |
| ASV0046 | d__Bacteria;p__Actinobacteriota;c__Coriobacteriia;o__Coriobacteriales;f__Eggerthellaceae | 13.818 | 1.39E-05 | 0.00047418 | BH - BA (p=0.000431); DH - BA (p=4e-06); BH - BBH (p=0.013); DH - BBH (p=0.000134) |
| ASV0047 | d__Bacteria;p__Actinobacteriota;c__Coriobacteriia;o__Coriobacteriales;f__Eggerthellaceae;g__DNF00809 | 10.91 | 8.05E-05 | 0.0018493 | BBH - BA (p=0.0142); BH - BA (p=6e-06); DH - BA (p=0.00132); BH - BBH (p=0.00695); BH - DH (p=0.0435) |
| ASV0063 | d__Bacteria;p__Actinobacteriota;c__Coriobacteriia;o__Coriobacteriales;f__Eggerthellaceae;g__DNF00809 | 6.1673 | 0.0026134 | 0.024765 | BH - BA (p=0.00112); BH - DH (p=0.00134) |
| ASV0062 | d__Bacteria;p__Actinobacteriota;c__Coriobacteriia;o__Coriobacteriales;f__Eggerthellaceae;g__Enterorhabdus | 8.1996 | 0.00052655 | 0.0079582 | DH - BA (p=0.000179); DH - BBH (p=0.00059); DH - BH (p=0.00141) |
| ASV0585 | d__Bacteria;p__Actinobacteriota;c__Coriobacteriia;o__Coriobacteriales;f__Eggerthellaceae;g__Slackia | 5.9252 | 0.0032045 | 0.027929 | DH - BA (p=0.00219); DH - BBH (p=0.00219); DH - BH (p=0.00212) |
| ASV0296 | d__Bacteria;p__Bacteroidota;c__Bacteroidia;o__Bacteroidales;f__Bacteroidaceae;g__Bacteroides | 33.649 | 4.20E-09 | 7.17E-07 | BA - BBH (p=2e-06); BA - BH (p<1e-06); BA - DH (p<1e-06); BBH - BH (p=0.0363); BBH - DH (p=0.00681) |
| ASV0292 | d__Bacteria;p__Bacteroidota;c__Bacteroidia;o__Bacteroidales;f__Bacteroidaceae;g__Bacteroides | 7.9705 | 0.00062504 | 0.0085591 | BA - BBH (p=0.00389); BA - BH (p=0.000746); BA - DH (p=9.8e-05) |
| ASV0376 | d__Bacteria;p__Bacteroidota;c__Bacteroidia;o__Bacteroidales;f__Bacteroidaceae;g__Bacteroides | 6.4043 | 0.0021466 | 0.021906 | BA - BBH (p=0.00946); BA - BH (p=0.0394); BA - DH (p=0.000206); BH - DH (p=0.0354) |
| ASV0399 | d__Bacteria;p__Bacteroidota;c__Bacteroidia;o__Bacteroidales;f__Bacteroidales_RF16_group | 40.998 | 5.33E-10 | 1.06E-07 | BBH - BA (p<1e-06); BH - BA (p=0.000136); BBH - BH (p=1.9e-05); BBH - DH (p<1e-06); BH - DH (p=9e-05) |
| ASV0370 | d__Bacteria;p__Bacteroidota;c__Bacteroidia;o__Bacteroidales;f__Bacteroidales_RF16_group | 5.6827 | 0.0039424 | 0.032242 | BBH - BA (p=0.00517); BH - BA (p=0.00433); BBH - DH (p=0.0121); BH - DH (p=0.0104) |
| ASV0390 | d__Bacteria;p__Bacteroidota;c__Bacteroidia;o__Bacteroidales;f__Bacteroidales_RF16_group | 5.0562 | 0.0068339 | 0.049155 | BBH - BA (p=0.00231); BBH - DH (p=0.00233) |
| ASV0324 | d__Bacteria;p__Bacteroidota;c__Bacteroidia;o__Bacteroidales;f__Dysgonomonadaceae | 6.2072 | 0.0025278 | 0.024308 | BBH - BA (p=0.00171); BBH - BH (p=0.00128); BBH - DH (p=0.00139) |
| ASV0366 | d__Bacteria;p__Bacteroidota;c__Bacteroidia;o__Bacteroidales;f__F082 | 5.5295 | 0.0045014 | 0.03551 | BH - BA (p=0.00369); BBH - DH (p=0.0376); BH - DH (p=0.00159) |
| ASV0380 | d__Bacteria;p__Bacteroidota;c__Bacteroidia;o__Bacteroidales;f__p-251-o5 | 6.0087 | 0.0029858 | 0.026454 | BA - BH (p=0.03); BA - DH (p=0.0181); BBH - BH (p=0.00274); BBH - DH (p=0.00154) |
| ASV0360 | d__Bacteria;p__Bacteroidota;c__Bacteroidia;o__Bacteroidales;f__p-251-o5 | 5.6667 | 0.0039974 | 0.032469 | BBH - BA (p=0.043); BBH - BH (p=0.0408); BBH - DH (p=0.000339) |
| ASV0369 | d__Bacteria;p__Bacteroidota;c__Bacteroidia;o__Bacteroidales;f__Paludibacteraceae | 10.688 | 9.30E-05 | 0.0020197 | BBH - BA (p=5.5e-05); BH - BA (p=0.0415); BBH - BH (p=0.00898); BBH - DH (p=3.6e-05); BH - DH (p=0.0353) |
| ASV0358 | d__Bacteria;p__Bacteroidota;c__Bacteroidia;o__Bacteroidales;f__Paludibacteraceae;g__Paludibacter | 21.581 | 3.14E-07 | 2.49E-05 | BBH - BA (p<1e-06); BH - BA (p=0.00602); BBH - BH (p=5.3e-05); BBH - DH (p=1e-06) |
| ASV0317 | d__Bacteria;p__Bacteroidota;c__Bacteroidia;o__Bacteroidales;f__Prevotellaceae | 8.4146 | 0.00044918 | 0.0070569 | BBH - BA (p=0.0029); BH - BA (p=0.00111); BBH - DH (p=0.00221); BH - DH (p=0.000796) |
| ASV0308 | d__Bacteria;p__Bacteroidota;c__Bacteroidia;o__Bacteroidales;f__Prevotellaceae;g__Alloprevotella | 6.1313 | 0.0026933 | 0.024928 | BBH - BA (p=0.00738); BH - BA (p=0.00119); BBH - DH (p=0.0324); BH - DH (p=0.00589) |
| ASV0288 | d__Bacteria;p__Bacteroidota;c__Bacteroidia;o__Bacteroidales;f__Prevotellaceae;g__Prevotellaceae_Ga6A1_group | 6.2577 | 0.0024236 | 0.023526 | BA - BBH (p=0.00156); BA - BH (p=0.000765); BA - DH (p=0.00274) |
| ASV0291 | d__Bacteria;p__Bacteroidota;c__Bacteroidia;o__Bacteroidales;f__Prevotellaceae;g__Prevotellaceae_UCG-003 | 46.196 | 1.47E-10 | 4.40E-08 | BBH - BA (p<1e-06); BH - BA (p=3e-06); BBH - BH (p=0.000255); BBH - DH (p<1e-06); BH - DH (p=3e-06) |
| ASV0289 | d__Bacteria;p__Bacteroidota;c__Bacteroidia;o__Bacteroidales;f__Prevotellaceae;g__Prevotellaceae_UCG-003 | 5.0365 | 0.0069561 | 0.049734 | BBH - DH (p=0.00334); BH - DH (p=0.00238) |
| ASV0329 | d__Bacteria;p__Bacteroidota;c__Bacteroidia;o__Bacteroidales;f__Rikenellaceae;g__Alistipes | 9.333 | 0.00023251 | 0.004271 | BA - BBH (p=0.000165); BA - BH (p=0.000168); BA - DH (p=0.000206) |
| ASV0361 | d__Bacteria;p__Bacteroidota;c__Bacteroidia;o__Bacteroidales;f__Rikenellaceae;g__Rikenella | 7.9583 | 0.00063082 | 0.0085591 | DH - BA (p=0.00061); DH - BBH (p=0.00061); DH - BH (p=0.000427) |
| ASV0367 | d__Bacteria;p__Bacteroidota;c__Bacteroidia;o__Bacteroidales;f__Rikenellaceae;g__Rikenellaceae_RC9_gut_group | 13.375 | 1.79E-05 | 0.00054803 | BBH - BA (p=1.4e-05); BBH - BH (p=7.6e-05); BBH - DH (p=9e-06) |
| ASV0373 | d__Bacteria;p__Bacteroidota;c__Bacteroidia;o__Bacteroidales;f__Rikenellaceae;g__Rikenellaceae_RC9_gut_group | 11.149 | 6.91E-05 | 0.0018326 | BA - DH (p=0.00125); BBH - DH (p=6.2e-05); BH - DH (p=2.4e-05) |
| ASV0396 | d__Bacteria;p__Bacteroidota;c__Bacteroidia;o__Bacteroidales;f__Tannerellaceae;g__Parabacteroides | 16.707 | 2.98E-06 | 0.00013452 | BA - BBH (p=0.0017); BA - BH (p=0.0145); BA - DH (p<1e-06); BBH - DH (p=0.00254); BH - DH (p=0.000131) |
| ASV0443 | d__Bacteria;p__Desulfobacterota;c__Desulfovibrionia;o__Desulfovibrionales;f__Desulfovibrionaceae;g__Mailhella | 12.285 | 3.41E-05 | 0.00094632 | BBH - BA (p=6e-06); BBH - BH (p=0.000201); BBH - DH (p=8e-05) |
| ASV0538 | d__Bacteria;p__Firmicutes;c__Bacilli;o__Acholeplasmatales;f__Acholeplasmataceae;g__Anaeroplasma | 11.115 | 7.06E-05 | 0.0018326 | BA - BBH (p=0.000448); BA - BH (p=8.3e-05); BA - DH (p=1.6e-05) |
| ASV0539 | d__Bacteria;p__Firmicutes;c__Bacilli;o__Acholeplasmatales;f__Acholeplasmataceae;g__EMP-G18 | 12.918 | 2.34E-05 | 0.00066384 | BA - BBH (p=1.6e-05); BA - BH (p=5.5e-05); BA - DH (p=1.7e-05) |
| ASV1041 | d__Bacteria;p__Firmicutes;c__Bacilli;o__Bacillales;f__Bacillaceae;g__Aeribacillus | 9.7712 | 0.0001717 | 0.0034169 | BBH - BA (p=3.7e-05); BH - BA (p=0.00268); DH - BA (p=0.000158) |
| ASV1033 | d__Bacteria;p__Firmicutes;c__Bacilli;o__Bacillales;f__Bacillaceae;g__Bacillus;s__Bacillus_coagulans | 9.565 | 0.00019786 | 0.0038103 | BBH - BA (p=0.02); BH - BA (p=1.4e-05); DH - BA (p=0.00912); BH - BBH (p=0.00982); BH - DH (p=0.0146) |
| ASV1038 | d__Bacteria;p__Firmicutes;c__Bacilli;o__Bacillales;f__Bacillaceae;g__Bacillus;s__Bacillus_smithii | 8.577 | 0.00039885 | 0.0063496 | BBH - BA (p=0.000131); BH - BA (p=0.042); DH - BA (p=0.000327); BBH - BH (p=0.0194); DH - BH (p=0.0487) |
| ASV1094 | d__Bacteria;p__Firmicutes;c__Bacilli;o__Bacillales;f__Bacillaceae;g__Bacillus;s__Bacillus_thermoamylovorans | 9.6992 | 0.00018039 | 0.0035309 | BH - BA (p=3.4e-05); BH - BBH (p=0.0029); BH - DH (p=0.000307) |
| ASV0020 | d__Bacteria;p__Firmicutes;c__Bacilli;o__Bacillales;f__Bacillaceae;g__Caldibacillus | 31.29 | 8.80E-09 | 1.17E-06 | BBH - BA (p<1e-06); BH - BA (p=0.00563); DH - BA (p<1e-06); BBH - BH (p=1.9e-05); DH - BH (p=2.3e-05) |
| ASV0039 | d__Bacteria;p__Firmicutes;c__Bacilli;o__Bacillales;f__Bacillaceae;g__Geobacillus | 43.089 | 3.13E-10 | 7.47E-08 | BBH - BA (p<1e-06); BH - BA (p<1e-06); DH - BA (p<1e-06); BBH - BH (p=0.000716); DH - BH (p=0.018) |
| ASV1042 | d__Bacteria;p__Firmicutes;c__Bacilli;o__Bacillales;f__Planococcaceae;g__Rummeliibacillus | 26.981 | 3.82E-08 | 4.14E-06 | BH - BA (p<1e-06); BH - BBH (p<1e-06); BH - DH (p<1e-06) |
| ASV0832 | d__Bacteria;p__Firmicutes;c__Bacilli;o__Bacillales;f__Sporolactobacillaceae;g__Tuberibacillus;s__Caenibacillus_caldisaponilyticus | 16.664 | 3.04E-06 | 0.00013452 | BBH - BA (p<1e-06); BH - BA (p=0.00175); DH - BA (p=3.4e-05); BBH - BH (p=0.00147) |
| ASV1031 | d__Bacteria;p__Firmicutes;c__Bacilli;o__Brevibacillales;f__Brevibacillaceae;g__Brevibacillus;s__Brevibacillus_limnophilus | 5.6226 | 0.0041526 | 0.033276 | BBH - BA (p=0.00171); BBH - BH (p=0.00128); BBH - DH (p=0.045) |
| ASV0797 | d__Bacteria;p__Firmicutes;c__Bacilli;o__Brevibacillales;f__Brevibacillaceae;g__Brevibacillus;s__Brevibacillus_thermoruber | 10.999 | 7.60E-05 | 0.0018493 | BBH - BA (p=6.2e-05); BH - BA (p=0.0131); DH - BA (p=2.7e-05); BBH - BH (p=0.0322); DH - BH (p=0.0189) |
| ASV0973 | d__Bacteria;p__Firmicutes;c__Bacilli;o__Erysipelotrichales;f__Erysipelatoclostridiaceae | 9.3521 | 0.00022942 | 0.004271 | BA - BBH (p=0.000465); BA - BH (p=0.00882); BA - DH (p=3.2e-05); BH - DH (p=0.0321) |
| ASV1120 | d__Bacteria;p__Firmicutes;c__Bacilli;o__Erysipelotrichales;f__Erysipelatoclostridiaceae;g__Erysipelatoclostridium | 8.0185 | 0.00060285 | 0.0084683 | BBH - BA (p=0.000193); BH - BA (p=0.00162); BBH - DH (p=0.00397); BH - DH (p=0.0314) |
| ASV1121 | d__Bacteria;p__Firmicutes;c__Bacilli;o__Erysipelotrichales;f__Erysipelatoclostridiaceae;g__UCG-004 | 6.5184 | 0.0019544 | 0.020835 | BA - BBH (p=0.00235); BA - BH (p=0.00579); BA - DH (p=0.000311) |
| ASV0961 | d__Bacteria;p__Firmicutes;c__Bacilli;o__Erysipelotrichales;f__Erysipelatoclostridiaceae;g__UCG-004 | 6.1991 | 0.0025448 | 0.024308 | BBH - BA (p=0.00126); BBH - DH (p=0.000933) |
| ASV1003 | d__Bacteria;p__Firmicutes;c__Bacilli;o__Erysipelotrichales;f__Erysipelotrichaceae | 60.512 | 7.37E-12 | 4.40E-09 | BBH - BA (p<1e-06); BH - BA (p<1e-06); BBH - DH (p<1e-06); BH - DH (p<1e-06) |
| ASV0955 | d__Bacteria;p__Firmicutes;c__Bacilli;o__Erysipelotrichales;f__Erysipelotrichaceae;g__[Anaerorhabdus]_furcosa_group | 16.165 | 3.92E-06 | 0.00016725 | BBH - BA (p<1e-06); BH - BA (p=2.7e-05); DH - BA (p=0.000487); BBH - DH (p=0.00686) |
| ASV0953 | d__Bacteria;p__Firmicutes;c__Bacilli;o__Erysipelotrichales;f__Erysipelotrichaceae;g__Solobacterium | 5.3413 | 0.0053064 | 0.041142 | BH - BBH (p=0.00149); BH - DH (p=0.00286) |
| ASV1123 | d__Bacteria;p__Firmicutes;c__Bacilli;o__Erysipelotrichales;f__Erysipelotrichaceae;g__Solobacterium | 5.2718 | 0.0056418 | 0.043181 | BBH - BA (p=0.017); BH - BA (p=0.0042); DH - BA (p=0.000954) |
| ASV1098 | d__Bacteria;p__Firmicutes;c__Bacilli;o__Lactobacillales;f__Aerococcaceae;g__Aerococcus;s__Aerococcus_vaginalis | 5.1756 | 0.0061438 | 0.045282 | BBH - BA (p=0.00333); BBH - BH (p=0.00379); BBH - DH (p=0.00256) |
| ASV1090 | d__Bacteria;p__Firmicutes;c__Bacilli;o__Lactobacillales;f__Lactobacillaceae;g__Lactobacillus | 13.182 | 2.00E-05 | 0.00059757 | DH - BA (p=3.3e-05); DH - BBH (p=3.2e-05); DH - BH (p=2e-05) |
| ASV1095 | d__Bacteria;p__Firmicutes;c__Bacilli;o__Lactobacillales;f__Lactobacillaceae;g__Ligilactobacillus;s__Lactobacillus_ruminis | 30.407 | 1.17E-08 | 1.40E-06 | BA - BBH (p<1e-06); BA - BH (p<1e-06); BA - DH (p<1e-06) |
| ASV0998 | d__Bacteria;p__Firmicutes;c__Bacilli;o__Lactobacillales;f__Lactobacillaceae;g__Limosilactobacillus | 6.138 | 0.0026783 | 0.024928 | DH - BA (p=0.00206); DH - BBH (p=0.00206); DH - BH (p=0.00152) |
| ASV0999 | d__Bacteria;p__Firmicutes;c__Bacilli;o__Lactobacillales;f__Lactobacillaceae;g__Limosilactobacillus;s__Lactobacillus_mucosae | 5.8491 | 0.0034187 | 0.029157 | DH - BA (p=0.00349); DH - BBH (p=0.00221); DH - BH (p=0.00163) |
| ASV0833 | d__Bacteria;p__Firmicutes;c__Bacilli;o__Paenibacillales;f__Paenibacillaceae;g__Thermobacillus | 9.0484 | 0.00028416 | 0.0049896 | BBH - BA (p=0.0161); DH - BA (p=0.000169); BBH - BH (p=0.0219); DH - BH (p=0.000205) |
| ASV0836 | d__Bacteria;p__Firmicutes;c__Bacilli;o__Paenibacillales;f__Paenibacillaceae;g__Thermobacillus | 8.0464 | 0.00059035 | 0.0083915 | BBH - BA (p=0.000177); DH - BA (p=0.000857); BBH - BH (p=0.00917); DH - BH (p=0.0419) |
| ASV0837 | d__Bacteria;p__Firmicutes;c__Bacilli;o__Paenibacillales;f__Paenibacillaceae;g__Thermobacillus;s__Thermobacillus_composti | 23.135 | 1.65E-07 | 1.42E-05 | BBH - BA (p<1e-06); BH - BA (p=6.6e-05); DH - BA (p<1e-06); BBH - BH (p=0.0153); DH - BH (p=0.0113) |
| ASV0806 | d__Bacteria;p__Firmicutes;c__Bacilli;o__Thermoactinomycetales;f__Thermoactinomycetaceae;g__Thermoactinomyces | 13.714 | 1.47E-05 | 0.00047566 | BBH - BA (p=3.9e-05); DH - BA (p=1e-05); BBH - BH (p=0.00296); DH - BH (p=0.000829) |
| ASV0566 | d__Bacteria;p__Firmicutes;c__Clostridia;o__Christensenellales;f__Christensenellaceae | 6.7904 | 0.001567 | 0.017165 | BBH - BA (p=0.00133); BH - BA (p=0.00239); BBH - DH (p=0.0063); BH - DH (p=0.0116) |
| ASV0839 | d__Bacteria;p__Firmicutes;c__Clostridia;o__Christensenellales;f__Christensenellaceae;g__Christensenellaceae_R-7_group;s__bacterium_YE57 | 19.356 | 8.36E-07 | 4.76E-05 | DH - BA (p=3e-06); DH - BBH (p=5e-06); DH - BH (p<1e-06) |

| | | | | | |
|---|---|---|---|---|---|
| ASV1010 | d__Bacteria;p__Firmicutes;c__Clostridia;o__Clostridia_vadinBB60_group | 21.399 | 3.40E-07 | 2.49E-05 | BBH - BA (p=2.6e-05); BH - BA (p=2e-06); BBH - DH (p=2.8e-05); BH - DH (p=2e-06) |
| ASV0331 | d__Bacteria;p__Firmicutes;c__Clostridia;o__Clostridia_vadinBB60_group | 6.4759 | 0.0020237 | 0.021196 | BA - BBH (p=0.0208); BA - BH (p=0.00116); BA - DH (p=0.000427) |
| ASV0940 | d__Bacteria;p__Firmicutes;c__Clostridia;o__Clostridiales;f__Clostridiaceae;g__Clostridium_sensu_stricto_1 | 17.365 | 2.15E-06 | 0.00010255 | BA - BBH (p=4e-06); BA - BH (p=1e-06); BA - DH (p=1.7e-05) |
| ASV0783 | d__Bacteria;p__Firmicutes;c__Clostridia;o__Eubacteriales;f__Anaerofustaceae;g__Anaerofustis | 6.4132 | 0.0021309 | 0.021906 | BBH - BA (p=0.00052); BH - BA (p=0.00302); DH - BA (p=0.00182) |
| ASV0663 | d__Bacteria;p__Firmicutes;c__Clostridia;o__Eubacteriales;f__Eubacteriaceae;g__Eubacterium | 18.131 | 1.48E-06 | 7.37E-05 | BBH - BA (p=6.6e-05); BH - BA (p=3e-06); BBH - DH (p=0.000169); BH - DH (p=8e-06) |
| ASV0705 | d__Bacteria;p__Firmicutes;c__Clostridia;o__Lachnospirales;f__Lachnospiraceae | 8.6087 | 0.00038976 | 0.0063496 | BBH - BA (p=0.00019); BH - BA (p=0.00452); DH - BA (p=0.000125) |
| ASV0500 | d__Bacteria;p__Firmicutes;c__Clostridia;o__Lachnospirales;f__Lachnospiraceae | 5.1659 | 0.0061969 | 0.045393 | BA - BH (p=0.0336); DH - BBH (p=0.00777); DH - BH (p=0.00151) |
| ASV0745 | d__Bacteria;p__Firmicutes;c__Clostridia;o__Lachnospirales;f__Lachnospiraceae;g__[Eubacterium]_fissicatena_group | 19.52 | 7.76E-07 | 4.64E-05 | DH - BA (p=1e-06); DH - BBH (p=3e-06); DH - BH (p=2e-06) |
| ASV1171 | d__Bacteria;p__Firmicutes;c__Clostridia;o__Lachnospirales;f__Lachnospiraceae;g__[Eubacterium]_hallii_group | 10.699 | 9.23E-05 | 0.0020197 | BH - BA (p=3.5e-05); BH - BBH (p=7.4e-05); BH - DH (p=0.0043) |
| ASV1170 | d__Bacteria;p__Firmicutes;c__Clostridia;o__Lachnospirales;f__Lachnospiraceae;g__[Eubacterium]_hallii_group | 9.067 | 0.00028043 | 0.0049896 | BBH - BA (p=0.000809); BH - BA (p=4.3e-05); DH - BA (p=0.00077) |
| ASV1176 | d__Bacteria;p__Firmicutes;c__Clostridia;o__Lachnospirales;f__Lachnospiraceae;g__[Eubacterium]_ventriosum_group | 6.3612 | 0.0022244 | 0.022208 | DH - BA (p=0.00106); DH - BBH (p=0.0122); DH - BH (p=0.000761) |
| ASV0733 | d__Bacteria;p__Firmicutes;c__Clostridia;o__Lachnospirales;f__Lachnospiraceae;g__[Ruminococcus]_gauvreauii_group | 18.48 | 1.26E-06 | 6.52E-05 | BH - BA (p<1e-06); BH - BBH (p=4.9e-05); BH - DH (p=6e-06) |
| ASV0708 | d__Bacteria;p__Firmicutes;c__Clostridia;o__Lachnospirales;f__Lachnospiraceae;g__[Ruminococcus]_gauvreauii_group | 15.962 | 4.35E-06 | 0.0001793 | BBH - BA (p=0.00197); BH - BA (p=1e-06); BH - BBH (p=0.00573); BH - DH (p=3.7e-05) |
| ASV0731 | d__Bacteria;p__Firmicutes;c__Clostridia;o__Lachnospirales;f__Lachnospiraceae;g__[Ruminococcus]_gauvreauii_group | 7.9613 | 0.00062939 | 0.0085591 | DH - BA (p=0.000609); DH - BBH (p=0.000609); DH - BH (p=0.000426) |
| ASV1188 | d__Bacteria;p__Firmicutes;c__Clostridia;o__Lachnospirales;f__Lachnospiraceae;g__[Ruminococcus]_torques_group | 8.2734 | 0.00049849 | 0.0077298 | BA - BBH (p=0.000281); BA - BH (p=0.00107); BA - DH (p=0.000196) |
| ASV0681 | d__Bacteria;p__Firmicutes;c__Clostridia;o__Lachnospirales;f__Lachnospiraceae;g__Blautia | 6.5454 | 0.0019118 | 0.020565 | BH - BA (p=0.0314); BBH - DH (p=0.00287); BH - DH (p=0.00038) |
| ASV0766 | d__Bacteria;p__Firmicutes;c__Clostridia;o__Lachnospirales;f__Lachnospiraceae;g__Coprococcus | 15.522 | 5.48E-06 | 0.00021809 | BBH - BA (p=0.000125); BH - BA (p=1e-06); DH - BA (p=4.9e-05) |
| ASV1186 | d__Bacteria;p__Firmicutes;c__Clostridia;o__Lachnospirales;f__Lachnospiraceae;g__Coprococcus | 8.2343 | 0.00051316 | 0.0078553 | DH - BA (p=0.000457); DH - BBH (p=0.00401); DH - BH (p=0.000137) |
| ASV0751 | d__Bacteria;p__Firmicutes;c__Clostridia;o__Lachnospirales;f__Lachnospiraceae;g__Dorea | 7.8975 | 0.00066042 | 0.0087615 | BA - BBH (p=0.000348); BA - BH (p=0.000244); BA - DH (p=0.00158) |
| ASV0763 | d__Bacteria;p__Firmicutes;c__Clostridia;o__Lachnospirales;f__Lachnospiraceae;g__Frisingicoccus | 32.27 | 6.44E-09 | 9.61E-07 | BBH - BA (p<1e-06); BH - BA (p=1e-05); BBH - BH (p=0.0371); BBH - DH (p<1e-06); BH - DH (p=2e-06) |
| ASV0764 | d__Bacteria;p__Firmicutes;c__Clostridia;o__Lachnospirales;f__Lachnospiraceae;g__Frisingicoccus | 7.7269 | 0.00075182 | 0.0096524 | DH - BA (p=0.00648); DH - BBH (p=0.000504); DH - BH (p=0.000225) |
| ASV0765 | d__Bacteria;p__Firmicutes;c__Clostridia;o__Lachnospirales;f__Lachnospiraceae;g__GCA-900066575 | 9.378 | 0.0002253 | 0.00427 | BA - BBH (p=0.000924); BA - BH (p=7.9e-05); BA - DH (p=0.000126) |
| ASV1174 | d__Bacteria;p__Firmicutes;c__Clostridia;o__Lachnospirales;f__Lachnospiraceae;g__Howardella | 7.669 | 0.00078587 | 0.0098772 | BBH - BA (p=0.00098); BH - BA (p=0.000229); DH - BA (p=0.00079) |
| ASV1167 | d__Bacteria;p__Firmicutes;c__Clostridia;o__Lachnospirales;f__Lachnospiraceae;g__Lachnospira | 7.7317 | 0.00074907 | 0.0096524 | BA - BBH (p=0.000623); BA - BH (p=0.000447); BA - DH (p=0.000447) |
| ASV0707 | d__Bacteria;p__Firmicutes;c__Clostridia;o__Lachnospirales;f__Lachnospiraceae;g__Lachnospiraceae_NK3A20_group | 8.5837 | 0.00039693 | 0.0063496 | BBH - BA (p=0.0143); BH - BA (p=4.3e-05); BH - BBH (p=0.0375); BH - DH (p=0.00213) |
| ASV0706 | d__Bacteria;p__Firmicutes;c__Clostridia;o__Lachnospirales;f__Lachnospiraceae;g__Lachnospiraceae_NK3A20_group | 6.8919 | 0.0014443 | 0.016117 | BH - BA (p=0.000226); DH - BA (p=0.00192); BH - BBH (p=0.0404) |
| ASV0713 | d__Bacteria;p__Firmicutes;c__Clostridia;o__Lachnospirales;f__Lachnospiraceae;g__Lachnospiraceae_NK3A20_group | 5.1908 | 0.0060615 | 0.045282 | BBH - BA (p=0.0187); BH - BA (p=0.000879); BH - DH (p=0.0211) |
| ASV0739 | d__Bacteria;p__Firmicutes;c__Clostridia;o__Lachnospirales;f__Lachnospiraceae;g__Lachnospiraceae_NK4A136_group | 14.807 | 8.03E-06 | 0.00029072 | BA - BBH (p=0.00102); BA - BH (p=7e-06); BA - DH (p=2e-06); BBH - DH (p=0.0369) |
| ASV0743 | d__Bacteria;p__Firmicutes;c__Clostridia;o__Lachnospirales;f__Lachnospiraceae;g__Lachnospiraceae_UCG-002 | 7.1582 | 0.0011688 | 0.01355 | BH - BA (p=0.000405); DH - BA (p=0.0268); BH - BBH (p=0.00104) |
| ASV0715 | d__Bacteria;p__Firmicutes;c__Clostridia;o__Lachnospirales;f__Lachnospiraceae;g__Lachnospiraceae_UCG-002; | 7.5793 | 0.00084193 | 0.010364 | BBH - BA (p=0.00716); BH - BA (p=8.8e-05); DH - BA (p=0.0022) |
| ASV0762 | d__Bacteria;p__Firmicutes;c__Clostridia;o__Lachnospirales;f__Lachnospiraceae;g__Lachnospiraceae_UCG-008 | 5.683 | 0.0039416 | 0.032242 | BH - BA (p=0.00147); BH - DH (p=0.00168) |
| ASV0717 | d__Bacteria;p__Firmicutes;c__Clostridia;o__Lachnospirales;f__Lachnospiraceae;g__Marvinbryantia | 6.0621 | 0.0028544 | 0.025819 | BA - BBH (p=0.0243); BA - BH (p=0.00736); DH - BBH (p=0.00628); DH - BH (p=0.00157) |
| ASV0683 | d__Bacteria;p__Firmicutes;c__Clostridia;o__Lachnospirales;f__Lachnospiraceae;g__Tuzzerella | 6.064 | 0.0028499 | 0.025819 | BBH - BA (p=0.03); BH - BA (p=0.00446); BBH - DH (p=0.0115); BH - DH (p=0.00137) |
| ASV0571 | d__Bacteria;p__Firmicutes;c__Clostridia;o__Monoglobales;f__Monoglobaceae;g__Monoglobus | 6.8333 | 0.0015139 | 0.016737 | BH - BA (p=0.00237); BH - BBH (p=0.0131); BH - DH (p=0.000232) |
| ASV0948 | d__Bacteria;p__Firmicutes;c__Clostridia;o__Monoglobales;f__Monoglobaceae;g__Monoglobus | 6.357 | 0.0022319 | 0.022208 | BBH - BA (p=0.0065); BH - BA (p=0.000712); BBH - DH (p=0.0477); BH - DH (p=0.00646) |
| ASV1079 | d__Bacteria;p__Firmicutes;c__Clostridia;o__Monoglobales;f__Monoglobaceae;g__Monoglobus | 5.8377 | 0.0034521 | 0.029232 | BH - BA (p=0.00121); DH - BA (p=0.00422); BH - BBH (p=0.0172) |
| ASV0980 | d__Bacteria;p__Firmicutes;c__Clostridia;o__Oscillospirales;f__[Eubacterium]_coprostanoligenes_group;g__[Eubacterium]_coprostanoligenes_group | 8.0572 | 0.00058561 | 0.0083915 | BBH - BA (p=0.000694); BH - BA (p=0.0086); BBH - DH (p=0.000488); BH - DH (p=0.00662) |
| ASV0985 | d__Bacteria;p__Firmicutes;c__Clostridia;o__Oscillospirales;f__Oscillospiraceae | 7.1693 | 0.0011586 | 0.01355 | BBH - BA (p=0.00243); BH - BA (p=0.0322); BBH - DH (p=0.000415); BH - DH (p=0.0068) |
| ASV0991 | d__Bacteria;p__Firmicutes;c__Clostridia;o__Oscillospirales;f__Oscillospiraceae;g__Colidextribacter | 11.276 | 6.37E-05 | 0.0017288 | BBH - BA (p=1.9e-05); BH - BA (p=0.00116); BBH - DH (p=0.000278); BH - DH (p=0.017) |
| ASV0779 | d__Bacteria;p__Firmicutes;c__Clostridia;o__Oscillospirales;f__Oscillospiraceae;g__Colidextribacter | 5.5538 | 0.0044074 | 0.035083 | BA - BBH (p=0.00415); BA - BH (p=0.00538); DH - BBH (p=0.0116); DH - BH (p=0.0154) |
| ASV0548 | d__Bacteria;p__Firmicutes;c__Clostridia;o__Oscillospirales;f__Oscillospiraceae;g__Colidextribacter | 5.0742 | 0.0067252 | 0.048666 | BA - BBH (p=0.00333); BA - BH (p=0.00502); BA - DH (p=0.00256) |
| ASV0544 | d__Bacteria;p__Firmicutes;c__Clostridia;o__Oscillospirales;f__Oscillospiraceae;g__NK4A214_group | 14.955 | 7.42E-06 | 0.00028482 | BBH - BA (p=3e-06); BH - BA (p=9e-06); DH - BA (p=0.00694); BBH - DH (p=0.00365); BH - DH (p=0.0135) |
| ASV0986 | d__Bacteria;p__Firmicutes;c__Clostridia;o__Oscillospirales;f__Oscillospiraceae;g__NK4A214_group | 5.4251 | 0.0049306 | 0.038478 | BBH - BA (p=0.0149); BH - BA (p=0.00049); DH - BA (p=0.0334) |
| ASV1058 | d__Bacteria;p__Firmicutes;c__Clostridia;o__Oscillospirales;f__Oscillospiraceae;g__NK4A214_group | 5.1189 | 0.006462 | 0.047046 | BBH - BA (p=0.00251); BBH - BH (p=0.0455); BBH - DH (p=0.0019) |
| ASV0589 | d__Bacteria;p__Firmicutes;c__Clostridia;o__Oscillospirales;f__Oscillospiraceae;g__Papillibacter | 11.008 | 7.56E-05 | 0.0018493 | BBH - BA (p=3.3e-05); BH - BA (p=0.000435); BBH - DH (p=0.000661); BH - DH (p=0.00907) |
| ASV1066 | d__Bacteria;p__Firmicutes;c__Clostridia;o__Oscillospirales;f__Oscillospiraceae;g__UCG-002 | 7.1642 | 0.0011634 | 0.01355 | BBH - BA (p=0.000932); DH - BA (p=0.000328); DH - BH (p=0.0339) |
| ASV1078 | d__Bacteria;p__Firmicutes;c__Clostridia;o__Oscillospirales;f__Oscillospiraceae;g__UCG-005 | 6.338 | 0.0022673 | 0.022373 | BA - BBH (p=0.000948); BA - BH (p=0.000691); BA - DH (p=0.023) |
| ASV1154 | d__Bacteria;p__Firmicutes;c__Clostridia;o__Oscillospirales;f__Oscillospiraceae;g__UCG-005 | 5.6889 | 0.0039217 | 0.032242 | BA - BBH (p=0.0226); BA - BH (p=0.000419); DH - BH (p=0.0207) |
| ASV1059 | d__Bacteria;p__Firmicutes;c__Clostridia;o__Oscillospirales;f__Oscillospiraceae;g__UCG-007 | 11.018 | 7.51E-05 | 0.0018493 | BBH - BA (p=1.7e-05); BBH - BH (p=7.4e-05); BBH - DH (p=0.00107) |
| ASV0606 | d__Bacteria;p__Firmicutes;c__Clostridia;o__Oscillospirales;f__Ruminococcaceae | 80.244 | 2.86E-13 | 3.42E-10 | BA - BBH (p<1e-06); BA - BH (p<1e-06); BA - DH (p<1e-06) |
| ASV0137 | d__Bacteria;p__Firmicutes;c__Clostridia;o__Oscillospirales;f__Ruminococcaceae | 10.945 | 7.88E-05 | 0.0018493 | BH - BA (p=0.000141); DH - BA (p=2.7e-05); BH - BBH (p=0.0186); DH - BBH (p=0.00418) |
| ASV1138 | d__Bacteria;p__Firmicutes;c__Clostridia;o__Oscillospirales;f__Ruminococcaceae;g__[Eubacterium]_siraeum_group | 5.6308 | 0.004123 | 0.033263 | BA - BBH (p=0.00227); BA - BH (p=0.00273); BA - DH (p=0.00182) |
| ASV0575 | d__Bacteria;p__Firmicutes;c__Clostridia;o__Oscillospirales;f__Ruminococcaceae;g__CAG-352 | 7.8231 | 0.00069871 | 0.0091677 | DH - BA (p=0.00176); DH - BBH (p=0.000467); DH - BH (p=0.000323) |
| ASV0187 | d__Bacteria;p__Firmicutes;c__Clostridia;o__Oscillospirales;f__Ruminococcaceae;g__Incertae_Sedis | 10.859 | 8.33E-05 | 0.0018759 | BA - BBH (p=0.00307); BA - BH (p=0.00139); BA - DH (p=6e-06); BBH - DH (p=0.03); BH - DH (p=0.0399) |
| ASV0576 | d__Bacteria;p__Firmicutes;c__Clostridia;o__Oscillospirales;f__Ruminococcaceae;g__Incertae_Sedis | 5.6937 | 0.0039055 | 0.032242 | BBH - BA (p=0.00127); BBH - BH (p=0.00177) |
| ASV1014 | d__Bacteria;p__Firmicutes;c__Clostridia;o__Oscillospirales;f__Ruminococcaceae;g__Pygmaiobacter | 6.4984 | 0.0019867 | 0.020992 | BBH - DH (p=0.00188); BH - DH (p=0.00046) |
| ASV1017 | d__Bacteria;p__Firmicutes;c__Clostridia;o__Oscillospirales;f__Ruminococcaceae;g__Ruminococcaceae | 5.7866 | 0.0036062 | 0.030323 | BA - BBH (p=0.00523); BA - BH (p=0.00531); BA - DH (p=0.000584) |
| ASV0607 | d__Bacteria;p__Firmicutes;c__Clostridia;o__Oscillospirales;f__Ruminococcaceae;g__Ruminococcus | 6.4419 | 0.002081 | 0.021606 | BA - BBH (p=0.00271); BA - BH (p=0.000818); BA - DH (p=0.000963) |
| ASV0553 | d__Bacteria;p__Firmicutes;c__Clostridia;o__Oscillospirales;f__Ruminococcaceae;g__Ruminococcus | 6.03 | 0.0029326 | 0.026327 | DH - BA (p=0.000505); DH - BBH (p=0.00362); DH - BH (p=0.0322) |
| ASV0535 | d__Bacteria;p__Firmicutes;c__Clostridia;o__Oscillospirales;f__Ruminococcaceae;g__Ruminococcus | 5.227 | 0.0058698 | 0.04464 | BH - BA (p=0.00227); DH - BA (p=0.00188) |
| ASV0031 | d__Bacteria;p__Firmicutes;c__Clostridia;o__Oscillospirales;f__Ruminococcaceae;g__uncultured;s__uncultured_Clostridiales | 5.8796 | 0.003331 | 0.028613 | BBH - BA (p=0.00216); BH - BA (p=0.00545); BBH - DH (p=0.0083); BH - DH (p=0.0211) |
| ASV1005 | d__Bacteria;p__Firmicutes;c__Clostridia;o__Oscillospirales;f__UCG-010 | 10.925 | 7.98E-05 | 0.0018493 | BBH - BA (p=0.0123); BH - BA (p=3.4e-05); BH - BBH (p=0.0352); BBH - DH (p=0.0299); BH - DH (p=7.9e-05) |
| ASV0619 | d__Bacteria;p__Firmicutes;c__Clostridia;o__Peptococcales;f__Peptococcaceae | 8.1164 | 0.00056022 | 0.0083613 | BA - BH (p=0.035); BBH - BH (p=0.000207); BBH - DH (p=0.000327) |
| ASV0603 | d__Bacteria;p__Firmicutes;c__Clostridia;o__Peptococcales;f__Peptococcaceae | 6.0061 | 0.0029925 | 0.026454 | BBH - BA (p=0.0277); DH - BA (p=0.000347); DH - BH (p=0.0106) |
| ASV0565 | d__Bacteria;p__Firmicutes;c__Clostridia;o__Peptococcales;f__Peptococcaceae;g__Dehalobacterium | 10.448 | 0.00010894 | 0.0023227 | DH - BA (p=0.000147); DH - BBH (p=0.000693); DH - BH (p=3.3e-05) |
| ASV0407 | d__Bacteria;p__Firmicutes;c__Clostridia;o__Peptostreptococcales-Tissierellales;f__Anaerovoracaceae;g__[Eubacterium]_brachy_group | 7.3391 | 0.0010142 | 0.012232 | BH - BA (p=0.000246); BH - BBH (p=0.00657); BH - DH (p=0.000899) |
| ASV0673 | d__Bacteria;p__Firmicutes;c__Clostridia;o__Peptostreptococcales-Tissierellales;f__Anaerovoracaceae;g__[Eubacterium]_nodatum_group | 9.184 | 0.00025816 | 0.0046704 | BH - BA (p=0.00165); BH - BBH (p=0.000323); BH - DH (p=6.7e-05) |
| ASV0676 | d__Bacteria;p__Firmicutes;c__Clostridia;o__Peptostreptococcales-Tissierellales;f__Anaerovoracaceae;g__[Eubacterium]_nodatum_group | 7.5934 | 0.00083283 | 0.010358 | BBH - BA (p=0.0109); BH - BA (p=0.0165); DH - BA (p=6.3e-05); DH - BH (p=0.0315) |
| ASV0674 | d__Bacteria;p__Firmicutes;c__Clostridia;o__Peptostreptococcales-Tissierellales;f__Anaerovoracaceae;g__[Eubacterium]_nodatum_group | 6.2752 | 0.0023886 | 0.023377 | BH - BA (p=0.000804); BH - BBH (p=0.00126); BH - DH (p=0.0123) |
| ASV0675 | d__Bacteria;p__Firmicutes;c__Clostridia;o__Peptostreptococcales-Tissierellales;f__Anaerovoracaceae;g__Anaerovorax | 8.9671 | 0.00030108 | 0.00521 | BBH - BA (p=0.000336); BH - BA (p=0.018); BBH - DH (p=0.000137); BH - DH (p=0.00893) |
| ASV0689 | d__Bacteria;p__Firmicutes;c__Clostridia;o__Peptostreptococcales-Tissierellales;f__Anaerovoracaceae;g__Family_XIII_AD3011_group | 7.708 | 0.00076279 | 0.009689 | BBH - BA (p=0.0193); BH - BA (p=0.000141); BH - DH (p=0.00143) |
| ASV0688 | d__Bacteria;p__Firmicutes;c__Clostridia;o__Peptostreptococcales-Tissierellales;f__Anaerovoracaceae;g__Family_XIII_AD3011_group | 7.3653 | 0.00099369 | 0.012107 | BBH - BA (p=0.0211); BH - BA (p=0.045); DH - BA (p=8e-05); DH - BBH (p=0.0421); DH - BH (p=0.0133) |
| ASV0805 | d__Bacteria;p__Firmicutes;c__Clostridia;o__Peptostreptococcales-Tissierellales;f__Anaerovoracaceae;g__Family_XIII_AD3011_group;s__bacterium_YSD2010 | 14.238 | 1.10E-05 | 0.00038555 | BA - BBH (p=1.6e-05); BA - BH (p=1e-05); BA - DH (p=1e-05) |
| ASV0692 | d__Bacteria;p__Firmicutes;c__Clostridia;o__Peptostreptococcales-Tissierellales;f__Anaerovoracaceae;g__Mogibacterium | 8.0559 | 0.0005862 | 0.0083915 | BBH - BA (p=0.0019); BH - BA (p=0.000328); DH - BA (p=0.000179) |
| ASV0846 | d__Bacteria;p__Firmicutes;c__Clostridia;o__Peptostreptococcales-Tissierellales;f__Peptostreptococcaceae;g__Intestinibacter | 7.066 | 0.0012572 | 0.014296 | BA - BBH (p=0.000971); BA - BH (p=0.000708); BA - DH (p=0.000708) |
| ASV0843 | d__Bacteria;p__Firmicutes;c__Clostridia;o__Peptostreptococcales-Tissierellales;f__Peptostreptococcaceae;g__Paeniclostridium | 5.8905 | 0.0033003 | 0.028554 | DH - BA (p=0.0263); BH - BBH (p=0.00912); DH - BBH (p=0.000416) |
| ASV0848 | d__Bacteria;p__Firmicutes;c__Clostridia;o__Peptostreptococcales-Tissierellales;f__Peptostreptococcaceae;g__Terrisporobacter | 5.3189 | 0.0054124 | 0.041693 | BH - BA (p=0.0184); DH - BA (p=0.000563); DH - BBH (p=0.043) |
| ASV1141 | d__Bacteria;p__Firmicutes;c__Negativicutes;o__Acidaminococcales;f__Acidaminococcaceae;g__Phascolarctobacterium | 14.901 | 7.63E-06 | 0.00028482 | BBH - BA (p=1.2e-05); BBH - BH (p=7e-06); BBH - DH (p=7e-06) |
| ASV1143 | d__Bacteria;p__Firmicutes;c__Negativicutes;o__Veillonellales-Selenomonadales;f__Selenomonadaceae;g__Anaerovibrio | 6.1412 | 0.0026709 | 0.024928 | BA - BBH (p=0.00125); BA - BH (p=0.000702); BA - DH (p=0.014) |

| | | | | | |
|---|---|---|---|---|---|
| ASV1046 | d__Bacteria;p__Firmicutes;c__Negativicutes;o__Veillonellales-Selenomonadales;f__Veillonellaceae;g__Megasphaera;s__Megasphaera_elsdenii | 7.9034 | 0.0006575 | 0.0087615 | BA - BBH (p=0.000558); BA - BH (p=0.000398); BA - DH (p=0.000398) |
| ASV0827 | d__Bacteria;p__Firmicutes;c__Symbiobacteriia;o__Symbiobacteriales;f__Symbiobacteraceae;g__Symbiobacterium;s__Symbiobacterium_thermophilum | 21.303 | 3.54E-07 | 2.49E-05 | BBH - BA (p=3e-06); DH - BA (p=2.9e-05); BBH - BH (p=2e-06); DH - BH (p=1.8e-05) |
| ASV0523 | d__Bacteria;p__Patescibacteria;c__Saccharimonadia;o__Saccharimonadales;f__Saccharimonadaceae;g__Candidatus_Saccharimonas | 23.114 | 1.67E-07 | 1.42E-05 | BBH - BA (p=0.0108); BH - BA (p<1e-06); DH - BA (p=0.000157); BH - BBH (p=1.6e-05); BH - DH (p=0.000714) |
| ASV0227 | d__Bacteria;p__Proteobacteria;c__Alphaproteobacteria;o__Rhizobiales;f__Beijerinckiaceae;g__Methylobacterium-Methylorubrum | 5.9979 | 0.0030131 | 0.026454 | BBH - BA (p=0.00112); DH - BA (p=0.00835); BBH - BH (p=0.00598); DH - BH (p=0.0421) |
| ASV0269 | d__Bacteria;p__Proteobacteria;c__Alphaproteobacteria;o__Rhodospirillales | 6.0914 | 0.0027849 | 0.025578 | BA - BBH (p=0.00256); BA - BH (p=0.000695); BA - DH (p=0.0029) |
| ASV0161 | d__Bacteria;p__Proteobacteria;c__Alphaproteobacteria;o__Rickettsiales | 7.2719 | 0.0010689 | 0.012763 | BBH - BA (p=0.000861); BBH - BH (p=0.00234); BBH - DH (p=0.000261) |
| ASV0160 | d__Bacteria;p__Proteobacteria;c__Alphaproteobacteria;o__Rickettsiales | 5.2041 | 0.0059902 | 0.045268 | BBH - BA (p=0.00575); BBH - BH (p=0.0253); BBH - DH (p=0.000903) |
| ASV0872 | d__Bacteria;p__Proteobacteria;c__Gammaproteobacteria;o__Burkholderiales;f__Sutterellaceae;g__Sutterella | 8.0828 | 0.00057448 | 0.0083915 | BA - BBH (p=0.000337); BA - BH (p=0.000236); BA - DH (p=0.000965) |
| ASV0842 | d__Bacteria;p__Spirochaetota;c__Spirochaetia;o__Spirochaetales;f__Spirochaetaceae;g__Sphaerochaeta | 8.7619 | 0.00034884 | 0.0059501 | BBH - BA (p=0.000132); BH - BA (p=0.00153); BBH - DH (p=0.00173); BH - DH (p=0.0196) |
| ASV0049 | d__Bacteria;p__Spirochaetota;c__Spirochaetia;o__Spirochaetales;f__Spirochaetaceae;g__Treponema | 9.9941 | 0.00014755 | 0.002986 | BA - BBH (p=0.000254); BA - BH (p=9.5e-05); BA - DH (p=7.9e-05) |
| ASV0050 | d__Bacteria;p__Spirochaetota;c__Spirochaetia;o__Spirochaetales;f__Spirochaetaceae;g__Treponema | 6.3719 | 0.0022047 | 0.022208 | BA - BBH (p=0.0104); BA - BH (p=0.000758); BA - DH (p=0.000724) |
| ASV0048 | d__Bacteria;p__Spirochaetota;c__Spirochaetia;o__Spirochaetales;f__Spirochaetaceae;g__Treponema;s__Treponema_berlinense | 8.6222 | 0.00038595 | 0.0063496 | BA - BBH (p=0.000353); BA - BH (p=0.000248); BA - DH (p=0.000248) |
