## Supplementary material for "Multi-domain analysis of the fecal microbiome of dairy heifers, beef heifers, bred beef heifers, and adult beef cows in the central United States using paired 16S rRNA and 18S rRNA metabarcoding": Table S5

**Table S5. Results of correlation analyses of 16S rRNA and 18S rRNA metabarcoding data among all samples using Spearman's rank correlation. Only comparisons with raw p value < 0.05 and rho > 0.5 are listed.**

| Taxon1 | Taxon2 | Correlation | p value | Statistic |
|---|---|---|---|---|
| g__Acremonium | g__Aspergillus | 0.5838 | 9.00E-04 | 1689.76 |
| g__Acremonium | g__Chaetomium | 0.8961 | 0 | 421.635 |
| g__Acremonium | g__Lichtheimia | 0.6314 | 2.00E-04 | 1496.63 |
| g__Acremonium | g__Microascus | 0.8793 | 0 | 490.226 |
| g__Acremonium | g__Neocallimastigaceae | -0.6249 | 3.00E-04 | 6597.2 |
| g__Acremonium | g__Rhizomucor | 0.8975 | 0 | 416.314 |
| g__Acremonium | g__Schizoplasmodiida | 0.7599 | 0 | 974.938 |
| g__Acremonium | g__Simplicimonas | -0.5045 | 0.0053 | 6108.28 |
| g__Acremonium | g__Spiromyces | 0.5686 | 0.0013 | 1751.33 |
| g__Acremonium | g__Stemonitis | 0.6248 | 3.00E-04 | 1523.14 |
| g__Acremonium | g__Talaromyces | 0.9448 | 0 | 223.93 |
| g__Acremonium | g__Trichocomaceae | 0.833 | 0 | 677.843 |
| g__Acremonium | g__Trichomonascus | 0.7085 | 0 | 1183.64 |
| g__Acremonium | g__Tritirachium | 0.9682 | 0 | 129.113 |
| g__Acremonium | s__Brevibacillus_thermoruber | 0.5509 | 0.002 | 1823.18 |
| g__Acremonium | s__Symbiobacterium_thermophilum | 0.6453 | 2.00E-04 | 1440.11 |
| g__Acremonium | s__Thermobifida_fusca | 0.5229 | 0.0036 | 1937.2 |
| g__Acremonium | s__Thermobispora_bispora | 0.5268 | 0.0033 | 1921.05 |
| g__Acremonium | s__uncultured_Bacteroides | -0.5187 | 0.0039 | 6165.85 |
| g__Acremonium | s__uncultured_compost | 0.5128 | 0.0045 | 1978.14 |
| g__Arachnomyces | s__Bacillus_smithii | 0.5549 | 0.0018 | 1807.03 |
| g__Arachnomyces | s__Brevibacillus_thermoruber | 0.6743 | 1.00E-04 | 1322.34 |
| g__Arachnomyces | s__Caenibacillus_caldisaponilyticus | 0.5774 | 0.001 | 1715.62 |
| g__Arachnomyces | s__Paenibacillus_barengoltzii | 0.5388 | 0.0026 | 1872.55 |
| g__Arachnomyces | s__Symbiobacterium_thermophilum | 0.6744 | 1.00E-04 | 1321.74 |
| g__Arachnomyces | s__Thermobacillus_composti | 0.506 | 0.0051 | 2005.46 |
| g__Arachnomyces | s__Thermobifida_fusca | 0.5433 | 0.0023 | 1854.33 |
| g__Arachnomyces | s__Thermobispora_bispora | 0.5871 | 8.00E-04 | 1676.54 |
| g__Arachnomyces | s__uncultured_compost | 0.5178 | 0.004 | 1957.74 |
| g__Arachnomyces | s__uncultured_Thermobacillus | 0.6424 | 2.00E-04 | 1452.02 |
| g__Arthrinium | g__Thelebolus | 0.5485 | 0.0021 | 1832.91 |
| g__Arthrinium | s__uncultured_Acetanaerobacterium | 0.505 | 0.0052 | 2009.53 |
| g__Arthrinium | s__uncultured_Paludibacter | 0.5242 | 0.0035 | 1931.61 |
| g__Aspergillus | g__Chaetomium | 0.6145 | 4.00E-04 | 1565.22 |
| g__Aspergillus | g__Microascus | 0.5541 | 0.0018 | 1810.28 |
| g__Aspergillus | g__Rhizomucor | 0.5546 | 0.0018 | 1808.45 |
| g__Aspergillus | g__Talaromyces | 0.6387 | 2.00E-04 | 1466.8 |
| g__Aspergillus | g__Trichocomaceae | 0.6011 | 6.00E-04 | 1619.46 |
| g__Aspergillus | g__Tritirachium | 0.6174 | 4.00E-04 | 1553.17 |
| g__Aspergillus | g__Wallemia | 0.5309 | 0.003 | 1904.69 |
| g__Aspergillus | s__Bacillus_vietnamensis | -0.5101 | 0.0047 | 6131.02 |
| g__Aspergillus | s__metagenome | -0.6195 | 3.00E-04 | 6575.31 |
| g__Chaetomium | g__Lichtheimia | 0.6666 | 1.00E-04 | 1353.66 |
| g__Chaetomium | g__Microascus | 0.9041 | 0 | 389.357 |
| g__Chaetomium | g__Neocallimastigaceae | -0.5368 | 0.0027 | 6239.48 |
| g__Chaetomium | g__Rhizomucor | 0.8439 | 0 | 633.787 |
| g__Chaetomium | g__Schizoplasmodiida | 0.8136 | 0 | 756.761 |
| g__Chaetomium | g__Spiromyces | 0.5426 | 0.0024 | 1857 |
| g__Chaetomium | g__Stemonitis | 0.5225 | 0.0036 | 1938.73 |
| g__Chaetomium | g__Talaromyces | 0.8862 | 0 | 461.983 |
| g__Chaetomium | g__Trichocomaceae | 0.8842 | 0 | 470.052 |
| g__Chaetomium | g__Trichomonascus | 0.6791 | 1.00E-04 | 1302.74 |
| g__Chaetomium | g__Tritirachium | 0.9125 | 0 | 355.061 |
| g__Chaetomium | s__metagenome | -0.5246 | 0.0035 | 6189.96 |
| g__Chaetomium | s__Symbiobacterium_thermophilum | 0.5409 | 0.0025 | 1864.14 |
| g__Chaetomium | s__uncultured_compost | 0.56 | 0.0016 | 1786.47 |
| g__Cryptosporidium | g__Entamoeba | -0.5541 | 0.0018 | 6309.69 |
| g__Cryptosporidium | s__Bacillus_smithii | 0.5033 | 0.0054 | 2016.77 |
| g__Cryptosporidium | s__bacterium | 0.5033 | 0.0054 | 2016.47 |
| g__Cryptosporidium | s__uncultured_Bacteroidales | -0.5309 | 0.003 | 6215.53 |
| g__Cyllamyces | g__Microascus | 0.5168 | 0.0041 | 1961.91 |
| g__Cyllamyces | g__Schizoplasmodiida | 0.5725 | 0.0012 | 1735.5 |
| g__Echinamoeba | g__Pilobolus | 0.5462 | 0.0022 | 1842.55 |
| g__Eimeria | s__Bacillus_vietnamensis | 0.5787 | 0.001 | 1710.64 |
| g__Eimeria | s__bacterium_PS7 | -0.5167 | 0.0041 | 6157.82 |
| g__Entamoeba | g__Simplicimonas | 0.6433 | 2.00E-04 | 1448.05 |
| g__Entamoeba | g__Trebouxiophyceae | -0.5132 | 0.0044 | 6143.4 |
| g__Entamoeba | Not_Assigned | -0.6431 | 2.00E-04 | 6670.87 |
| g__Entamoeba | s__Bacillus_coagulans | -0.6655 | 1.00E-04 | 6761.86 |
| g__Entamoeba | s__Bacillus_smithii | -0.6571 | 1.00E-04 | 6727.99 |
| g__Entamoeba | s__bacterium | -0.7068 | 0 | 6929.7 |
| g__Entamoeba | s__Brevibacillus_thermoruber | -0.565 | 0.0014 | 6353.8 |
| g__Entamoeba | s__Caenibacillus_caldisaponilyticus | -0.6033 | 5.00E-04 | 6509.57 |
| g__Entamoeba | s__rumen_bacterium | -0.5625 | 0.0015 | 6343.81 |
| g__Entamoeba | s__Thermobacillus_composti | -0.5846 | 9.00E-04 | 6433.52 |
| g__Entamoeba | s__Thermobifida_fusca | -0.5713 | 0.0012 | 6379.4 |
| g__Entamoeba | s__Thermobispora_bispora | -0.5548 | 0.0018 | 6312.58 |
| g__Entamoeba | s__uncultured_compost | -0.5416 | 0.0024 | 6258.99 |
| g__Entamoeba | s__uncultured_rumen | -0.7284 | 0 | 7017.39 |
| g__Fusarium | g__Naganishia | 0.5547 | 0.0018 | 1807.9 |
| g__Fusarium | g__Rhodotorula | 0.6699 | 1.00E-04 | 1340.33 |
| g__Fusarium | s__Brevibacillus_thermoruber | -0.5021 | 0.0055 | 6098.38 |
| g__Fusarium | s__gut_metagenome | 0.5859 | 8.00E-04 | 1681.05 |
| g__Fusarium | s__uncultured_Burkholderiales | 0.6243 | 3.00E-04 | 1525.24 |
| g__Fusarium | s__uncultured_Erysipelotrichaceae | 0.6916 | 0 | 1251.97 |
| g__Gregarina | Not_Assigned | 0.5183 | 0.004 | 1955.51 |
| g__Gregarina | s__uncultured_Dehalobacterium | 0.5659 | 0.0014 | 1762.56 |
| g__Hannaella | g__Trebouxiophyceae | 0.7566 | 0 | 988.385 |
| g__Hannaella | s__Saccharopolyspora_rectivirgula | 0.5087 | 0.0048 | 1994.49 |
| g__Incertae_Sedis | s__Bacillus_coagulans | 0.5628 | 0.0015 | 1775.05 |
| g__Incertae_Sedis | s__Bacillus_thermoamylovorans | 0.7067 | 0 | 1190.62 |
| g__Lichtheimia | g__Microascus | 0.6215 | 3.00E-04 | 1536.91 |
| g__Lichtheimia | g__Rhizomucor | 0.6995 | 0 | 1220.01 |
| g__Lichtheimia | g__Schizoplasmodiida | 0.6041 | 5.00E-04 | 1607.53 |
| g__Lichtheimia | g__Talaromyces | 0.6095 | 4.00E-04 | 1585.23 |
| g__Lichtheimia | g__Trichocomaceae | 0.7087 | 0 | 1182.5 |
| g__Lichtheimia | g__Tritirachium | 0.6691 | 1.00E-04 | 1343.6 |
| g__Microascus | g__Rhizomucor | 0.9039 | 0 | 390.136 |

| | | | | |
|---|---|---|---|---|
| g__Microascus | g__Schizoplasmodiida | 0.8468 | 0 | 622.185 |
| g__Microascus | g__Talaromyces | 0.8589 | 0 | 572.939 |
| g__Microascus | g__Trichocomaceae | 0.8812 | 0 | 482.157 |
| g__Microascus | g__Trichomonascus | 0.5701 | 0.0012 | 1745.43 |
| g__Microascus | g__Tritirachium | 0.9011 | 0 | 401.461 |
| g__Neocallimastigaceae | g__Talaromyces | -0.6343 | 2.00E-04 | 6635.39 |
| g__Neocallimastigaceae | g__Trebouxiophyceae | -0.5286 | 0.0032 | 6205.98 |
| g__Neocallimastigaceae | g__Tritirachium | -0.5487 | 0.0021 | 6287.72 |
| g__Neocallimastigaceae | s__Bacillus_smithii | -0.5544 | 0.0018 | 6310.79 |
| g__Neocallimastigaceae | s__Brevibacillus_thermoruber | -0.5884 | 8.00E-04 | 6448.73 |
| g__Neocallimastigaceae | s__Caenibacillus_caldisaponilyticus | -0.6455 | 2.00E-04 | 6680.53 |
| g__Neocallimastigaceae | s__Symbiobacterium_thermophilum | -0.5988 | 6.00E-04 | 6490.98 |
| g__Neocallimastigaceae | s__Thermobacillus_composti | -0.6268 | 3.00E-04 | 6604.86 |
| g__Neocallimastigaceae | s__Thermobifida_fusca | -0.6705 | 1.00E-04 | 6782.09 |
| g__Neocallimastigaceae | s__Thermobispora_bispora | -0.6364 | 2.00E-04 | 6643.72 |
| g__Neocallimastigaceae | s__uncultured_Acetanaerobacterium | -0.6859 | 0 | 6844.76 |
| g__Neocallimastigaceae | s__uncultured_compost | -0.5128 | 0.0044 | 6142.01 |
| g__Neocallimastigaceae | s__uncultured_Paludibacter | -0.6469 | 1.00E-04 | 6686.59 |
| g__Neocallimastigaceae | s__uncultured_Parabacteroides | -0.6173 | 4.00E-04 | 6566.16 |
| g__Neocallimastigaceae | s__uncultured_Thermobacillus | -0.615 | 4.00E-04 | 6556.73 |
| g__Neocallimastigaceae | s__uncultured_Veillonellaceae | -0.5143 | 0.0043 | 6148.06 |
| g__Pilobolus | g__Trichocomaceae | 0.5079 | 0.0049 | 1997.86 |
| g__Pilobolus | s__Olsenella_umbonata | 0.6008 | 6.00E-04 | 1620.82 |
| g__Rhabditida | s__bacterium_YE57 | 0.5229 | 0.0036 | 1937.1 |
| g__Rhabditida | s__gut_metagenome | -0.586 | 8.00E-04 | 6439.04 |
| g__Rhabditida | s__Lactobacillus_mucosae | 0.5442 | 0.0023 | 1850.57 |
| g__Rhabditida | s__Symbiobacterium_thermophilum | 0.5301 | 0.0031 | 1907.76 |
| g__Rhabditida | s__uncultured_bacterium | -0.5308 | 0.0031 | 6215.1 |
| g__Rhabditida | s__uncultured_compost | 0.5437 | 0.0023 | 1852.67 |
| g__Rhabditida | s__uncultured_Dehalobacterium | 0.5833 | 9.00E-04 | 1691.94 |
| g__Rhizomucor | g__Schizoplasmodiida | 0.8001 | 0 | 811.745 |
| g__Rhizomucor | g__Stemonitis | 0.6011 | 6.00E-04 | 1619.47 |
| g__Rhizomucor | g__Talaromyces | 0.8885 | 0 | 452.559 |
| g__Rhizomucor | g__Trichocomaceae | 0.8846 | 0 | 468.668 |
| g__Rhizomucor | g__Trichomonascus | 0.5304 | 0.0031 | 1906.74 |
| g__Rhizomucor | g__Tritirachium | 0.9277 | 0 | 293.481 |
| g__Rhodotorula | g__Wallemia | -0.5359 | 0.0027 | 6235.84 |
| g__Rhodotorula | s__uncultured_Burkholderiales | 0.5863 | 8.00E-04 | 1679.58 |
| g__Schizoplasmodiida | g__Stemonitis | 0.5554 | 0.0018 | 1805.16 |
| g__Schizoplasmodiida | g__Talaromyces | 0.6755 | 1.00E-04 | 1317.5 |
| g__Schizoplasmodiida | g__Trichocomaceae | 0.7307 | 0 | 1093.2 |
| g__Schizoplasmodiida | g__Trichomonascus | 0.5287 | 0.0032 | 1913.66 |
| g__Schizoplasmodiida | g__Tritirachium | 0.7739 | 0 | 917.845 |
| g__Schizoplasmodiida | s__bacterium | 0.5299 | 0.0031 | 1908.74 |
| g__Schizoplasmodiida | s__uncultured_prokaryote | -0.5101 | 0.0047 | 6131.09 |
| g__Simplicimonas | Not_Assigned | -0.5004 | 0.0057 | 6091.6 |
| g__Simplicimonas | s__Bacillus_coagulans | -0.6752 | 1.00E-04 | 6801.41 |
| g__Simplicimonas | s__Bacillus_smithii | -0.5896 | 8.00E-04 | 6453.78 |
| g__Simplicimonas | s__Brevibacillus_thermoruber | -0.5857 | 8.00E-04 | 6438.01 |
| g__Simplicimonas | s__Caenibacillus_caldisaponilyticus | -0.5728 | 0.0012 | 6385.39 |
| g__Simplicimonas | s__rumen_bacterium | -0.5876 | 8.00E-04 | 6445.77 |
| g__Simplicimonas | s__Thermobacillus_composti | -0.606 | 5.00E-04 | 6520.32 |
| g__Simplicimonas | s__Thermobifida_fusca | -0.5861 | 8.00E-04 | 6439.74 |
| g__Simplicimonas | s__Thermobispora_bispora | -0.5533 | 0.0019 | 6306.28 |
| g__Spiromyces | g__Talaromyces | 0.5314 | 0.003 | 1902.48 |
| g__Spiromyces | g__Tritirachium | 0.5321 | 0.003 | 1899.81 |
| g__Spiromyces | Not_Assigned | 0.5796 | 0.001 | 1706.7 |
| g__Spiromyces | s__bacterium_PS7 | 0.5276 | 0.0033 | 1917.98 |
| g__Spiromyces | s__Symbiobacterium_thermophilum | 0.525 | 0.0035 | 1928.49 |
| g__Stemonitis | g__Talaromyces | 0.5991 | 6.00E-04 | 1627.68 |
| g__Stemonitis | g__Trichomonascus | 0.7149 | 0 | 1157.54 |
| g__Stemonitis | g__Tritirachium | 0.6481 | 1.00E-04 | 1428.81 |
| g__Stemonitis | s__uncultured_bacterium | -0.509 | 0.0048 | 6126.67 |
| g__Talaromyces | g__Trichocomaceae | 0.9205 | 0 | 322.783 |
| g__Talaromyces | g__Trichomonascus | 0.6769 | 1.00E-04 | 1311.73 |
| g__Talaromyces | g__Tritirachium | 0.9538 | 0 | 187.617 |
| g__Talaromyces | s__Brevibacillus_thermoruber | 0.5811 | 9.00E-04 | 1700.83 |
| g__Talaromyces | s__Symbiobacterium_thermophilum | 0.6917 | 0 | 1251.89 |
| g__Talaromyces | s__Thermobacillus_composti | 0.5485 | 0.0021 | 1833.04 |
| g__Talaromyces | s__Thermobispora_bispora | 0.5494 | 0.002 | 1829.29 |
| g__Talaromyces | s__uncultured_compost | 0.558 | 0.0017 | 1794.63 |
| g__Trebouxiophyceae | Not_Assigned | 0.5445 | 0.0023 | 1849.27 |
| g__Trebouxiophyceae | s__Caenibacillus_caldisaponilyticus | 0.5083 | 0.0049 | 1996.44 |
| g__Trebouxiophyceae | s__Saccharopolyspora_rectivirgula | 0.5229 | 0.0036 | 1937.14 |
| g__Trebouxiophyceae | s__Thermobifida_fusca | 0.5957 | 7.00E-04 | 1641.4 |
| g__Trebouxiophyceae | s__uncultured_Paludibacter | 0.6375 | 2.00E-04 | 1471.66 |
| g__Trichocomaceae | g__Trichomonascus | 0.5225 | 0.0036 | 1938.69 |
| g__Trichocomaceae | g__Tritirachium | 0.8957 | 0 | 423.652 |
| g__Trichocomaceae | s__Symbiobacterium_thermophilum | 0.5515 | 0.0019 | 1820.87 |
| g__Trichomonascus | g__Tritirachium | 0.6813 | 0 | 1293.75 |
| g__Trichomonascus | s__Bacillus_vietnamensis | -0.6278 | 3.00E-04 | 6609.05 |
| g__Trichomonascus | s__Thermobispora_bispora | 0.5678 | 0.0013 | 1754.67 |
| g__Trichomonascus | s__uncultured_compost | 0.5533 | 0.0019 | 1813.72 |
| g__Trichostomatia | s__Caenibacillus_caldisaponilyticus | 0.587 | 8.00E-04 | 1676.7 |
| g__Trichostomatia | s__uncultured_Alphaproteobacteria | 0.5064 | 0.0051 | 2004.21 |
| g__Tritirachium | s__Brevibacillus_thermoruber | 0.5309 | 0.003 | 1904.74 |
| g__Tritirachium | s__metagenome | -0.532 | 0.003 | 6220.1 |
| g__Tritirachium | s__Symbiobacterium_thermophilum | 0.6709 | 1.00E-04 | 1336.27 |
| g__Tritirachium | s__Thermobispora_bispora | 0.5183 | 0.004 | 1955.71 |
| g__Tritirachium | s__uncultured_Bacteroides | -0.5127 | 0.0045 | 6141.75 |
| g__Tritirachium | s__uncultured_compost | 0.5228 | 0.0036 | 1937.36 |
| Not_Assigned | s__Anaerofustis_stercorihominis | 0.5066 | 0.005 | 2003.23 |
| Not_Assigned | s__Bacillus_coagulans | 0.5071 | 0.005 | 2001.35 |
| Not_Assigned | s__bacterium | 0.5101 | 0.0047 | 1988.89 |
| Not_Assigned | s__Brevibacillus_thermoruber | 0.6281 | 3.00E-04 | 1510.03 |
| Not_Assigned | s__Caenibacillus_caldisaponilyticus | 0.564 | 0.0014 | 1770.19 |
| Not_Assigned | s__rumen_bacterium | 0.6549 | 1.00E-04 | 1400.95 |
| Not_Assigned | s__Thermobacillus_composti | 0.5719 | 0.0012 | 1737.93 |
| Not_Assigned | s__Thermobifida_fusca | 0.5291 | 0.0032 | 1911.72 |
| Not_Assigned | s__Thermobispora_bispora | 0.5886 | 8.00E-04 | 1670.42 |
| Not_Assigned | s__uncultured_compost | 0.7001 | 0 | 1217.68 |

| | | | | |
|---|---|---|---|---|
| Not_Assigned | s__uncultured_Thermobacillus | 0.6216 | 3.00E-04 | 1536.49 |
| s__Anaerofustis_stercorihominis | s__bacterium_YE57 | 0.5 | 0.0057 | 2029.94 |
| s__Anaerofustis_stercorihominis | s__rumen_bacterium | 0.5805 | 0.001 | 1703.04 |
| s__Anaerofustis_stercorihominis | s__Thermobispora_bispora | 0.5596 | 0.0016 | 1787.89 |
| s__Anaerofustis_stercorihominis | s__uncultured_compost | 0.5643 | 0.0014 | 1768.94 |
| s__Bacillus_coagulans | s__Bacillus_thermoamylovorans | 0.7837 | 0 | 878.107 |
| s__Bacillus_coagulans | s__bacterium | 0.5349 | 0.0028 | 1888.39 |
| s__Bacillus_coagulans | s__rumen_bacterium | 0.5034 | 0.0054 | 2016.26 |
| s__Bacillus_coagulans | s__Thermobifida_fusca | 0.5163 | 0.0041 | 1963.87 |
| s__Bacillus_coagulans | s__uncultured_actinobacterium | 0.5338 | 0.0029 | 1892.75 |
| s__Bacillus_smithii | s__Bacillus_vietnamensis | -0.5344 | 0.0028 | 6229.8 |
| s__Bacillus_smithii | s__Brevibacillus_thermoruber | 0.8229 | 0 | 719.226 |
| s__Bacillus_smithii | s__Caenibacillus_caldisaponilyticus | 0.862 | 0 | 560.401 |
| s__Bacillus_smithii | s__Paenibacillus_barengoltzii | 0.5341 | 0.0028 | 1891.71 |
| s__Bacillus_smithii | s__Saccharopolyspora_rectivirgula | 0.5098 | 0.0047 | 1990.31 |
| s__Bacillus_smithii | s__Symbiobacterium_thermophilum | 0.6466 | 2.00E-04 | 1434.82 |
| s__Bacillus_smithii | s__Thermasporomyces_composti | 0.6299 | 3.00E-04 | 1502.61 |
| s__Bacillus_smithii | s__Thermobacillus_composti | 0.8337 | 0 | 675.012 |
| s__Bacillus_smithii | s__Thermobifida_fusca | 0.752 | 0 | 1006.78 |
| s__Bacillus_smithii | s__Thermobispora_bispora | 0.7782 | 0 | 900.321 |
| s__Bacillus_smithii | s__uncultured_compost | 0.5983 | 6.00E-04 | 1631.02 |
| s__Bacillus_smithii | s__uncultured_Thermobacillus | 0.7088 | 0 | 1182.28 |
| s__Bacillus_thermoamylovorans | s__bacterium | 0.5147 | 0.0043 | 1970.28 |
| s__Bacillus_thermoamylovorans | s__Paenibacillus_barengoltzii | 0.5146 | 0.0043 | 1970.67 |
| s__Bacillus_vietnamensis | s__Brevibacillus_thermoruber | -0.5734 | 0.0011 | 6387.87 |
| s__Bacillus_vietnamensis | s__Caenibacillus_caldisaponilyticus | -0.5549 | 0.0018 | 6312.71 |
| s__Bacillus_vietnamensis | s__metagenome | 0.5102 | 0.0047 | 1988.73 |
| s__Bacillus_vietnamensis | s__Symbiobacterium_thermophilum | -0.5338 | 0.0029 | 6227.34 |
| s__Bacillus_vietnamensis | s__Thermobispora_bispora | -0.6315 | 2.00E-04 | 6624.06 |
| s__Bacillus_vietnamensis | s__uncultured_bacterium | 0.5644 | 0.0014 | 1768.6 |
| s__Bacillus_vietnamensis | s__uncultured_Mollicutes | 0.5511 | 0.0019 | 1822.71 |
| s__Bacillus_vietnamensis | s__uncultured_Thermobacillus | -0.5028 | 0.0054 | 6101.52 |
| s__bacterium | s__bacterium_AC2043 | 0.5349 | 0.0028 | 1888.11 |
| s__bacterium | s__Campylobacter_hyointestinalis | 0.5472 | 0.0021 | 1838.33 |
| s__bacterium | s__rumen_bacterium | 0.5591 | 0.0016 | 1790.2 |
| s__bacterium | s__uncultured_rumen | 0.7235 | 0 | 1122.39 |
| s__bacterium_AC2043 | s__bacterium_FE2018 | 0.5966 | 6.00E-04 | 1637.97 |
| s__bacterium_AC2043 | s__uncultured_actinobacterium | 0.5408 | 0.0025 | 1864.38 |
| s__bacterium_AC2043 | s__uncultured_Coriobacteriaceae | 0.5832 | 9.00E-04 | 1692.26 |
| s__bacterium_AC2043 | s__uncultured_Mollicutes | 0.593 | 7.00E-04 | 1652.52 |
| s__bacterium_AC2043 | s__uncultured_rumen | 0.7104 | 0 | 1175.65 |
| s__bacterium_AC2043 | s__unidentified_rumen | 0.5797 | 0.001 | 1706.24 |
| s__bacterium_enrichment | s__Paenibacillus_barengoltzii | 0.5317 | 0.003 | 1901.12 |
| s__bacterium_enrichment | s__uncultured_archaeon | 0.521 | 0.0038 | 1944.93 |
| s__bacterium_enrichment | s__uncultured_Bacteroidetes | 0.5022 | 0.0055 | 2021.2 |
| s__bacterium_enrichment | s__uncultured_Firmicutes | 0.5541 | 0.0018 | 1810.52 |
| s__bacterium_FE2018 | s__rumen_bacterium | 0.6078 | 5.00E-04 | 1592.28 |
| s__bacterium_FE2018 | s__uncultured_actinobacterium | 0.5318 | 0.003 | 1900.78 |
| s__bacterium_FE2018 | s__uncultured_rumen | 0.6793 | 1.00E-04 | 1301.9 |
| s__bacterium_FE2018 | s__unidentified_rumen | 0.6089 | 5.00E-04 | 1587.79 |
| s__bacterium_PS7 | s__bacterium_YE57 | 0.5284 | 0.0032 | 1914.88 |
| s__bacterium_PS7 | s__Symbiobacterium_thermophilum | 0.6047 | 5.00E-04 | 1604.74 |
| s__bacterium_WCE2003 | s__uncultured_Clostridium | 0.5403 | 0.0025 | 1866.26 |
| s__bacterium_YE57 | s__Thermomonospora_curvata | 0.5158 | 0.0042 | 1965.68 |
| s__bacterium_YE57 | s__uncultured_Dehalobacterium | 0.581 | 0.001 | 1701.33 |
| s__bacterium_YE57 | s__uncultured_Lachnospiraceae | 0.5065 | 0.005 | 2003.49 |
| s__bacterium_YGD2005 | s__unidentified | 0.5514 | 0.0019 | 1821.45 |
| s__Brevibacillus_thermoruber | s__Caenibacillus_caldisaponilyticus | 0.8979 | 0 | 414.374 |
| s__Brevibacillus_thermoruber | s__Paenibacillus_barengoltzii | 0.656 | 1.00E-04 | 1396.69 |
| s__Brevibacillus_thermoruber | s__rumen_bacterium | 0.5224 | 0.0036 | 1939.02 |
| s__Brevibacillus_thermoruber | s__Saccharopolyspora_rectivirgula | 0.5032 | 0.0054 | 2017.1 |
| s__Brevibacillus_thermoruber | s__Symbiobacterium_thermophilum | 0.8122 | 0 | 762.573 |
| s__Brevibacillus_thermoruber | s__Thermasporomyces_composti | 0.7261 | 0 | 1111.9 |
| s__Brevibacillus_thermoruber | s__Thermobacillus_composti | 0.9275 | 0 | 294.258 |
| s__Brevibacillus_thermoruber | s__Thermobifida_fusca | 0.788 | 0 | 860.752 |
| s__Brevibacillus_thermoruber | s__Thermobispora_bispora | 0.8761 | 0 | 502.863 |
| s__Brevibacillus_thermoruber | s__uncultured_Acetanaerobacterium | 0.5035 | 0.0054 | 2015.86 |
| s__Brevibacillus_thermoruber | s__uncultured_Alphaproteobacteria | 0.536 | 0.0027 | 1883.91 |
| s__Brevibacillus_thermoruber | s__uncultured_compost | 0.7396 | 0 | 1057.25 |
| s__Brevibacillus_thermoruber | s__uncultured_Thermobacillus | 0.8799 | 0 | 487.594 |
| s__Caenibacillus_caldisaponilyticus | s__Paenibacillus_barengoltzii | 0.6552 | 1.00E-04 | 1400.07 |
| s__Caenibacillus_caldisaponilyticus | s__Saccharopolyspora_rectivirgula | 0.5066 | 0.005 | 2003.1 |
| s__Caenibacillus_caldisaponilyticus | s__Symbiobacterium_thermophilum | 0.7136 | 0 | 1162.81 |
| s__Caenibacillus_caldisaponilyticus | s__Thermasporomyces_composti | 0.6855 | 0 | 1276.76 |
| s__Caenibacillus_caldisaponilyticus | s__Thermobacillus_composti | 0.8967 | 0 | 419.206 |
| s__Caenibacillus_caldisaponilyticus | s__Thermobifida_fusca | 0.7882 | 0 | 860.045 |
| s__Caenibacillus_caldisaponilyticus | s__Thermobispora_bispora | 0.906 | 0 | 381.549 |
| s__Caenibacillus_caldisaponilyticus | s__uncultured_Alphaproteobacteria | 0.6326 | 2.00E-04 | 1491.64 |
| s__Caenibacillus_caldisaponilyticus | s__uncultured_compost | 0.7168 | 0 | 1149.86 |
| s__Caenibacillus_caldisaponilyticus | s__uncultured_Paludibacter | 0.5234 | 0.0036 | 1934.86 |
| s__Caenibacillus_caldisaponilyticus | s__uncultured_Thermobacillus | 0.8697 | 0 | 528.958 |
| s__Clostridiales_bacterium | s__Lactobacillus_mucosae | -0.615 | 4.00E-04 | 6556.79 |
| s__Clostridiales_bacterium | s__uncultured_actinobacterium | 0.5982 | 6.00E-04 | 1631.3 |
| s__Clostridiales_bacterium | s__uncultured_Mollicutes | 0.6413 | 2.00E-04 | 1456.5 |
| s__Clostridiales_bacterium | s__uncultured_Paludibacter | 0.7319 | 0 | 1088.56 |
| s__Clostridiales_bacterium | s__uncultured_Parabacteroides | 0.6302 | 2.00E-04 | 1501.5 |
| s__Clostridiales_bacterium | s__uncultured_Veillonellaceae | 0.6401 | 2.00E-04 | 1461.39 |
| s__Clostridiales_bacterium | s__unidentified_rumen | 0.5105 | 0.0047 | 1987.19 |
| s__gut_metagenome | s__uncultured_bacterium | 0.6388 | 2.00E-04 | 1466.36 |
| s__gut_metagenome | s__uncultured_Bacteroidales | 0.5304 | 0.0031 | 1906.47 |
| s__gut_metagenome | s__uncultured_Erysipelotrichaceae | 0.5649 | 0.0014 | 1766.44 |
| s__gut_metagenome | s__uncultured_Mollicutes | 0.5699 | 0.0012 | 1746.16 |
| s__Lactobacillus_amylovorus | s__Lactobacillus_mucosae | 0.9073 | 0 | 376.437 |
| s__Lactobacillus_amylovorus | s__uncultured_Dehalobacterium | 0.5917 | 7.00E-04 | 1657.65 |
| s__Lactobacillus_amylovorus | s__uncultured_organism | 0.5555 | 0.0018 | 1804.87 |
| s__Lactobacillus_amylovorus | s__uncultured_Parabacteroides | -0.5251 | 0.0034 | 6191.94 |
| s__Lactobacillus_amylovorus | s__unidentified | 0.5259 | 0.0034 | 1924.82 |
| s__Lactobacillus_mucosae | s__uncultured_Dehalobacterium | 0.5925 | 7.00E-04 | 1654.54 |
| s__Lactobacillus_mucosae | s__uncultured_organism | 0.5804 | 0.001 | 1703.68 |
| s__Lactobacillus_mucosae | s__uncultured_Parabacteroides | -0.6088 | 5.00E-04 | 6531.67 |
| s__Lactobacillus_mucosae | s__uncultured_Ruminococcaceae | 0.5837 | 9.00E-04 | 1690.25 |

| | | | | |
|---|---|---|---|---|
| s__Lactobacillus_mucosae | s__unidentified | 0.6131 | 4.00E-04 | 1570.73 |
| s__metagenome | s__uncultured_Anaeroplasma | 0.5726 | 0.0012 | 1735.13 |
| s__metagenome | s__uncultured_bacterium | 0.5659 | 0.0014 | 1762.43 |
| s__Paenibacillus_barengoltzii | s__Thermobacillus_composti | 0.6261 | 3.00E-04 | 1517.84 |
| s__Paenibacillus_barengoltzii | s__Thermobifida_fusca | 0.705 | 0 | 1197.81 |
| s__Paenibacillus_barengoltzii | s__Thermobispora_bispora | 0.5876 | 8.00E-04 | 1674.39 |
| s__Paenibacillus_barengoltzii | s__uncultured_Acetanaerobacterium | 0.6577 | 1.00E-04 | 1389.94 |
| s__Paenibacillus_barengoltzii | s__uncultured_Alphaproteobacteria | 0.7373 | 0 | 1066.53 |
| s__Paenibacillus_barengoltzii | s__uncultured_Firmicutes | 0.5163 | 0.0041 | 1963.65 |
| s__Paenibacillus_barengoltzii | s__uncultured_Thermobacillus | 0.7036 | 0 | 1203.58 |
| s__rumen_bacterium | s__uncultured_compost | 0.5538 | 0.0018 | 1811.68 |
| s__rumen_bacterium | s__uncultured_Coriobacteriaceae | 0.5491 | 0.002 | 1830.81 |
| s__rumen_bacterium | s__uncultured_rumen | 0.5618 | 0.0015 | 1779.07 |
| s__Saccharopolyspora_rectivirgula | s__Symbiobacterium_thermophilum | 0.5647 | 0.0014 | 1767.4 |
| s__Saccharopolyspora_rectivirgula | s__Thermobacillus_composti | 0.5021 | 0.0055 | 2021.4 |
| s__Saccharopolyspora_rectivirgula | s__Thermobifida_fusca | 0.7007 | 0 | 1215.22 |
| s__Saccharopolyspora_rectivirgula | s__Thermobispora_bispora | 0.5332 | 0.0029 | 1895.14 |
| s__Symbiobacterium_thermophilum | s__Thermasporomyces_composti | 0.6357 | 2.00E-04 | 1479.1 |
| s__Symbiobacterium_thermophilum | s__Thermobacillus_composti | 0.733 | 0 | 1084.03 |
| s__Symbiobacterium_thermophilum | s__Thermobifida_fusca | 0.6671 | 1.00E-04 | 1351.41 |
| s__Symbiobacterium_thermophilum | s__Thermobispora_bispora | 0.7944 | 0 | 834.731 |
| s__Symbiobacterium_thermophilum | s__uncultured_Acetanaerobacterium | 0.5128 | 0.0044 | 1977.93 |
| s__Symbiobacterium_thermophilum | s__uncultured_compost | 0.7368 | 0 | 1068.7 |
| s__Symbiobacterium_thermophilum | s__uncultured_Dehalobacterium | 0.5933 | 7.00E-04 | 1651.22 |
| s__Symbiobacterium_thermophilum | s__uncultured_Thermobacillus | 0.7151 | 0 | 1156.66 |
| s__Thermasporomyces_composti | s__Thermobacillus_composti | 0.6676 | 1.00E-04 | 1349.59 |
| s__Thermasporomyces_composti | s__Thermobispora_bispora | 0.7038 | 0 | 1202.75 |
| s__Thermasporomyces_composti | s__uncultured_compost | 0.668 | 1.00E-04 | 1348.11 |
| s__Thermasporomyces_composti | s__uncultured_Dehalobacterium | 0.5274 | 0.0033 | 1918.65 |
| s__Thermasporomyces_composti | s__uncultured_Thermobacillus | 0.7499 | 0 | 1015.27 |
| s__Thermobacillus_composti | s__Thermobifida_fusca | 0.7911 | 0 | 847.993 |
| s__Thermobacillus_composti | s__Thermobispora_bispora | 0.8575 | 0 | 578.427 |
| s__Thermobacillus_composti | s__uncultured_Acetanaerobacterium | 0.5222 | 0.0037 | 1939.89 |
| s__Thermobacillus_composti | s__uncultured_Alphaproteobacteria | 0.5606 | 0.0016 | 1783.82 |
| s__Thermobacillus_composti | s__uncultured_compost | 0.705 | 0 | 1197.87 |
| s__Thermobacillus_composti | s__uncultured_Thermobacillus | 0.8474 | 0 | 619.658 |
| s__Thermobifida_fusca | s__Thermobispora_bispora | 0.7548 | 0 | 995.328 |
| s__Thermobifida_fusca | s__Thermomonospora_curvata | 0.5023 | 0.0055 | 2020.72 |
| s__Thermobifida_fusca | s__uncultured_Acetanaerobacterium | 0.6011 | 6.00E-04 | 1619.59 |
| s__Thermobifida_fusca | s__uncultured_Alphaproteobacteria | 0.6264 | 3.00E-04 | 1516.91 |
| s__Thermobifida_fusca | s__uncultured_archaeon | 0.5077 | 0.0049 | 1998.75 |
| s__Thermobifida_fusca | s__uncultured_compost | 0.5901 | 8.00E-04 | 1664.13 |
| s__Thermobifida_fusca | s__uncultured_Paludibacter | 0.6019 | 6.00E-04 | 1616.24 |
| s__Thermobifida_fusca | s__uncultured_Parabacteroides | 0.5891 | 8.00E-04 | 1668.42 |
| s__Thermobifida_fusca | s__uncultured_Thermobacillus | 0.7674 | 0 | 944.279 |
| s__Thermobispora_bispora | s__uncultured_Alphaproteobacteria | 0.5737 | 0.0011 | 1730.94 |
| s__Thermobispora_bispora | s__uncultured_archaeon | 0.5073 | 0.005 | 2000.23 |
| s__Thermobispora_bispora | s__uncultured_compost | 0.8193 | 0 | 733.685 |
| s__Thermobispora_bispora | s__uncultured_Thermobacillus | 0.8799 | 0 | 487.594 |
| s__uncultured_Acetanaerobacterium | s__uncultured_Paludibacter | 0.6043 | 5.00E-04 | 1606.55 |
| s__uncultured_Acetanaerobacterium | s__uncultured_Parabacteroides | 0.5129 | 0.0044 | 1977.61 |
| s__uncultured_Acetanaerobacterium | s__uncultured_Thermobacillus | 0.5316 | 0.003 | 1901.77 |
| s__uncultured_Acetanaerobacterium | s__uncultured_Veillonellaceae | 0.5497 | 0.002 | 1828.17 |
| s__uncultured_actinobacterium | s__uncultured_rumen | 0.5408 | 0.0025 | 1864.47 |
| s__uncultured_actinobacterium | s__unidentified_rumen | 0.635 | 2.00E-04 | 1481.93 |
| s__uncultured_Alphaproteobacteria | s__uncultured_Parabacteroides | 0.5244 | 0.0035 | 1931.13 |
| s__uncultured_Alphaproteobacteria | s__uncultured_Ruminobacter | 0.519 | 0.0039 | 1952.88 |
| s__uncultured_Alphaproteobacteria | s__uncultured_Thermobacillus | 0.559 | 0.0016 | 1790.34 |
| s__uncultured_Anaeroplasma | s__uncultured_bacterium | 0.5413 | 0.0024 | 1862.28 |
| s__uncultured_archaeon | s__uncultured_Firmicutes | 0.5401 | 0.0025 | 1867.33 |
| s__uncultured_archaeon | s__uncultured_Thermobacillus | 0.5734 | 0.0011 | 1732.17 |
| s__uncultured_bacterium | s__uncultured_Bacteroidales | 0.7177 | 0 | 1146.28 |
| s__uncultured_bacterium | s__uncultured_Bacteroides | 0.6452 | 2.00E-04 | 1440.35 |
| s__uncultured_bacterium | s__uncultured_Burkholderiales | 0.5654 | 0.0014 | 1764.59 |
| s__uncultured_Bacteroidales | s__uncultured_Bacteroides | 0.7221 | 0 | 1128.28 |
| s__uncultured_Bacteroidales | s__uncultured_Burkholderiales | 0.7162 | 0 | 1152.21 |
| s__uncultured_Bacteroidales | s__uncultured_Porphyromonadaceae | 0.5127 | 0.0045 | 1978.49 |
| s__uncultured_Bacteroides | s__uncultured_Burkholderiales | 0.6368 | 2.00E-04 | 1474.41 |
| s__uncultured_Bacteroides | s__uncultured_organism | 0.5841 | 9.00E-04 | 1688.42 |
| s__uncultured_Bacteroides | s__uncultured_prokaryote | 0.6807 | 0 | 1296.32 |
| s__uncultured_Burkholderiales | s__uncultured_Veillonellaceae | 0.5764 | 0.0011 | 1719.69 |
| s__uncultured_Clostridiales | s__uncultured_Ruminobacter | 0.5304 | 0.0031 | 1906.59 |
| s__uncultured_Clostridiales | s__uncultured_Veillonellaceae | 0.5979 | 6.00E-04 | 1632.51 |
| s__uncultured_compost | s__uncultured_Dehalobacterium | 0.5002 | 0.0057 | 2029.36 |
| s__uncultured_compost | s__uncultured_Thermobacillus | 0.7204 | 0 | 1135.11 |
| s__uncultured_Coriobacteriaceae | s__uncultured_rumen | 0.5053 | 0.0052 | 2008.57 |
| s__uncultured_Dehalobacterium | s__uncultured_Thermobacillus | 0.5191 | 0.0039 | 1952.26 |
| s__uncultured_Erysipelotrichaceae | s__uncultured_Porphyromonadaceae | 0.5531 | 0.0019 | 1814.45 |
| s__uncultured_Firmicutes | s__uncultured_Ruminobacter | 0.5306 | 0.0031 | 1905.77 |
| s__uncultured_Firmicutes | s__uncultured_Veillonellaceae | 0.5136 | 0.0044 | 1974.97 |
| s__uncultured_Lachnospiraceae | s__uncultured_Ruminococcaceae | 0.5294 | 0.0031 | 1910.47 |
| s__uncultured_organism | s__uncultured_prokaryote | 0.5644 | 0.0014 | 1768.44 |
| s__uncultured_organism | s__uncultured_Ruminococcaceae | 0.6438 | 2.00E-04 | 1446.36 |
| s__uncultured_Paludibacter | s__uncultured_Parabacteroides | 0.7764 | 0 | 907.9 |
| s__uncultured_Paludibacter | s__uncultured_Ruminobacter | 0.6857 | 0 | 1276.19 |
| s__uncultured_Paludibacter | s__uncultured_Veillonellaceae | 0.7382 | 0 | 1063.05 |
| s__uncultured_Paludibacter | s__unidentified_rumen | 0.5289 | 0.0032 | 1912.68 |
| s__uncultured_Parabacteroides | s__uncultured_Ruminobacter | 0.5861 | 8.00E-04 | 1680.46 |
| s__uncultured_Parabacteroides | s__uncultured_Veillonellaceae | 0.5563 | 0.0017 | 1801.31 |
| s__uncultured_prokaryote | s__uncultured_Ruminococcaceae | 0.7295 | 0 | 1098.27 |
| s__uncultured_prokaryote | s__unidentified | 0.5649 | 0.0014 | 1766.59 |
| s__uncultured_rumen | s__unidentified_rumen | 0.7226 | 0 | 1126.28 |
| s__uncultured_Ruminobacter | s__uncultured_Veillonellaceae | 0.7581 | 0 | 981.969 |
| s__uncultured_Ruminococcaceae | s__unidentified | 0.5762 | 0.0011 | 1720.56 |
